# Sulfated mannan helps diatoms domesticate their microbiome

**DOI:** 10.1101/2024.07.03.601839

**Authors:** J. Krull, C.J. Crawford, C. Sidhu, V. Solanki, M. Bligh, L. Rößler, R.K. Singh, G. Huang, C.S. Robb, H. Teeling, P.H. Seeberger, T. Schweder, J-H. Hehemann

**Affiliations:** BIOM, Faculty of Chemistry & Biology, University of Bremen, Germany; MARUM, University of Bremen; Max Planck Institute for Marine Microbiology, Bremen; Max Planck Institute of Colloids and Interfaces, Potsdam, Germany; Institute of Pharmacy, University of Greifswald, Germany; Institute of Marine Biotechnology, Greifswald

## Abstract

Diatoms are a keystone phylum in Earth’s ecosystems, specializing in oxygen production and carbohydrate fixation that fuels global food webs. Diatoms host a microbiome, but how they preferentially collect bacteria with complementary traits remains unknown. Here we show that diatoms exude a C6-sulfated α-1,3-mannan that serves as a selective carbon source for adapted bacteria. Its structure was resolved by NMR spectroscopy, chromatography, chemical synthesis, and enzymatic dissection. Biochemical, physiological, and structural analyses revealed that specialized *Bacteroidota* employ a four-enzyme pathway to metabolize this mannan. Metagenomic and transcriptomic data indicate that the mannan globally selects for bacteria carrying these enzymes and associated traits. Because the mannan provides only carbon, oxygen, sulfur, and hydrogen, bacteria must obtain other essential elements from alternative sources, reinforcing metabolic interdependence. We propose that diatoms use sulfated mannans to attract beneficial partners and exclude competitors, thereby engineering a microbiome that enhances their productivity and underpins carbon cycling.

**Significance statement:** Eukaryotes host microbial partners that shape their health, yet how they selectively assemble beneficial microbes remains unclear. Using diatom microalgae as a model, we show they exude a sulfated mannan that nourishes highly adapted bacteria tracking them across the global ocean. Our findings suggest that single-celled eukaryotes can “domesticate” prokaryotes—analogous to how humans have domesticated animals—albeit on a microscopic scale. Dominating much of Earth’s aquatic surface, diatoms drive ∼20% of global photosynthesis. We propose that sulfated mannan contributes to this success by helping diatoms shape microbial partnerships that underpin planetary energy balance and atmospheric chemistry.

## Main

Domestication (lat. *domesticus,* belonging to the house) is a process where an organism with sufficient access to energy can maintain an environment that hosts other species with complementary traits (1). The process involves reciprocal exchange of molecules and activities enabled through cognate genetic adaptations in the interacting species. Over time, sustained environmental control can lead to the emergence of symbiosis (2) In multicellular eukaryotes, such as animals and plants, slow reproduction rates delay the detection of genetic changes in offspring, making it challenging to detect molecular signatures of domestication (3). As a result, the underlying process of domestication remains largely unknown, especially at the molecular level.

To investigate this process, we considered single celled eukaryotes, diatoms and their associated prokaryotes. In the sunlight fueled surface ocean, diatoms can produce several hundred thousand cells within days in just a milliliter of seawater (1, 4). They photosynthesize organic molecules that sustain the carbon metabolism of associated prokaryotes (4, 5). Importantly, both partners can be cultivated, which facilitates identifying coevolving genes and the molecules that mediate interactions. Some functions resemble feed, fertilizers, herbicides and fences. For example, diatoms release carbohydrates used by bacteria as a carbon source (6, 7). They also release antibiotics (5, 8, 9) that suppress bacteria with incompatible traits, analogous to fences deterring predators or herbicides suppressing weeds. Fed, protected, and relieved of competition, the retained bacteria can concentrate nutrients (10), and return them, along with hormones and vitamins (8, 10, 11), to the auxotrophic algal host. Hence, diatoms and bacteria form a model system to study the process of domestication.

Carbohydrates, such as the abundant carbon energy molecule glucose (100 Gt C annual synthesis rate (18), can mediate both positive and negative interactions. In closed ring form glucose provides energy, but in the open ring form, with an exposed aldehyde group, glucose is toxic. This aldehyde group is one reason why insulin must activate muscle cells to remove excess glucose, as overexposure will cause host cells to die (19). Similar dual effects of glucose on microbes have been described (20)(21). Thus, the same molecule can serve as both nutrient and toxin, offering hosts a mechanism to reward some partners while excluding others. Such “functional flips” provide a molecular basis for domestication, challenging our positively biased view (22) of how complex communities assemble. Extracting this ancient insight from science, fiction (23), and philosophy (24), and applying it to organic molecules thought to benefit microbes, may help to uncover unknown rules that regulate biomes, including those central to the carbon cycle.

Diatoms exude substantial quantities of organic carbon in the form of complex carbohydrates (glycans) (25–27). These glycans, alongside many other molecules, likely shape the reproducible communities of bacteria attracted to or repulsed by diatoms. These include *Alphaproteobacteria* (e.g., *Candidatus* Pelagibacter ubique; SAR11), *Roseobacteraceae, Gammaproteobacteria (e.g. Vibrionaceae), Bacteroidota (e.g. Polaribacter* spp., *Formosa* spp.), *Cyanobacteria, Verrucomicrobiota,* and many other taxa (28–30). The chemical structures and mechanisms that govern glycan effects on bacteria remain difficult to resolve. A glycan chain with over 1000 monomers, about the length of a bacterial cell, can be highly anionic. Sulfate and carboxyl groups render these glycans extremely surface-active, promoting coagulation with cations, minerals, bacterial cells, proteins, and other molecules (27, 31, 32). The resulting gels impede separation and purification, hindering structural determination. Without defined structures, the ecological effects of glycans on bacteria, with or without the enzymes to metabolize, evade, or extract nutrients, remain obscure.

Here we resolved the structure of a sulfated mannan secreted by diatoms collectively responsible for 20% of global primary production (18). This figure rises to 70% in coastal upwelling zones and ∼50% in temperate subpolar regions during spring and summer (33). T The mannan selectively recruits bacteria that harbor a genetic island for mannan utilization, consistently associated with diatoms across the global ocean microbiome. The mannan may mediate association and disassociation between specific bacterial taxa, traits, functions. From a eukaryotic perspective, prokaryotes are rapidly growing, abundant, mobile and accessible sources of essential nutrients. Here we deduce how diatoms may “domesticate” and farm prokaryotes.

## Results

### Diatoms may cultivate adapted bacterial taxa by exudation of sulfated mannan

During the analysis of diatom bloom-associated metagenomes and bacterioplankton genomes, we noted the recurrence of bacterial taxa with a putative polysaccharide utilization locus (PUL) for mannan that consistently co-occurred with diatoms (**Fig. 1**). The PUL is found in marine *Bacteroidota* including *Formosa* spp., *Ochrovirga* spp., and *Polaribacter* spp., which are which are globally distributed and often highly abundant, like the diatoms they frequently associate with. Some of these species can account for ∼10% of the bacterioplankton cells surrounding diatoms (34–36). We reasoned that, among many organic molecules, a mannan may contribute to a stable microbiome (37–40). To uncover this hypothetical mannan, we worked together with the centric diatom *Conticribra weissflogii* (formerly *Thalassiosira weissflogii*), a diatom model species (41).

**Figure 1:**
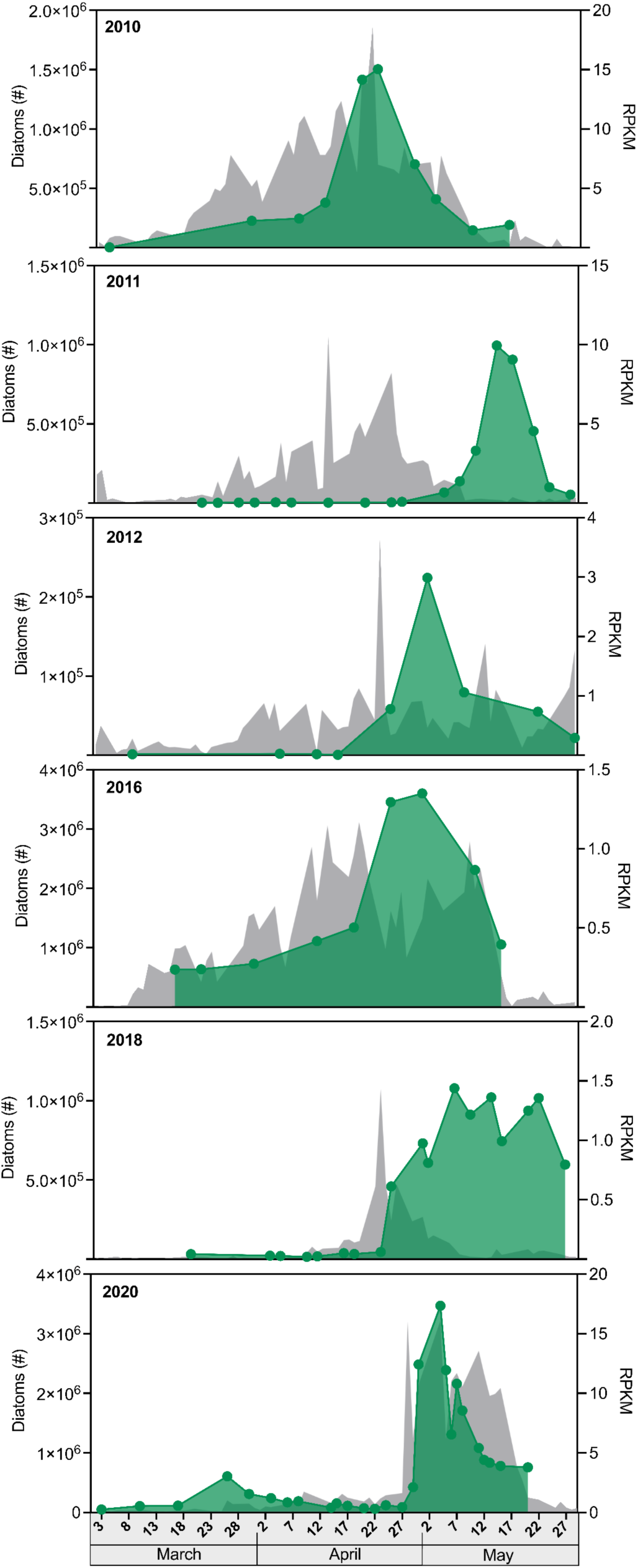
*Bacteroidota* with putative mannan utilization enzymes follow diatoms during algal blooms. Shown are data from six years from the long-term ecological research station (LTER) on Helgoland. The diatoms cells are followed by certain *Bacteroidota* taxa that have a polysaccharide utilization locus (PUL, **Fig. S2**) with putative enzyme and related genes that indicate utilization of a hypothetical, sulfated mannan secreted by the diatoms. The DNA sequence of the mannan PUL of *Polaribacter* sp. Hel1_33_49 was mapped against metagenomes collected previously from the LTER, Helgoland Roads. Grey shows diatom cell counts obtained by microscopy. Green shows the PUL abundance over years calculated in terms of Reads Per Kilobase per Million mapped reads (RPKM). For more information, see Materials and Methods.

### A C6-sulfated α-1,3-mannan secreted by a diatom

We previously used anionic exchange chromatography (AEX) to show that several diatom species, including *C. weissflogii*, secrete a fucose-containing sulfated polysaccharide (FCSP) of unknown structure (27). By refining AEX chromatography to achieve higher molecular resolution, we found that the material secreted by *C. weissflogii* comprises at least three anionic glycans: the FCSP, a glucuronomannan (42, 43) and a previously unknown homomannan. The FCSP and the glucuronomannan eluted at lower salt concentrations (0.5-1.2 M NaCl) and contained diverse monosaccharides, whereas, the homomannan eluted only at 2 M NaCl **(Fig. S1 A-C)**. Acid hydrolysis coupled with HPAEC-PAD (high-performance anion exchange chromatography with pulsed amperometric detection) revealed a composition of ∼95 mol % mannose. Additional ion chromatography revealed that the mannan contains an almost equimolar ratio of sulfate to mannose **(Fig. S1 D)**.

During chromatography, concentration, and dialysis, the mannan remained soluble in both seawater and deionized water. It reached solubility up to 20 mg mL^-1^, the highest tested, whereas other the algal polysaccharides (agar, κ-carrageenan, alginate and pectate) formed gels under seawater concentrations of calcium and magnesium ions (44). These insoluble glycans may target, or be targeted by, distinct bacterial populations (39), in contrast to the soluble mannan. From 160 L of diatom culture, we obtained between 15-20 mg of the mannan (∼100 µg L^-1^).

For configuration and connectivity, we used one- and two-dimensional NMR **(Fig. 2 A, B)**. Details are provided in the supplementary information. Briefly, the major component was α-1,3-D-mannopyranose with the C6-carbon de-shielded at 67.5 ppm, consistent with 6-*O*-sulfation and confirming the equimolar mannose-to-sulfate ratio **(Fig. 2B, Supplementary Text)**. For further verification, we synthesized a mannan oligosaccharide and used it as a reference for comparative NMR.

**Figure 2:**
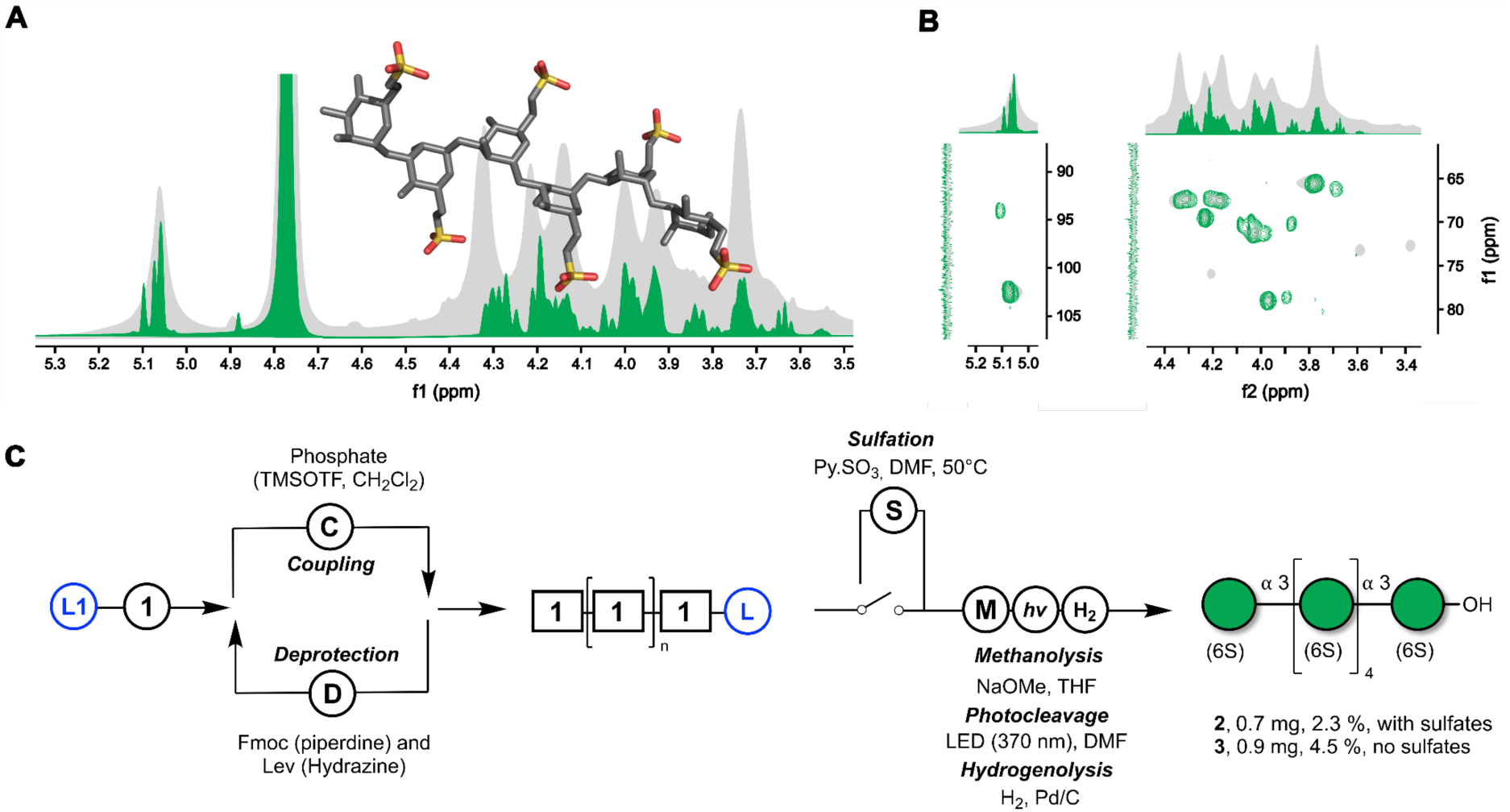
Diatoms synthesize a homomannan made of α-1,3-mannose-6-sulfate. In addition to chromatographic structure elucidation **(Fig. S1 B, D)** we used automated glycan assembly (AGA) to synthesize a mannan oligosaccharide of the NMR-hypothesized structure of the diatom mannan and compared the two spectra. (**A**) ^1^H-NMR overlay of an automated glycan assembly derived, synthetic α-1,3-mannose-6-sulfate hexaoligosaccharide (green) and the mannan purified from *Conticribra weissflogii* culture supernatant (gray). (**B**) Shown is an overlay of ^1^H-^13^C HSQC spectra from the synthetic mannan hexamer (green) and the purified, diatom mannan (gray). NMR spectra at 700 MHz were collected at 20 °C. HSQC-TOCSY and ^1^H-^13^C HMBC can be found in supplementary data. (**C**) Scheme of chemical synthesis of mannan oligosaccharides. L1: free reducing end solid-phase linker (45). 1: Dibutyl 2-*O*-benzoyl-4-*O*-benzyl-6-*O*-(9-fluorenylmethoxycarbonyl)-3-*O*-levulinoyl-1-phosphate-α-D-mannopyranoside.

We used automated glycan assembly to synthesize a reference mannan oligosaccharide. As a building block we prepared ethyl-2-*O*-benzoyl-4-*O*-benzyl-6-*O*-(9-fluorenylmethoxycarbonyl)-3-*O*-levulinoyl-1-thio-α-D-mannopyranoside, which was converted into a dibutyl phosphate donor (**1**) **(Fig. 2C, Supplementary Text).** This enabled the automated assembly of a hexa-α-1,3-mannose bearing one sulfate on each C6. Comparison of NMR spectra from natural and synthetic mannan showed deviations of <0.03 ppm in the proton and <0.1 ppm in the carbon dimension **(Fig. 2 A, B, Table S1)**, confirming the structure as a C6-sulfated α-1,3-mannan. **(Fig. S2)**.

### *Polaribacter* spp. can metabolize the mannan

We tested with *Polaribacter* strains - Hel1_33_49, _78 and _96 whether the PUL (46) enables mannan metabolism. These strains have 100% identical 16S RNA genes and 98.5% genome-wide average nucleotide identity (ANI) (47), providing three biological replicates. They have the PUL, indicating a similar ability to metabolize the mannan. Accordingly, when the mannan was provided as a limited carbon source, the three strains grew better **(Fig. S3-6, Table S2)**. As a negative control, we used *Polaribacter* sp. Hel1_88 (36, 46), which lacks the mannan-PUL (37). The Hel1_88 strain grew less on mannan in comparison to Hel1_33_49, _78 and _96. The mannan remained unconsumed in the culture of *Polaribacter* sp. Hel1_88 **(Fig. S4)**.

In 2020 we showed *Polaribacter* transcribe this PUL in situ (38), the North Sea. To confirm transcription and translation of enzymes capable of depolymerizing the mannan, we used cell lysates of *Polaribacter* sp. Hel1_33_78 for activity tests. Enzymes were extracted from bacterial cultures grown with or without the mannan **(Fig. S5,6)**. The enzymes in the lysates from cells grown with the mannan digested the mannan. These expression and activity assays suggest these enzymes of adapted *Polaribacter* spp. are for the metabolism of the mannan. The in situ enzyme expression indicates bacteria respond to and interact with the mannan released by diatoms into their environment.

### The bacterial enzymes can hydrolyze the mannan

To verify enzyme activity and specificity, we used recombinant gene expression, enzymology and structural biology **(Fig. 3)**. The PUL comprises 23 genes **(Fig. S2),** 21 of which encode putative enzymes. We cloned these genes from *Polaribacter* sp. Hel1_33_78 into vectors, transformed and expressed them in *Escherichia coli*, and conducted activity tests using purified mannan, synthesized mannan oligosaccharides, and para-nitrophenyl-labeled substrates (PNP-substrates). The products of these substrates digested with the enzymes were analyzed by fluorophore-assisted carbohydrate gel electrophoresis (FACE) and reducing sugar assays **(Table S3)**. To further substantiate the findings, we obtained homologous enzymes from another marine *Bacteroidota*, *Ochrovirga pacifica* S85, which was isolated from the Pacific Ocean (48) with 69.8% ANI with the *Polaribacter* sp. Hel1_33_78 genome.

**Figure 3:**
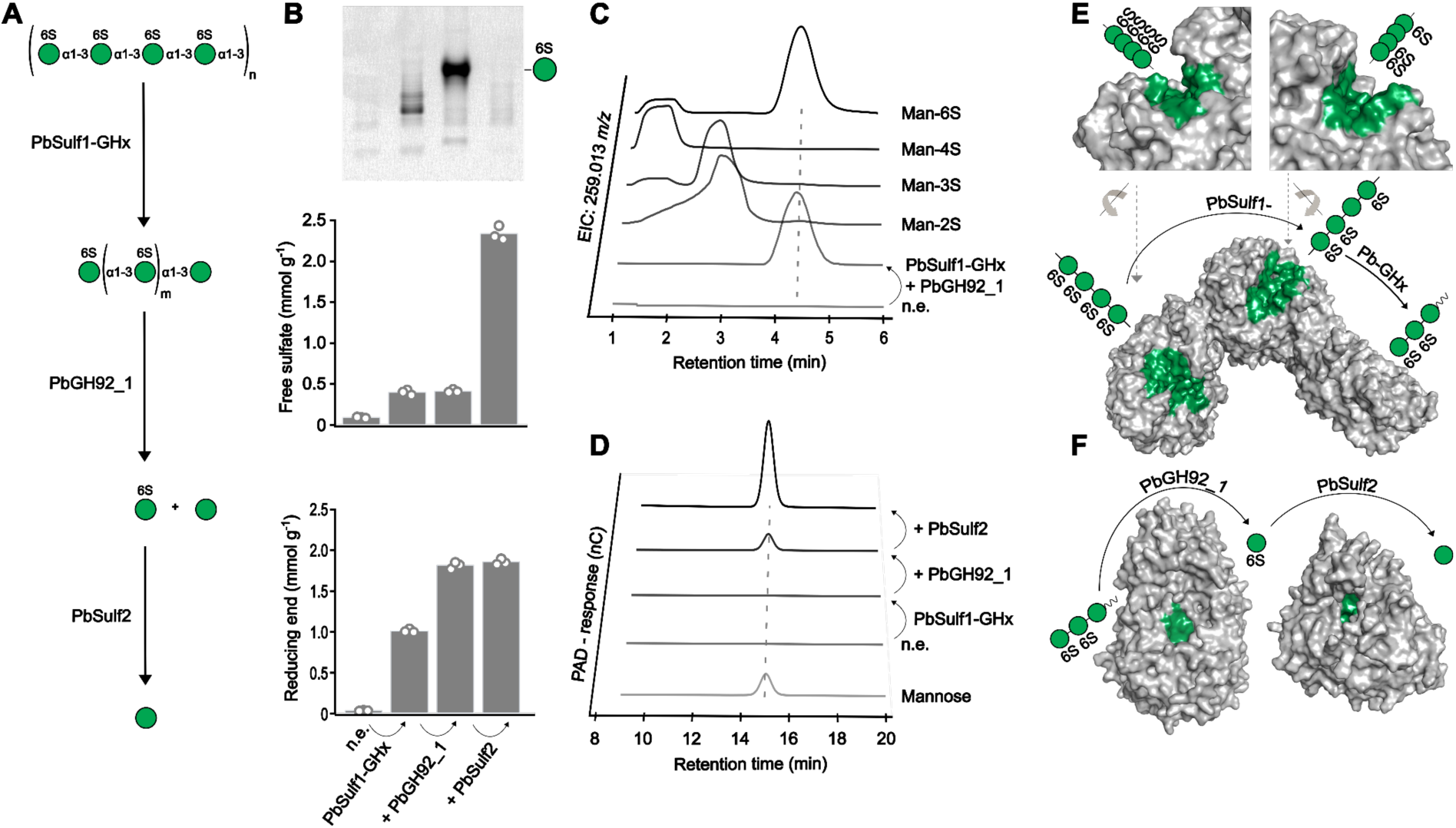
Four bacterial enzymes in three genes hydrolyze the sulfated α-mannan from diatoms. (**A**). Suggested degradative pathway for sulfated α-mannan. (**B**) Enzymatic digestions of the mannan via sequentially added PbSulf1-GHx, PbGH92_1 and PbSulf2. Top: Product separation by fluorophore-assisted carbohydrate electrophoresis using 2-aminoacridone as fluorophore. Middle: Free sulfate quantification. Bottom: Reducing end-assay. Bars represent mean of n=3 and circles display individual data points. n.e. = no enzyme. (**C**) Digestion of oligosaccharides produced by PbSulf1-GHx through PbGH92_1 were analyzed using LC-MS. Extracted ion chromatograms at 259.013 *m/z* of the digestion compared with synthesized or commercially purchased mannose-sulfate standards. Man-2S: Mannose-2-Sulfate; Man-3S: Mannose-3-Sulfate; Man-4S: Mannose-4-Sulfate; Man-6S: Mannose-6-Sulfate. (**D**) Mannose release after a sequential digestion of the mannan by PbSulf1-GHx, PbGH92_1 and PbSulf2 detected using HPAEC-PAD. (**E**-**F**) Structural model of enzymes involved in the degradative cascade were generated using AlphaFold2. Active site regions (green color) were identified using structural superimpositions with previously characterized enzymes. A simplified version of the degradative cascade is depicted according to the symbol nomenclature for glycans (51). (**E**) The PbSulf1-GHx tandem enzyme with the rotated active site region of the Sulfatase- and GHx-domain shown in top left and right, respectively. Grey arrows indicate the rotation axis of each domain towards the full-length protein (below). (**F**) Exo-acting PbGH92_1 and PbSulf2 have a pocket shape active site.

*O. pacifica* has a distantly related version of the PUL (49) with 68.8% ANI to the mannan PUL of *Polaribacter* sp. Hel1_33_78. We found that the four enzymes are present in both PULs (*Polaribacter*: PbSulf1-GHx, PbGH92_1, PbSulf2; *O. pacifica*: OpSulf1, OpGHx, OpGH92, OpSulf2). These enzymes, comprising one unknown family GHx, one GH92 and two sulfatases, hydrolyzed the mannan into mannose and sulfate **(Fig. 3A)**. The remaining 18 putative enzymes were cloned and tested but appeared inactive on the mannan **(Table S3)**. For one sulfatase (OpSulf1) the validity of biochemical experiments could be supported by X-ray crystallography analysis. For the other enzymes, which did not crystallize, we used *in silico* generated AlphaFold2 (50) structure models to falsify the biochemical data.

Mannan degradation is accomplished in four enzymatic steps. The first two steps are completed by the multimodular enzyme PbSulf1-GHx. The enzyme contains a T9SS domain for extracellular secretion. The sulfatase and GHx domain in PbSulf1-GHx work sequentially to catalyze steps one and two of the cascade. The sulfatase domain belongs to the S1 family, subfamily 51 (52). It is the first described activity of this subfamily and the first known sulfatase with this specificity. The GHx domain is distantly related to endo α-1,2 mannanases of the GH99 family and may therefore classify as a new family (53). Separating and visualizing fluorophore-labeled reaction products on FACE gels showed hydrolysis of the mannan by PbSulf1-GHx, producing a ladder-like pattern of oligosaccharides typical for endo-acting enzymes (54) **(Fig. 3B).** The tandem enzyme released sulfate and oligosaccharides as shown by ion chromatography and reducing sugar assays **(Fig. 3B)**. Liquid chromatography mass spectrometry (LC-MS) showed that the released oligosaccharides constituted sulfated hexose oligomers with degrees of polymerization between three and seven **(Fig. S7)**. The oligosaccharides were consistently missing one sulfate group, e.g. a mannose tetrasaccharide had only three sulfate groups, further indicating that both catalytic domains were active on the mannan.

We reproduced the results with enzymes from *O. pacifica,* which encodes the two catalytic domains as two separate genes. The enzymes OpSulf1 and OpGHx exhibit 53% and 36% amino acid identity to the respective PbSulf1-GHx domains. The separated enzymes showed that prior sulfatase activity is indeed necessary for GHx activity, suggesting that this enzyme requires a desulfated mannose for productive binding **(Fig. S8)**. The AlphaFold2 model of PbSulf1-GHx showed two appended catalytic domains with open, extended clefts **(Fig. 3E)** typical for endo-acting glycoside hydrolases (54) and sulfatases (55), corroborating the biochemical experiments. We obtained diffraction data from a crystal structure for the OpSulf1 at 1.49 Ȧ resolution (**Fig. S9, Table S4**). The crystal structure confirmed an extended substrate binding cleft compatible with the elongated mannan chain. The conserved catalytic residues present in the center may remove one sulfate from the center of the bound mannan chain, which then becomes accessible to the GHx **(Fig. S9B, C)**. In conclusion, we identified and characterized two new enzyme activities, one endo C6-sulfate α-1,3-mannanase forming a new CAZy family, and an endo C6-sulfate α-1,3-mannan sulfatase. These predicted outer membrane enzymes may produce oligosaccharides of suitable size and shape for import into the periplasm.

The third step in the mannan degradation cascade was catalyzed by the periplasmic PbGH92_1, a mannosidase that removed C6-sulfated mannose from the oligosaccharides. Periplasmic localization of PbGH92_1, which belongs to the GH92 family (56), was predicted by the Sec/SPI signal peptide (57). The GH92 family includes Ca^2+^-dependent exo-acting α-1,2/3/4/6 mannosidases, but an enzyme acting on, specific for sulfated mannan remains unknown. Hydrolysis of oligosaccharides into mannose-6-sulfate and a minor amount of mannose was shown by FACE, reducing sugar assays and HPAEC-PAD **(Fig. 3B-D)**. The mannose-6-sulfate product was verified using LC-MS **(Fig. 3C)**. For this analysis, we synthesized mannose-2/3/4-sulfate isomers **(Supplementary Text, Data S1, Fig. S10)**. The retention times revealed that only the mannose-6-sulfate aligned with the enzyme product peak and therefore confirmed this isomer as the product of the PbGH92_1 **(Fig. S11)**. The AlphaFold2 model showed a pocket topology of size and shape typical for an exo-acting GH **(Fig. 3F)** binding one monomer in the −1 subsite (54). Conclusively, PbGH92_1 is a 6-sulfate α-1,3-mannosidase, representing a new specificity within the GH92 family.

The fourth step was catalyzed by PbSulf2 that desulfates mannose-6-sulfate to yield mannose. A periplasmatic location was predicted by a Sec/SPII signal peptide. PbSulf2 is classified into sulfatase family S1, subfamily 11. Enzymes in this subfamily have been described to act on glucosamine-6-sulfate (58) and 6-sulfated porphyran oligosaccharides (59), but not on mannose-6-sulfate. PbSulf2 removes a sulfate group from mannose-6-sulfate as shown by appearance of the final product mannose in HPAEC-PAD measurements **(Fig. 3D)** and the disappearance of sulfated mannose in a complementary FACE experiment **(Fig. 3B).** The AlphaFold2 model aligned with the two previously described subfamily members (PDB ID Codes 5G2V / 7LJ2, rmsd: 0.613 Ȧ / 0.845 Ȧ) and showed a pocket topology typical for exo-acting sulfatases (55). PbSulf2 is a mannose-6-sulfate sulfatase, representing a new specificity within this enzyme family.

In summary, we elucidated a minimal enzymatic cascade consisting of four enzymes that achieve complete hydrolysis of the mannan into mannose and sulfate. The oligo- and monosaccharide structures produced by the enzymes confirmed the mannan structure solved with NMR.

### Biocatalytic quantification of mannan synthesis and degradation

To verify and quantify the synthesis of the mannan by diatom cells and its consumption by adapted *Polaribacter* spp. we used the here characterized enzymes to develop a quantitative assay. The assay involved a combination of chromatography and enzyme digest steps for structure-specific quantification. The enzymes specifically produce mannose that provides a much stronger signal than the undigested polysaccharide with different quantitative approaches, e.g., reducing sugar assay or HPAEC-PAD **(Fig. S12A)**. Chromatographic separation and enrichment of the mannan were crucial for enzyme digestion. Without this purification step, we found the enzymes remained inactive, potentially due to sulfate groups from FCSP or other sulfated glycans binding to and interfering with the enzymes in the mixture (60). After chromatographic isolation of the mannan, the biocatalytic assay detected up to 842 ± 62 µg L^-1^ in the diatom culture supernatant **(Fig. 4A).** The concentration was positively correlated with the number of diatom cells (*y* = 3.97 × 10^-3^ *x* - 2.94 × 10^-2^, *P* = 1 × 10^-23^, *R^2^* = 0.95). Each individual *C. weissflogii* cell produced on average a net amount of 3.9 fmol carbon in form of mannan **(Fig. 4B).** Substantially less mannan was present in diatom cells or particles **(Fig. S12B,C).** Notably, this new biocatalytic assay detects almost ten-fold higher concentrations (842 ± 62 µg L^-1^) compared to our initial assessment (∼100 µg L^-1^) based on the preparative mannan purification yield, highlighting the analytic value of enzyme-based assays.

**Figure 4:**
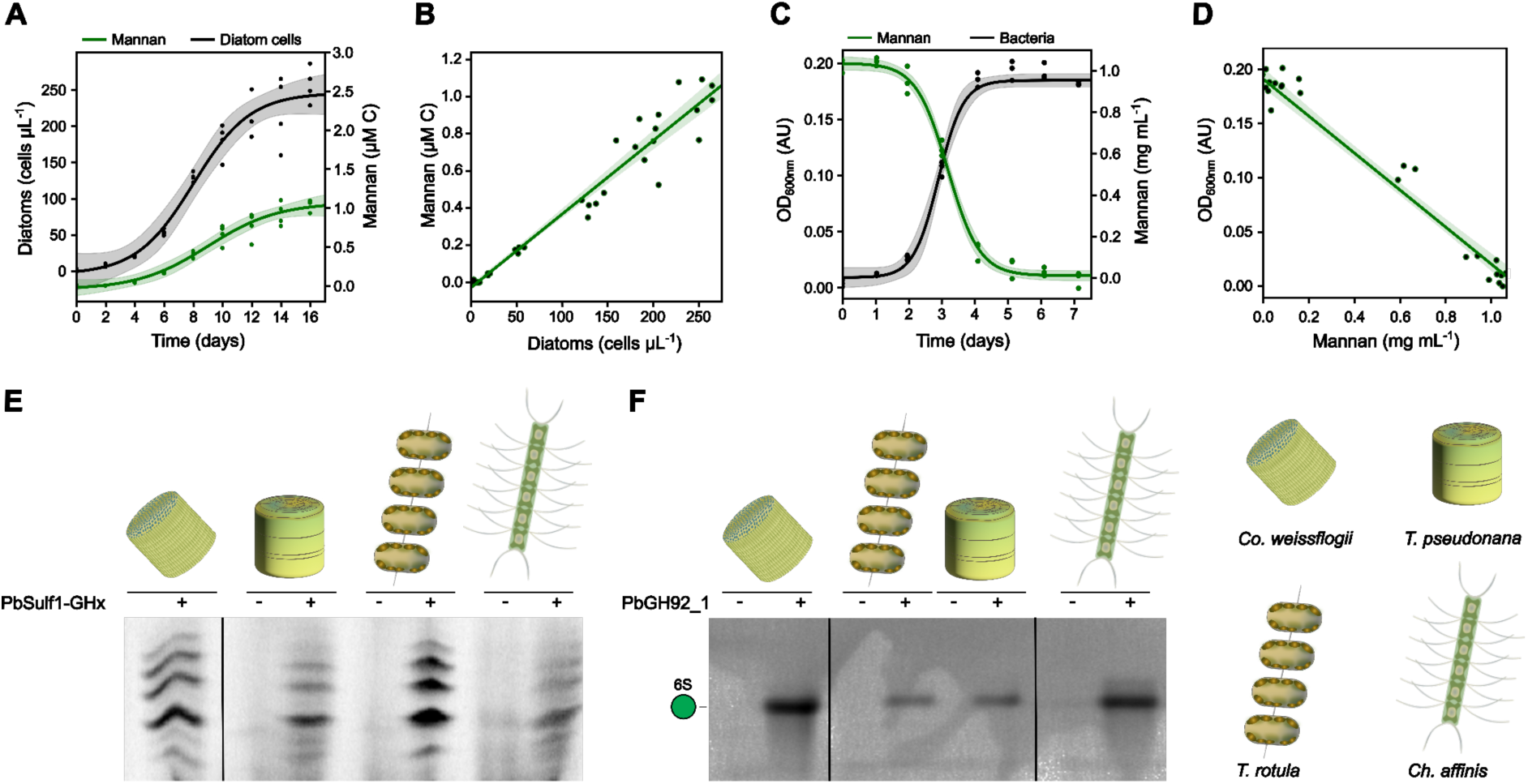
Synthesis of mannan by diatoms and consumption by bacteria quantified with the mannan degrading enzymes. (**A**). Growth analysis of *Conticribra weissflogii* cultures using cell counts (black) and enzymatic quantification of sulfated α-mannan (green) in the anionic fraction of the culture supernatant. Data points were modeled using a logistic growth function employing the non-linear least squares method for fitting. n=4. (**B**) Linear regression analysis of diatom cell counts vs. mannan C production. The net production of mannan per cell was 3.9 fmol C cell^-1^ with an R²=0.95. (**C**) *Polaribacter* sp. Hel1_33_49 grown with purified mannan for one week at 18 °C. The black line represents optical density measured at 600 nm. The green line shows the mannan consumption as detected via enzymatic quantification. Data points were modeled using a logistic growth function employing the non-linear least squares method for fitting. n=3. (**D**) Linear regression analysis of mannan concentration vs. optical density of the growth curve. (**E**-**F**) Mannan identification in anionic exudates purified from 0.5-5 M NaCl from AEX from *Conticribra weissflogii*, *Thalassiosira pseudonana*, *Thalassiosira rotula* and *Chaetoceros affinis*. Products were digested with PbSulf1-GHx (**E**) and PbGH92_1 (**F**) and analyzed via fluorophore-assisted carbohydrate electrophoresis using 8-aminonaphthalene-1,3,6-trisulfonic acid (**E**) or 2-aminoacridone (**F**) as fluorophore. Negative controls (-) contained heat inactivated enzyme.

Next, we conducted growth experiments with *Polaribacter* sp. Hel1_33_49 and applied the biocatalytic assay to quantify mannan consumption **(Fig. 4C)**. The bacteria grew to an OD_600_ of 0.2 when providing 0.1% (w/v) of the mannan as the limiting carbon source. Mannan concentration and bacterial abundance were negatively correlated (*y* = - 5.62 *x* + 1.09, *P* = 1 × 10^-18^, *R*^2^ = 0.96) **(Fig. 4D)**. Conclusively, under nutrient-replete conditions, the bacteria can remove the mannan within a few days.

To test whether other diatoms also release this type of mannan, we adapted the biocatalytic assay into a qualitative version that rapidly identifies the structure in different species. We tested *Thalassiosira pseudonana*, *Thalassiosira rotula* and *Chaetoceros affinis,* genera of global ecological and biogeochemical relevance (34). We used AEX to enrich the mannan from each culture supernatant, applied the enzymes and analyzed the products using FACE **(Fig. 4E, F)**. For each diatom the first two enzymes (PbSulf1-GHx) produced the same set of oligosaccharides that we identified above as oligomers of C6-sulfated α-1,3-mannose missing one sulfate group. PbGH92_1 released mannose-6-sulfate, showing that the same mannan structure is produced by different diatoms common around the globe.

### Mannan draws adapted bacteria towards diatoms around the globe

To test the hypothesis that the mannan selects for bacteria with these enzymes we quantified the mannan PUL within the *Tara* Oceans metaG dataset (61). As negative and positive controls for glycan structure-based selection of adapted microbiome members, we used bacterial PULs specific for known glycans from red, green and brown algal phyla. Preconditions for the PULs used as controls were that their target glycans are soluble and therefore accessible to planktonic bacteria, that the glycan structure has been verified by NMR and/or other methods, and that the structure and function of the PUL or of a closely related variant has been empirically verified with biochemical experiments. As negative control we used porphyran, a mixed linked α-, β-galactan that is also sulfated on C6, and the corresponding PUL from the flavobacterium *Zobellia galactanivorans* Dsij^T^ (62, 63). Compared to diatoms, porphyran-producing red macroalgae *Porphyra* spp., are sessile, require rocks or other hard substrates, grow along coasts. This range restriction makes the corresponding porphyran-PUL a suitable negative control.

As a positive control, we used laminarin (chrysolaminarin), a β-glucan that is also produced by diatoms, and the corresponding PUL from a metagenome-assembled genome (MAG C_MB344, accession: GCA_943788335) of a flavobacterial *Aurantivirga* species (38, 64, 65). Finally, we used ulvan, a sulfated rhamnogalacturonan produced by green macroalgal species including *Ulva* spp. *Ulva* spp. grows attached and planktonic, moves via currents between coastal and offshore regions (66), hence whether they, corresponding PUL can serve as negative or positive control was unclear. The corresponding PUL (67) is from the flavobacterium *Formosa agariphila* KMM 3901^T^ (68, 69).

Mannan PUL abundances ranged from 0.0025 to 0.41 RPKM in the 45 sampled sites including open and coastal ocean areas of the Atlantic, Pacific and Indian Ocean (**Fig. S13A**). Higher mannan PUL abundances were detected at upwelling sites (Benguela: 0.41 RPKM, Chile: 0.06 RPKM, Peru: 0.03 RPKM, California: 0.04 RPKM) where higher nutrient concentrations enhance diatom-dominated algal blooms (70, 71). The positive control, laminarin PUL showed comparable abundances between 0.006 and 2.27 RPKM **(Fig. S13C)**. The porphyran PUL showed the lowest RPKM values of the tested glycans and was absent in the open ocean **(Fig. S13E)**, as expected for this negative control. The ulvan PUL showed abundances between 0.002 and 0.08 RPKM but at different locations compared to the mannan and laminarin PULs **(Fig. S13D)**. The observed differences between the compared groups were significant across the dataset (Friedmann-Test: *P* = 6 × 10^-21^).

Metagenomics of bacterioplankton and eukaryotic 18S rRNA-based taxonomy revealed that the mannan PUL is associated with diatoms. To verify that diatoms were the main phytoplankton associated with the large RPKM values of PULs at the time of sampling for the *Tara* Ocean metaG dataset, we used previously reported total V9 read numbers as a proxy for diatoms and correlated those with PUL reads (71). This approach was previously verified by showing that V9 reads correlate with diatom algal cell counts (72). At the sites where microalgae were measured, diatoms were a major phytoplankton group (71). Linear regression analyses showed that the abundance of the mannan PUL correlated with diatom read numbers (*y* = 7.7 × 10^-5^ *x* + 6.2 × 10^-3^, *P* = 1.5 × 10^-3^, *R^2^* = 0.44) **(Fig. 5A)**. The laminarin PUL also significantly correlated with diatom abundance (*y* = 7.3 × 10^-4^ *x* + 1.7 × 10^-3^, *P* = 6.0 × 10^-4^, *R^2^* = 0.49) **(Fig. 5B)**. In contrast, we did not observe correlation between the ulvan PUL and diatom cell abundances (*P* = 0.35; R²= 0.05), which is consistent with the synthesis of this glycan by green algae instead of diatoms **(Fig. C)**.

**Figure 5:**
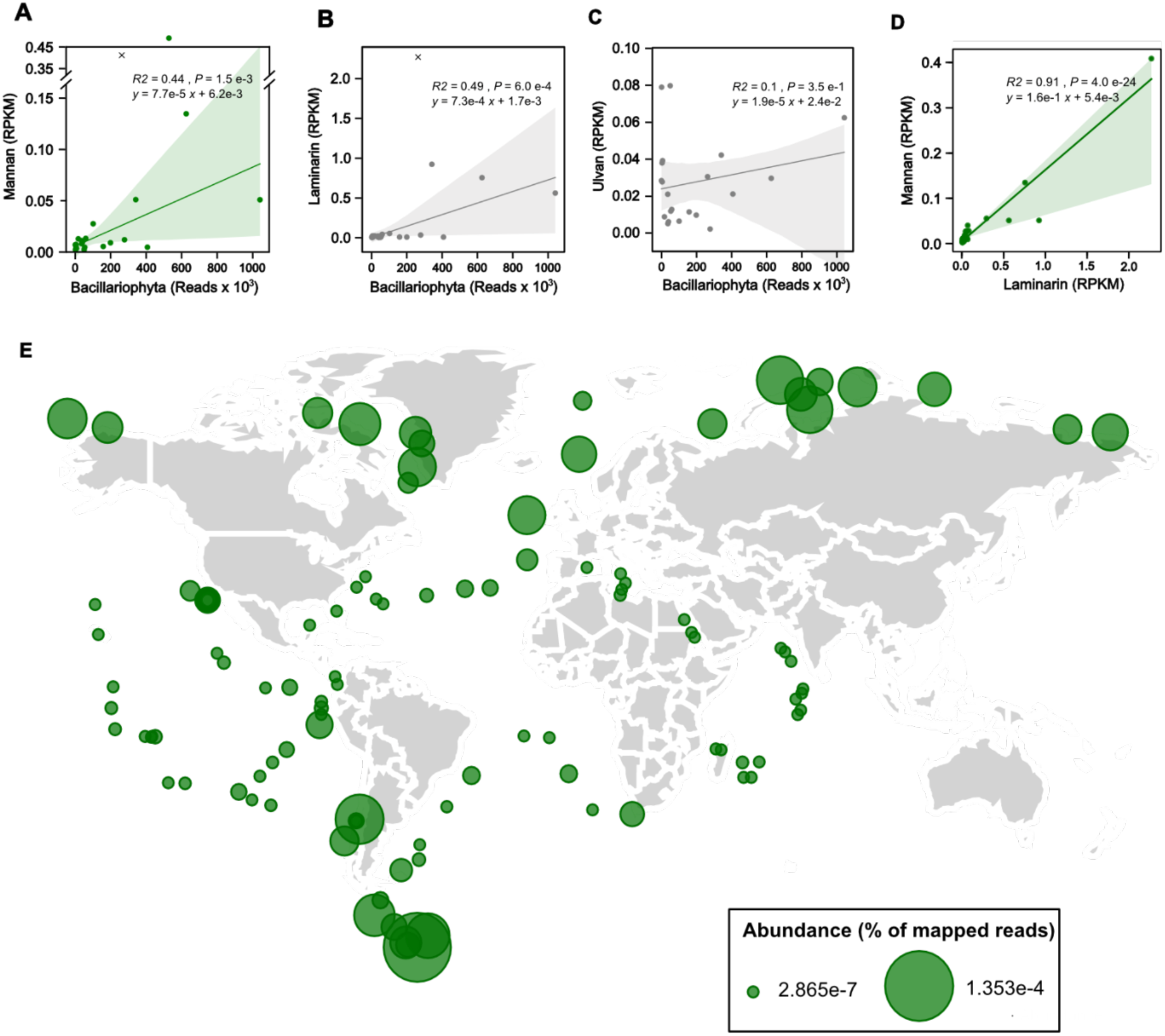
Bacteria with mannan degrading enzymes follow diatoms around the globe. Global abundance of the mannan utilization locus from *Polaribacter* sp. Hel1_33_49 from *Tara* Oceans database. The DNA sequence of the Mannan PUL was mapped against the raw metagenome reads of *Tara* Oceans surface water samples filtered at sizes ranging 0.2-3 µm across 45 different stations (61). Read abundances were calculated in terms of RPKM and normalized by length of the PUL and displayed as filled circles in SI. Corresponding plots for the laminarin PUL from *Aurantivirga* MAG C_MB344 (38), the ulvan PUL from *Formosa agariphila* KMM 3901^T^ (67) and the porphyran PUL from *Zobellia galactanivorans* Dsij^T^ (62) can be found in SI. (**A**) Raw read abundance of the clade of Bacillariophyta within the V9 rDNA metabarcoding dataset from *Tara* Oceans surface water samples filtered at sizes ranging 5-20 µm across 35 stations. Please note: No diatom data were available for the remaining stations. (**B**-**D**) Linear regression analyses of raw read abundance of Bacillariophyta in V9 rDNA metabarcoding dataset from *Tara* Oceans versus RPKM of different PULs across the same stations. Outliers where identified individually for each plot using the studentized residuals method with a threshold of three and excluded from the linear regression. Plots display the correlation analysis of Bacillariophyta vs. (**A**) mannan PUL, (**B**) laminarin PUL and (**C**) ulvan PUL. (**D**) Linear regression analyses of mannan PUL and Laminarin PUL RPKM within the *TARA* Oceans metagenomes. **(E)** The GH92 sequence from the *Polaribacter* cascade was searched against the Tara Oceans OM-RGCv2 database, which includes meta-transcriptomic (metaT) data, using BLASTp with an E-value cutoff of ≤ 1e-20 through the Ocean Gene Atlas v2 web platform. The resulting hits were used to calculate relative abundance, expressed as the percentage of mapped reads.

Diatoms are the only known microalgae that synthesize both, the mannan and the laminarin. Hence, we asked whether the PULs for those two glycans co-occurred, which would further support that diatoms are the source of these carbon-rich structures. We found a strong linear relationship between mannan and laminarin PULs (*y* = 0.158 *x* + 5.35 × 10^-3^, *P* = 4 × 10^-24^, *R²*= 0.91, **Fig. 5F**). In contrast, the PULs for mannan and ulvan did not correlate (*P* = 0.414; R²= 0.02). Finally, we queried for gene expression of the Polaribacter GH92 and found transcripts in the *Tara* Ocean metaT dataset (61). The presence of transcripts suggests associated bacteria degrade the mannan released by diatoms globally (**Fig. 5E)**. Spatially, temporally and phylogenetically separated micro- and macroalgae produce distinguished glycans. Conclusively, these diatom glycans, which select for specifically adapted bacteria, may contribute to the formation of a functional and predictable mosaic of host specific microbiomes in the sunlit ocean.

## Discussion

Here we show that diatoms secrete a system of anionic glycans that shape microbiomes in multiple ways. The structurally complex FCSP may protect against enzymatic degradation and invasion (73). The carboxyl groups of glucuronomannan may chelate rare trace metals (74). In contrast, the structurally simple mannan provides an easily accessible carbon source for adapted bacteria such as *Polaribacter* spp. which in return supply diatoms with missing nutrients, metabolites and vitamins (75). The characterized enzyme cascade can now be applied to quantify this mannan and assess its role in the carbon cycle (76). The global abundance of mannan degrading enzymes **(Fig. 5)** underscores its ecological importance.

The mannan can influence microbes both positively and negatively because sulfation renders glycans protein surface-active. Negatively charged sulfate groups bind to positively charged amino groups of proteins, and potentially interfere with their function (60). C6-sulfation extends the sulfate group farther from the sugar backbone, allowing it to reach positively charged residues even in protein pockets (62). Indeed, cation exchange resins designed to bind diverse protein surfaces with high capacity often use sulfated polysaccharides (77). C6-sulfation may be one of the reasons explaining why, of the three anionic diatom glycans, the mannan was the strongest binder of the surface-exposed amines **(Fig. S1 A-C).**

Prokaryotes depend on proteins to degrade, sense, bind, infect, import, move, replicate, transfer, and communicate (78). Many protein structures contain surface-exposed arginine, lysine or histidine residues that can be targeted by sulfate groups (77). This hypothesis aligns with Wheeler and colleagues who uncovered that mucilage glycans exuded by animal epithelia “tame” bacteria by binding to virulence proteins (79). By analogy, diatom-derived mannans may bind proteins across bacteria, eukaryotes, or even viruses, reducing their invasive potential.

During algal blooms, sulfated glycans secreted by diatoms, including the mannan **(Fig. S1 A-C)**, coagulate into aggregates that entrap bacterial cells, proteins, minerals and other molecules (27, 80–82). Bacteria can bind those glycans, or be bound by them. This interaction may especially capture cells lacking evasive mechanisms, promoting their incorporation into sinking particles. The outcome is a relative increase of some taxa and a decline of other in the sunlit ocean. As dominant algal taxa shift, their exuded sulfated glycans also change, selecting for new bacterial traits **(Fig. 5)** and driving host-specific microbiomes (83, 84).

Trade-offs may shape adaptive strategies used by bacteria to evade surface-active glycans. By digesting the mannan the *Polaribacter* spp. “domesticated” by diatoms obtain both carbon and energy, while also removing the interference caused by sulfation. In contrast, bacteria, lacking these enzymes minimize interaction as an alternative defense strategy. For instance SAR11 *Alphaproteobacteria*, among the most successful groups of bacterioplankters around diatoms (85), have a “non-stick” cell surface that bind fewer surface-active glycans (86). This reduced binding is thought to underlie their greater resistance to grazing (86), but it comes at the cost of being unable to access glycan-derived carbon. Similar trade-offs are seen in Roseobacteraceae (75), which focus on smaller organic molecules as carbon sources (87). By avoiding competition for glycans, they maintain a free-living, planktonic lifestyle that nonetheless supports a “domesticated” association with diatoms.

This “competition-dispersal” tradeoff has been described in *Vibrio cyclitrophicus,* where closely related strains show signs of recent sympatric speciation into two groups: strong vs. weak binders (88). Strains with surface pili, including the mannose sensitive hemagglutinin (MSH) type involved in host infection (89), bind and exploit surface glycans more effectively than the strains lacking MSH genes. However, non-piliated strains, though less efficient at glycan binding and exploitation, can avoid being irreversibly trapped by glycans (90). Surface-active, adhesive glycans (27) may therefore disproportionately bind bacteria that rely on MSH pili or other attachment proteins, constraining those most poised to intimately interact with eukaryotic hosts. Similar mechanisms operate in animals: epithelial cells secrete the heavily glycosylated immunoglobulin A (IgA) to constrain pathogens. IgA enchains and immobilizes bacteria in aggregates, facilitating their clearance from the intestine (91). Cells trapped by glycans or glycoproteins can be expelled or separated from microbiomes by multiple processes, including bowel movements (91), sinking (83), filtration (27) and consumption (86).

In summary, glycans can simultaneously support (92, 93) and suppress (79, 94) bacterial traits. Highly defensive types such as SAR11, with cell surfaces that minimize interaction, or *Polaribacter* spp., equipped with enzymes to deconstruct glycans, can avoid being trapped. In contrast, offensive bacteria such as Vibrio, rich in surface exposed proteins for attachment, degradation, or infection, are more easily trapped and may therefore remain rare (95, 96). Alternatively, they may be “tamed” by glycans, downregulating interactive surface proteins, while upregulating dispersal proteins such as flagella (79). This short-term adaptation, imposed by surface-active glycans, enhances dispersal, reduces competition (90), and protects the host with its domesticated partners. While we highlighted only a few cases, such interactions likely affect every cell to some extent. Through this dual capacity to nourish and constrain, glycans contribute to the emergence of beneficial microbiomes and productive hosts.

To conclude, with sufficient light, CO_2_, and nutrients, bacteria cannot escape the abundance of photosynthesis-derived surface-active glycans. Around diatoms, these glycans enforce a “domesticated” state through long-term genetic adaptation, or a non-virulent, tamed phase through short-term physiological shifts. Improved molecular understanding will help resolve uncertainties in the large-scale application of such glycans. Future work should focus on the mechanisms by which surface-active glycans protect the collective of domesticated prokaryotes and their eukaryotic hosts against competition, predation, and infection.

This process of “domestication” is important in today’s context. By fixing CO_2_ into simple glycans, diatoms sustain rapidly growing bacteria that in turn boost diatom productivity. In response, diatoms invest in more complex glycans that resist degradation, locking away carbon. Through these reciprocal interactions, mutualistic and antagonistic alike, CO_2_ is fixed into glycans in ways that reinforce climate stability. Thus, by secreting sulfated glycans that both nourish and constrain, diatoms domesticate their microbiomes, a microscopic strategy with planetary consequences.

## Materials and Methods

### Mannan PUL abundance in *TARA* Oceans dataset

Metagenome data of bacterioplankton during spring phytoplankton blooms was generated in previous studies (29, 38–40, 97). For the global analysis, we used metagenome data from the *TARA* Oceans Expedition (61). The surface water metagenomes (0.2 to 3 µm) from 45 different stations were included in the analysis. Mapping of PULs was done with Bowtie2 (98) in the SqueezeMeta v1.6.2 pipeline. PUL abundances as reads per kilobase per million mapped reads (RPKM) were calculated using the formula: (metagenome reads mapped to the PUL*10^6^)/(length of PUL in kbp × total reads in a metagenome). For correlation of PULs to diatom abundance, we extracted the read numbers from 25 surface water samples (5-20 µm) from the eukaryotic V9 metabarcoding data (71).

### Mannan purification

*C. weissflogii* was cultured non-axenic in NEPCC medium (99). Diatom growth was monitored by cell counting using a Neubauer counting chamber (Marienfeld, Königshofen, Germany). Cells were separated from culture the supernatant using a continuous flow centrifuge (Thermo Fisher Scientific, Walthan, MA, USA) set at 150 mL min^-1^ and 6800 x g. The culture supernatant was filtered using a peristaltic pump through Whatman® glass microfiber filters (Grade GF/F, 0.70 μm). 160 L of *C. weissflogii*, cell and debris free, culture supernatant were loaded onto a preparative scale, self-assembled AEX column. Elution occurred with appropriate NaCl concentrations. Samples were dialyzed against MilliQ water (Merck) and freeze-dried prior to monosaccharide composition and sulfate analysis.

### Monosaccharide composition and sulfate content analyses

Purified mannan was dissolved in MilliQ water at 1 mg mL^-1^ and hydrolyzed by 1 M HCl (100 °C, 24 h) in in pre-combusted glass vials. HCl was removed by vacuum or diluted 1:1000 in MilliQ water (Merck) prior to further analyses. Monosaccharide measurements occurred as described previously using a Dionex ICS-5000+ system (Thermo Fisher Scientific) (100) and for sulfate measurements we used a Metrohm 761 compact ion chromatograph (Metrohm, Herisau, Switzerland). All samples for monosaccharide and sulfate composition were prepared in triplicates.

### Cloning of enzyme genes into plasmids, recombinant expression and purification

Genes were cloned using the USER cloning or Gibson assembly methodologies (New England Biolabs, Ipswich, NE, USA) with genomic DNA from *Polaribacter* sp. Hel1_33_78. Primer pairs are provided in supplementary table S2. USER cloning constructs were cloned into the pet28A vector (101) without signal peptides (102). Clones were validated by DNA sequencing. Plasmids transformed into *E. coli* BL21 (DE3) were induced in 1 L lysogeny broth (LB) medium containing kanamycin and isopropyl-β-D-1-thiogalactopyranoside (IPTG). For sulfatase activation, the formylglycine generating enzyme plasmid (Addgene plasmid repository, Watertown, MA, USA) (103), which was co-transformed, was induced with L-arabinose and ampicillin added right before cooling to 12 °C. Protein purification was conducted using standard procedures (104).

### Enzyme activity and specificity assessment

Enzyme reactions were carried out in 10 mM MOPS pH 7.5 and 500 mM NaCl with 0.1% (w/v) of mannan and enzymes were added in excess (0.01-0.1 mg mL^-1^). 3.5% Sea Salts (Sigma Aldrich, Burlington, MA, USA) were also added. Enzyme reactions were monitored using fluorophore-assisted carbohydrate electrophoresis (FACE) with carbohydrates labeled with 2-aminoacridone as described in (105). Reducing-end measurements were carried out using PAHBAH as detection reagent as described previously (7). The release of sulfate and mannose from enzymatic reactions were measured using ion chromatography and HPAEC-PAD. Enzymatic assays were carried out as technical replicates in triplicates unless otherwise stated.

### Crystallization, data collection, structure solution, refinement and AlphaFold2

Crystallization was performed in two drop 96-well crystallization plates in sitting drop format using commercial crystallization screens. Diffraction data was collected on DESY beamline P11. Measured reflections intensities were indexed, integrated and scaled using the autoproc pipeline available at the beamline facility. OpSulf1 structure was solved using molecular replacement with the AlphaFold2 predicted model as reference in PHASER (106). For automatic model building we used BUCCANEER (107). Refinement was done in REFMAC5 (108) and COOT (109). Model and structure factors for OpSulf_1 were deposited in the Protein Data Bank (PDB) with accession 9FVT. Corresponding data-processing and refinement statistics are summarized in Table S4. The structural comparison of OpSulf with homologs were performed using the PyMOL v.2.3.2 (Schrödinger, New York, NY, USA). For enzymes that did not crystallize we used models generated with AlphaFold2 for structural analyzes (50).

### Liquid chromatography-mass spectrometry of mannose-sulfate and mannan digests

Standards and digests were diluted in 90% UHPLC-grade and 10% (v/v) MilliQ water (Merck) for LC-MS measurements. 5 µL were injected to an Accucore-150-amide HILIC column (Thermo Fisher Scientific) held at 60 °C on a Thermo Fisher Vanquish Horizon UHPLC coupled to a Thermo Fisher Q-Exactive Plus MS. Mobile phases A and B consisted of 10 mM ammonium formate at pH 5 and UHPLC-grade acetonitrile, respectively. A gradient started after 1 min at 90% B and decreased from 90% to 40% B over 40 min at a flow rate of 0.4 mL min^-1^. The column was equilibrated at 90% for 16 min at the end of each run. Mono- and oligosaccharides was detected in negative mode. The detailed MS conditions are provided in the SI material.

### Bacteria cultivation

Growth experiments with *Polaribacter* spp. Hel1_33_49/78/96 and Hel1_88 were carried out in HaHa-medium supplemented with 0.1 mg mL^-1^ yeast extract and 2 mg mL^-1^ mannan from either *C. weissflogii* or *C. affinis* (110). We used *Polaribacter* sp. Hel1_33_78 for in depth physiological experiments. Cells were harvested by centrifugation, the pellets chemically lysed. Lysed cells, enzymes were activity tested on *C. weissflogii* and *C. affinis* mannan. Digests occurred overnight using 3.5% Sea Salts (Sigma), 10 mM MOPS pH 7.5 and 0.5 mg mL^-1^ mannan and were analyzed using carbohydrate-polyacrylamide gel electrophoresis (C-PAGE). C-PAGE analysis was carried out as described in (111) with modifications described in the SI. *Polaribacter* sp. Hel1_33_49 was grown in 3.5% Sea Salts (Sigma) with supplements (110). The mannan was used as limiting carbon source at a concentration of 1 mg mL^-1^. Growth curves (absorbance 600 nm) were plotted with a logistic growth function using the non-linear least squares method. Substrate consumption was verified qualitatively using C-PAGE.

### Enzyme assisted quantification of the mannan

Cultures of *Conticribra weissflogii* were grown in NEPCC medium as described above. Subsamples of 200 mL of the culture were filtered to remove cells and debris. The mannan in the filtrate was purified using AEX. After desalting the mannan was digested PbSulf1-GHx and PbGH92_1. Undigested sample served as negative control. Detection occurred with the PAHBAH assay. Results were compared with a standard curve containing defined amounts of mannan. The recovery was determined 80 ± 5% (n=3) by comparison to untreated mannan **(fig S7C)**.

### Extraction of mannan from cellular, diatom biomass

After diatom cultivation the cell pellet was freeze-dried and sequentially extracted using MilliQ water (Merck), 0.4 M EDTA (pH 8) and 4 M NaOH with 0.1% NaBH4 (27). For 1 g of biomass 20 ml of sample were used. In each extraction step the sample was sonicated and centrifuged at 14.000 x g for 10 minutes. NaOH extracts were neutralized to pH 7.5 using HCl and diluted in MilliQ to < 500 mM NaCl. The mannan content in all extracts was determined as described above.

### Biocatalytic detection of mannan in multiple diatom species

The diatom strains *Thalassiosira pseudonana*, *Chaetoceros affinis* and *Thalassiosira rotula* were cultivated as described above. 20 L of culture supernatant were concentrated after filtration using GF/F. The filtrate was concentrated by ultrafiltration and then dialyzed to remove salts. The dialyzed concentrate was purified by anionic exchange chromatography. The purified mannan was again dialyzed and then lyophilized. The dialyzed and freeze-dried samples were used as substrates for enzymatic assays using PbSulf1-GHx_1 and PbGH92_1. Digested samples were analyzed with FACE (116).

### Nuclear magnetic resonance and chemical synthesis of oligosaccharides

All chemicals used were reagent grade and used as supplied unless otherwise noted. A detailed description of the structural elucidation and glycan synthesis processes is provided in the supplementary information. The automated syntheses were performed on a custom-built synthesizer (Potsdam-Golm, Germany)(112–115). More synthesis and analysis details are provided in the SI.

## Acknowledgements

We thank Manual Liebeke for access to LC-MS devices and advise. We also thank C. Eschen, P. Subramanya, P. Schoppmeier for large-scale cultivation of *Conticribra weissflogii* and K. Imhoff for advice and support during sulfate quantification and T. Trautmann, A. Bolte and K. Föll for technical support. We thank Hagen Buck-Wiese for help with the LC-MS analysis, S. Niggemeier for assistance in cloning and expression and A. Sichert for help with genome assembly. We thank André Scheffel for providing the *C. weissflogii* culture. We further thank M. Schultz-Johansen for valuable discussions and his substantial help with the sampling campaigns.

## Author contributions

J.K. and J-H.H. planned and analyzed the experiments with support by T.S.. J.K. and J-H.H. designed the artwork with help of M.B.. J.K. cloned and expressed the genes, conducted all activity assays, purified the glycans and cultured marine diatoms. P.H.S., C.J.C. and R.K.S. synthesized the oligosaccharide and mannose-sulfate standards and conducted all NMR experiments. V.S. provided structural models and refined the crystal structure. H.T., C.S., J.K. and J-H.H. analyzed the metagenomes. L.R. purified the proteins, set up crystallization experiments and edited the manuscript. M.B. conducted LC-MS experiments. J-H.H. and J.K. wrote the manuscript with input from all authors

## Data and materials availability

All data are available in the manuscript or the supplementary materials.

## List of Supplementary Materials

Supplementary Text

Figs. S1 to S31

Scheme S1

Table S1 to S6

Data S1 Chemical Synthesis

