## Supplementary Information for "Sulfated mannan helps diatoms domesticate their microbiome"

**Polyelectrolyte mannan from diatoms reshapes sunlit ocean microbiome**

**This file includes:**

- Supplementary Text
- Figs. S1 to S31
- Scheme S1
- Tables S1 to S6
- Chemical Synthesis

### Supplementary text

#### NMR analysis of *T. weissflogii* mannan

To establish the *T. weissflogii* mannan polysaccharide primary structure and establish connectivity, $^{1}H$ **(Fig. S12)**, $^{1}H$-$^{13}C$ HSQC **(Fig. S15-16** and **S19-20)**, $^{1}H$-$^{1}H$ COSY **(Fig. S23-24)**, $^{1}H$-$^{13}C$ HMBC **(Fig. S25-27)** and HSQC-TOCSY (SI Figure 21 and 22) at 700 MHz were collected at 20 °C. Monosaccharide composition and NMR analysis indicated the presence of two monosaccharides **(Fig. S15-16)**. The major component was an α-1,3-D-mannan with the majority of the C6 carbon de-shielded to 67.5ppm, suggesting 6-*O*-sulfation **(Fig. S15)**. The second less abundant monosaccharide appears to be β-D-xylose due to the characteristic splitting of the C-5 axial and equatorial protons **(Fig. S20)**. A single peak in the glycan region of the NMR at 5.46 and 99.47 ppm was not assigned to be part of the polysaccharides structure **(Fig.S4, 7)**. This single peak in $^{1}H$-$^{1}H$ COSY had no vicinal coupling to other protons in the spectra, additionally, it did appear to be a part of any spin system, with no cross-peaks appearing in the HSQC-TOCSY experiment **(Fig. S22)**. Inter-residue linkage connectivity was determined via a HMBC with a correlation peak between the possible anomeric xylose proton and a C6 carbon at 4.43 and 68.1 ppm **(Fig. S26)**. Suggesting, that xylose residues could exist as β-1,6 branches to the α-mannan backbone. A minor peak at 4.62 and 75.9 ppm was characteristic of 4-*O*-sulfation **(Fig. S24)**, and using $^{1}H$-$^{1}H$ COSY it was found to be connected to a non-sulfated 6-OH group **(Fig. S20)**. In summary, the major component of the *T. weissflogii* polysaccharide was hypothesized to be an α-1,3-mannan with a high degree of 6-*O*-sulfation. To verify this proposed structure, we synthesised defined standards of the structure, one sulfated and one non-sulfated α-1,3-mannan oligosaccharide.

#### Automated synthesis of sulfated mannan oligosaccharides

The automated synthesis of the sulfated α-1,3 mannan backbone required a single building block **(Scheme S1)**. Initially an ethyl 2-*O*-benzoyl-4-*O*-benzyl-3-*O*-(9-fluorenylmethoxycarbonyl)-6-*O*-levulinoyl-1-thio-α-D-mannopyranoside building block was tested. The choice of the 9-fluorenylmethoxycarbonyl (Fmoc) as a temporary protecting group at the 3-OH would allow for fast extension of the mannan backbone, while the 6-*O*-levulinoyl (Lev) ester would allow for chemoselective 6-OH sulfation. However, automated glycan assembly (AGA) with this building block led to unsuccessful syntheses with deletion sequences and *m/z* that could not be accurately assigned to any particular structure. These altered *m/z* can be observed when target oligosaccharides contain Lev esters (*113*). These altered *m/z* are hypothesised to originate from unwanted side-reactions of the Lev ester. To circumvent this issue, we swapped the positions of the 6-OH and 3-OH temporary protecting groups, so that the Lev esters would be now employed for backbone assembly and Fmoc groups were used to mask hydroxyl groups destined for sulfation.

This ethyl 2-*O*-benzoyl-4-*O*-benzyl-6-*O*-(9-fluorenylmethoxycarbonyl)-3-*O*-levulinoyl-1-thio-α-D-mannopyranoside (**4**) building block was allowed AGA up to 3-mers **(Fig.**

**S29)**. However, larger oligosaccharide sequences resulted in deletion sequences as analyzed by high-pressure liquid chromatography (HPLC) and matrix-assisted laser ionization time-of-flight (MALDI-TOF) mass spectrometry (MS). Even double glycosylation cycles with thioglycoside building block **4** did not improve coupling efficiency. The low nucleophilicity of the 3-OH mannose acceptor or non-ideal temperature selection of the glycosylation module could be the origin of this poor efficiency (*114, 115*). The thioglycoside donor **4** was therefore converted into a dibutyl phosphate donor **1** **(*116*)**(Plante 1999). The dibutyl phosphate block allowed the automated assembly of the desired target hexasaccharides but required the use of double cycles after the 3-mer stage **(Fig. S30)**. On resin sulfation was performed using a sulfur trioxide pyridine complex in a sealed microwave vial ($SO_3$.Py, 10eq. per OH, DMF, 50 °C, 16 h) with the sulfation stage tracked by Q-TOF MS analysis of microcleavage samples (*117*). On resin methanolysis (5% solution of 0.5 M NaOMe in THF) removed the remaining *O*-ester protecting groups with incubation times of 72 hours required for the sulfated α-mannan, and 16 hours for the non-sulfated mannan **(Fig. S31)**. The resin bound glycans were released from the solid-support using a LED lamp (385 nm, DMF, 16 h) with the crude photocleavage products subjected to palladium-catalyzed hydrogenolysis. Purification by reverse phase HPLC produced sulfated oligosaccharide **2** (0.7 mg) and non-sulfated oligosaccharide **3** (0.9 mg), in yields of 2.3% and 4.5% respectively.

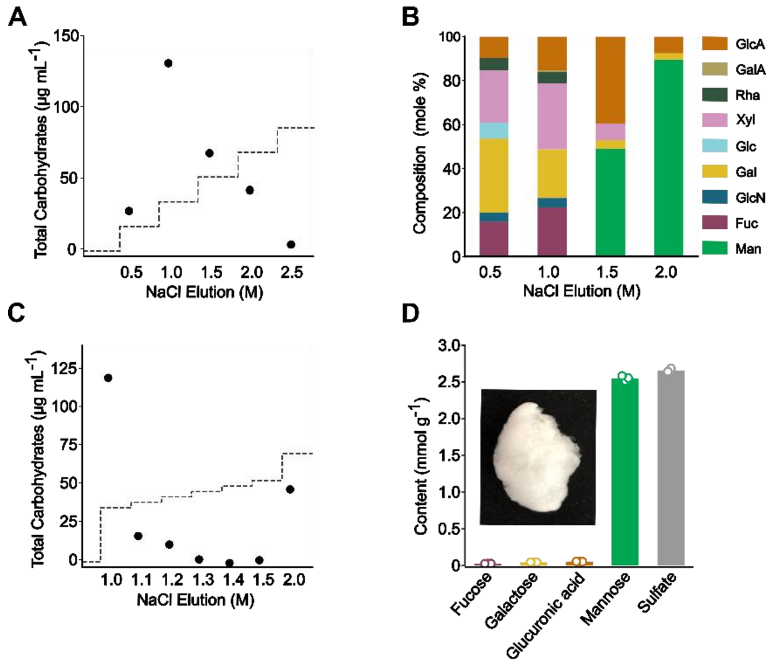


**Fig. S1.**

*Thalassiosira weissflogii* exudes mannan. (**A**,**C**) Anion exchange chromatography of culture supernatant of *Thalassiosira weissflogii* with stepwise increase of NaCl concentration. Total carbohydrate signals of each fraction are displayed as black dots, and stepwise NaCl elution profiles as dashed lines. (**A**) Step size of 0.5 M NaCl (**B**) Relative monosaccharide composition after acid hydrolysis of eluted fractions from analytical replicates (n=3) (**C**) Optimization of purification protocol with 0.1 M NaCl step size. A final window of 1.35-2.5 M NaCl was chosen for purification (**D**) Absolute monosaccharide and sulfate amount after acid hydrolysis of the optimized anion exchange protocol with elution steps of 1.35-2.5 M NaCl. The inset shows a photograph of the freeze-dried mannan.

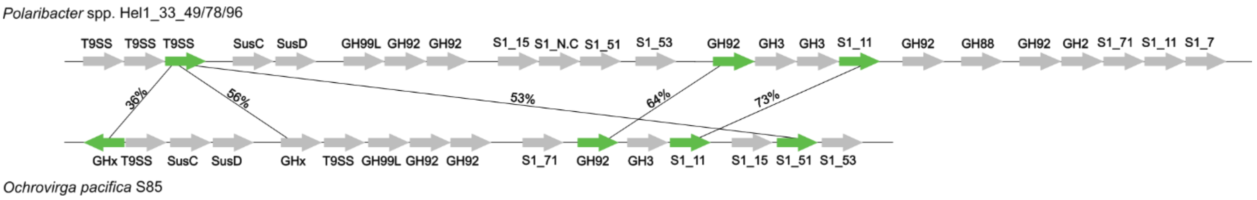


**Fig. S2.**

PUL structures in *Polaribacter* sp. Hel1_33_49/78/96 and *Ochrovirga pacifica* S85. The genes displayed in green are subject of this study.

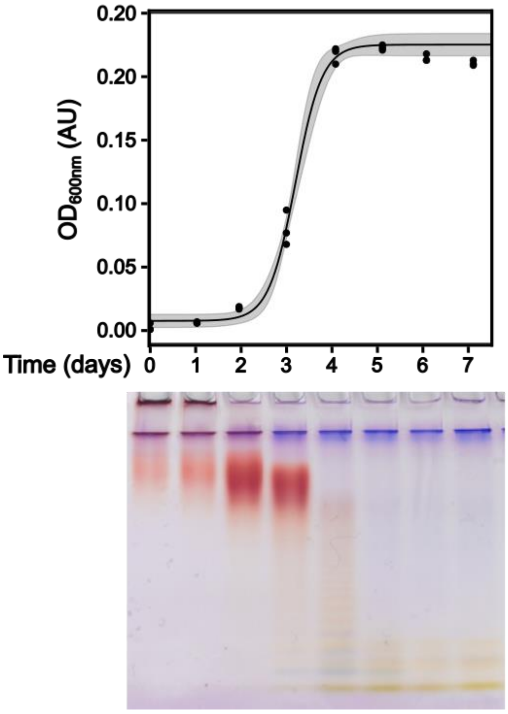


**Fig. S3.**

*Polaribacter* sp. Hel1_33_49 grown on mannan in 3.5% Sea Salt medium. Top: Growth curve as assessed by optical density measurements at 600 nm. Bottom: Carbohydrate-polyacrylamide gel electrophoresis of the culture supernatant during growth stained with 0.02% Stains-all.

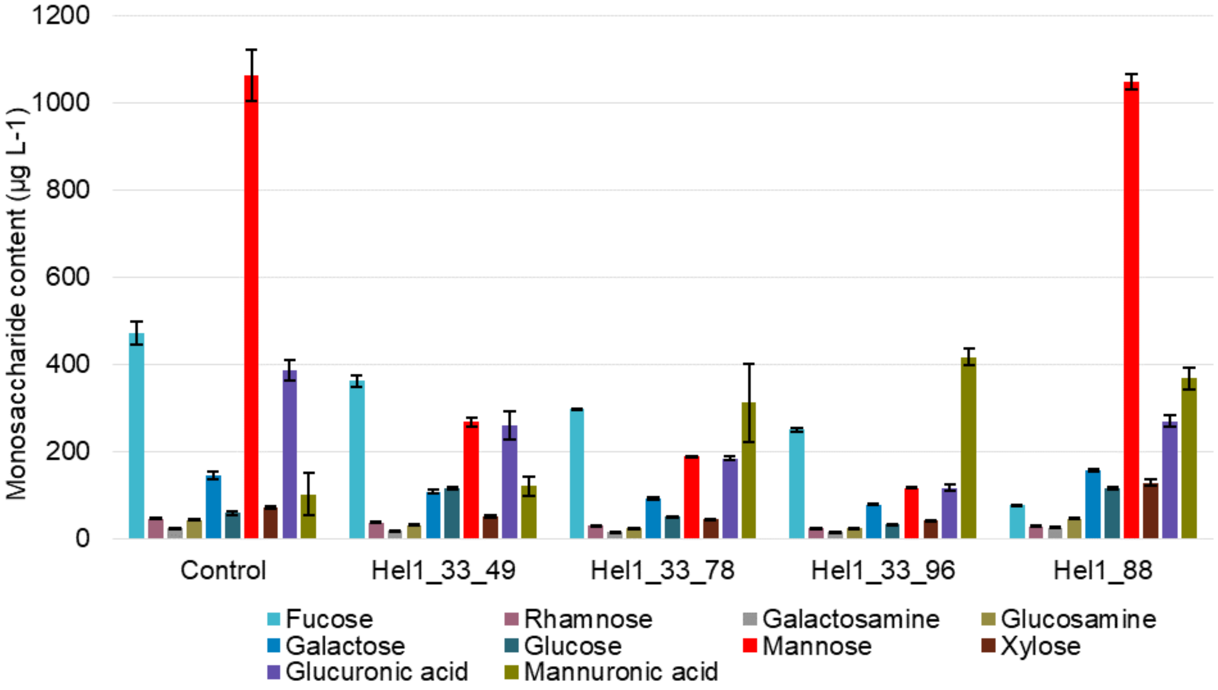


**Fig. S4.**

Growth of *Polaribacter* strains Hel1_33_49, _78, _96 and Hel1_88 on mannan from *Chaetoceros affinis* (n=3).

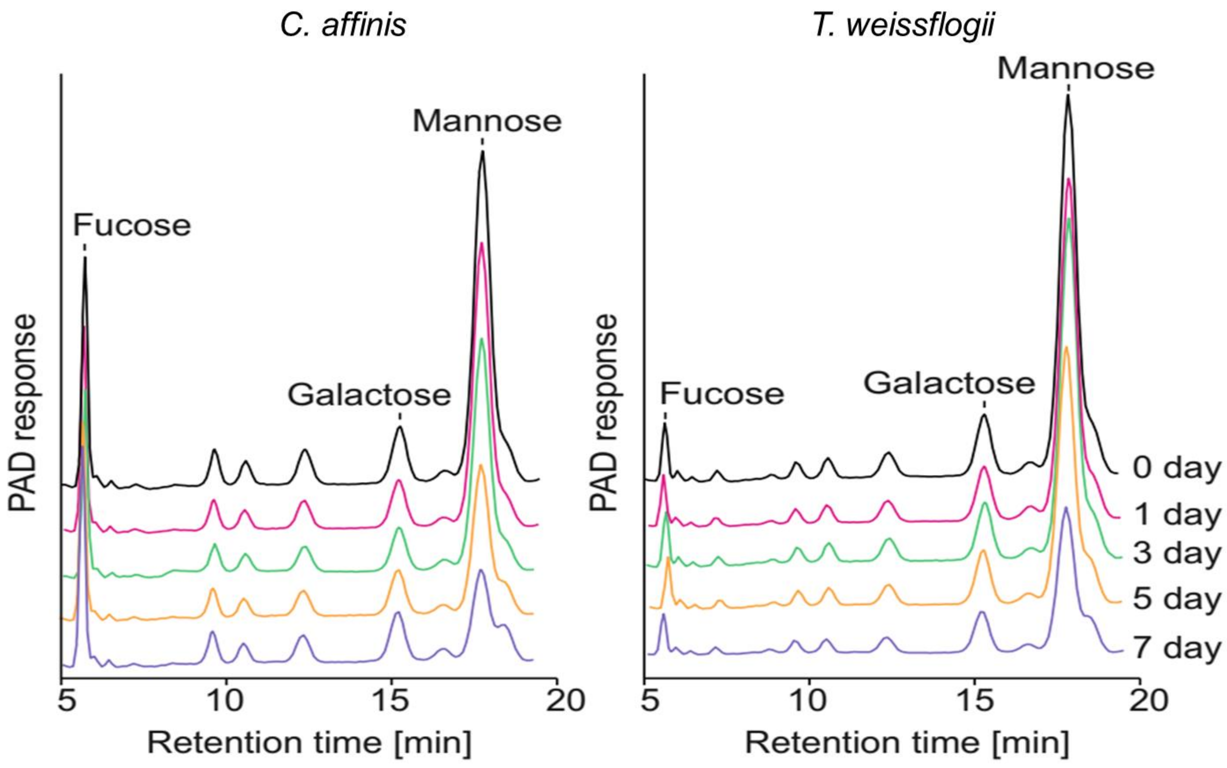


**Fig. S5.**

Growth of *Polaribacter* sp. Hel1_33_78 on mannan from *Chaetoceros affinis* and *Thalassiosira weissflogii.* Supernatants were sampled during growth and the decrease of monosaccharides after acid hydrolysis was determined using HPAEC-PAD.

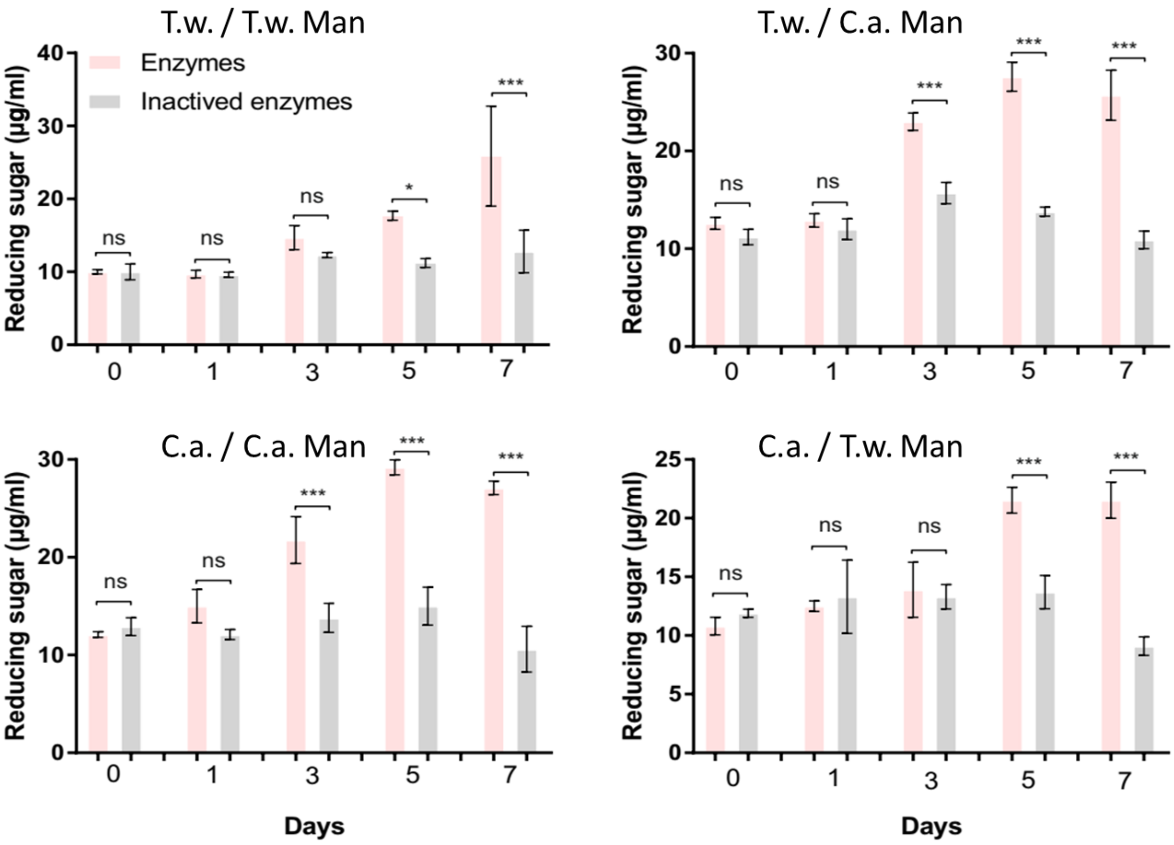


**Fig. S6.**

Growth of *Polaribacter* sp. Hel1_33_78 mannan from *Chaetoceros affinis* and *Thalassiosira weissflogii.* Cell lysates were produced after different induction times, incubated with both substrates and analyzed using a reducing sugar assay (n=3). Plot titles indicate “origin of assay substrate / origin of growth substrate.”

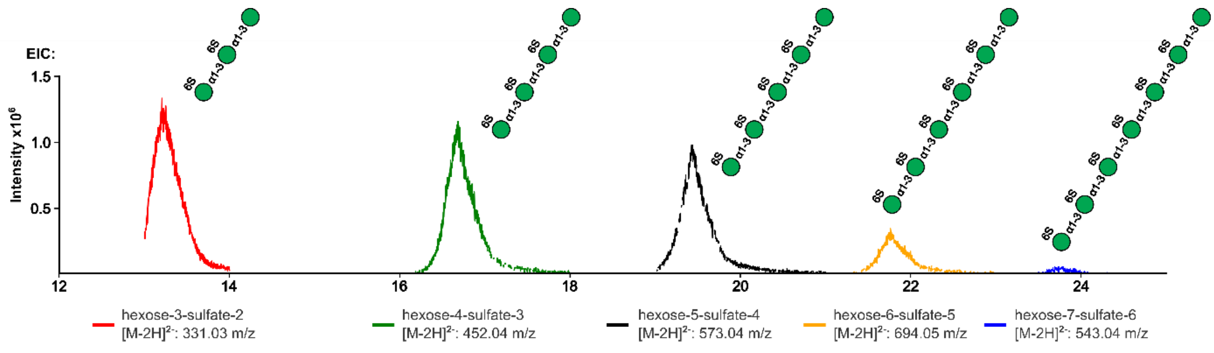


**Fig. S7.**

Mass-spectrometry shows oligosaccharide products of PbSulf1-GHx**.** Digestion of the mannan using PbSulf1-GHx analysed by Liquid Chromatography-Mass Spectrometry (LC-MS). Data show the extracted ion chromatograms of a subset of *m/z*-values corresponding to oligosaccharides found.

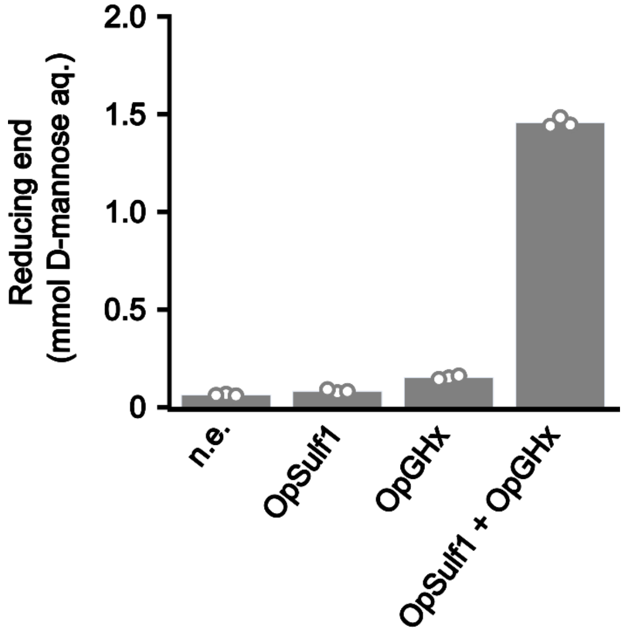


**Fig. S8.**

Initial mannan breakdown with two endo-acting enzymes. PbSulf1-GHx homologues OpSulf1 and OpGHx are both required for mannan oligomerization. Reducing-end assay of enzymes from *O. pacifica* (n=3).

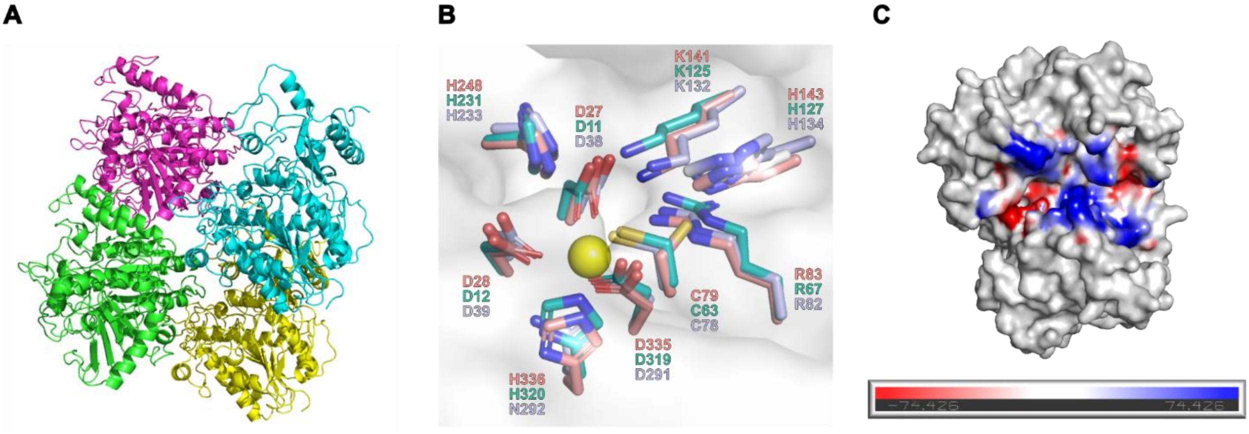


**Fig. S9.**

OpSulf1 crystal structure**.** (**A**) Cartoon representation of OpSulf1 with four molecules in the asymmetric unit. (**B**) Active site catalytic and binding residues of OpSulf1 (salmon) overlayed with PbSulf1-GHx (turquoise) and carrageenan sulfatase from *Pseudoalteromonas* (light blue, PDB-ID: 6B0K, rmsd: 1.317 Å). (**C**) Electrostatic surface charge representation of the active site region.

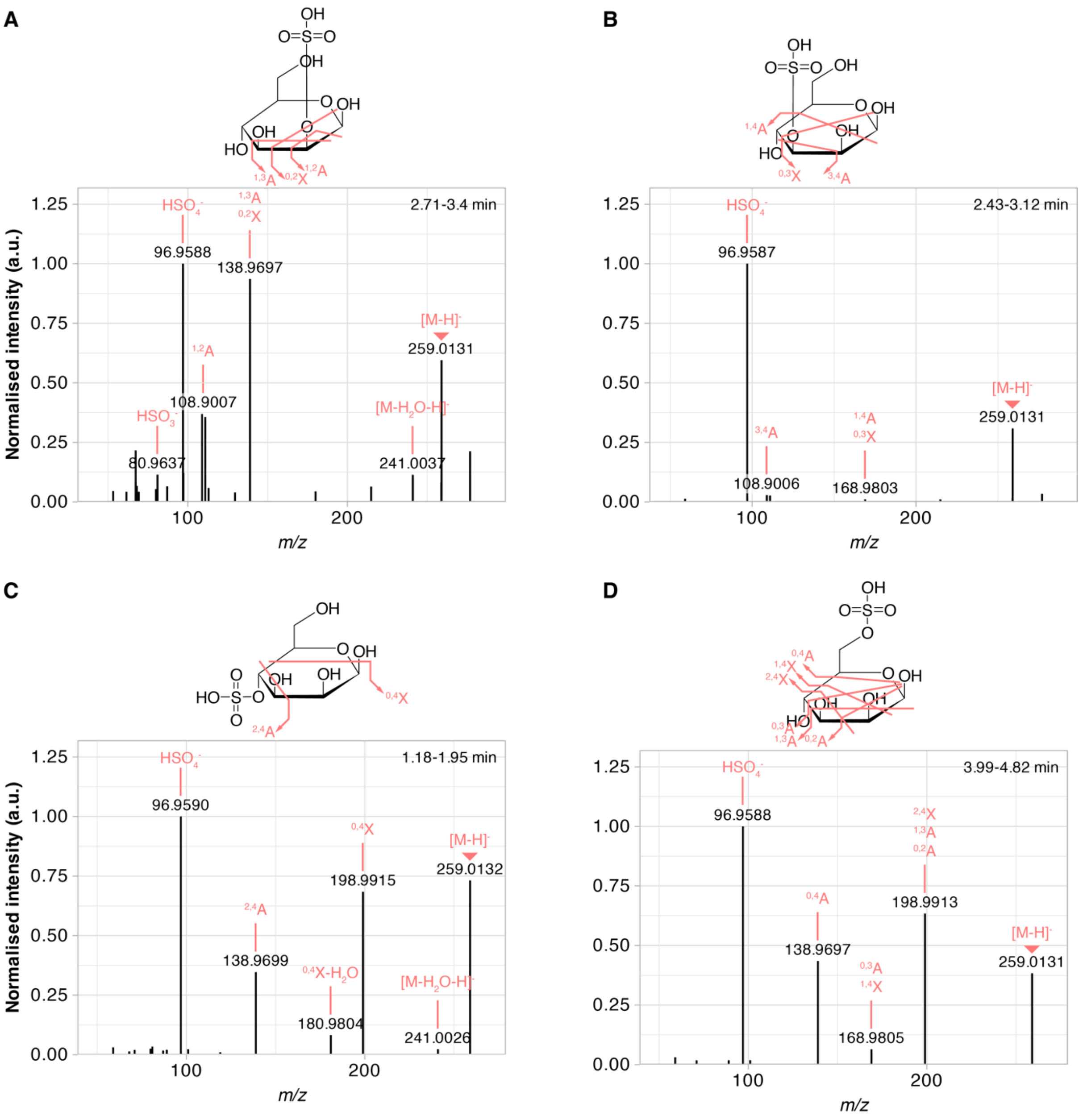


**Fig. S10.**

Fragmentation patterns discriminate different positions of sulfate on mannose. Averaged and normalized fragmentation spectra from LC-MS/MS experiments with mannose-2-sulfate (**A**), mannose-3-sulfate (**B**), mannose-4-sulfate (**C**) and mannose-6-sulfate (**D**). Structures show observed cleavages, with annotations according to the nomenclature of (*118*).

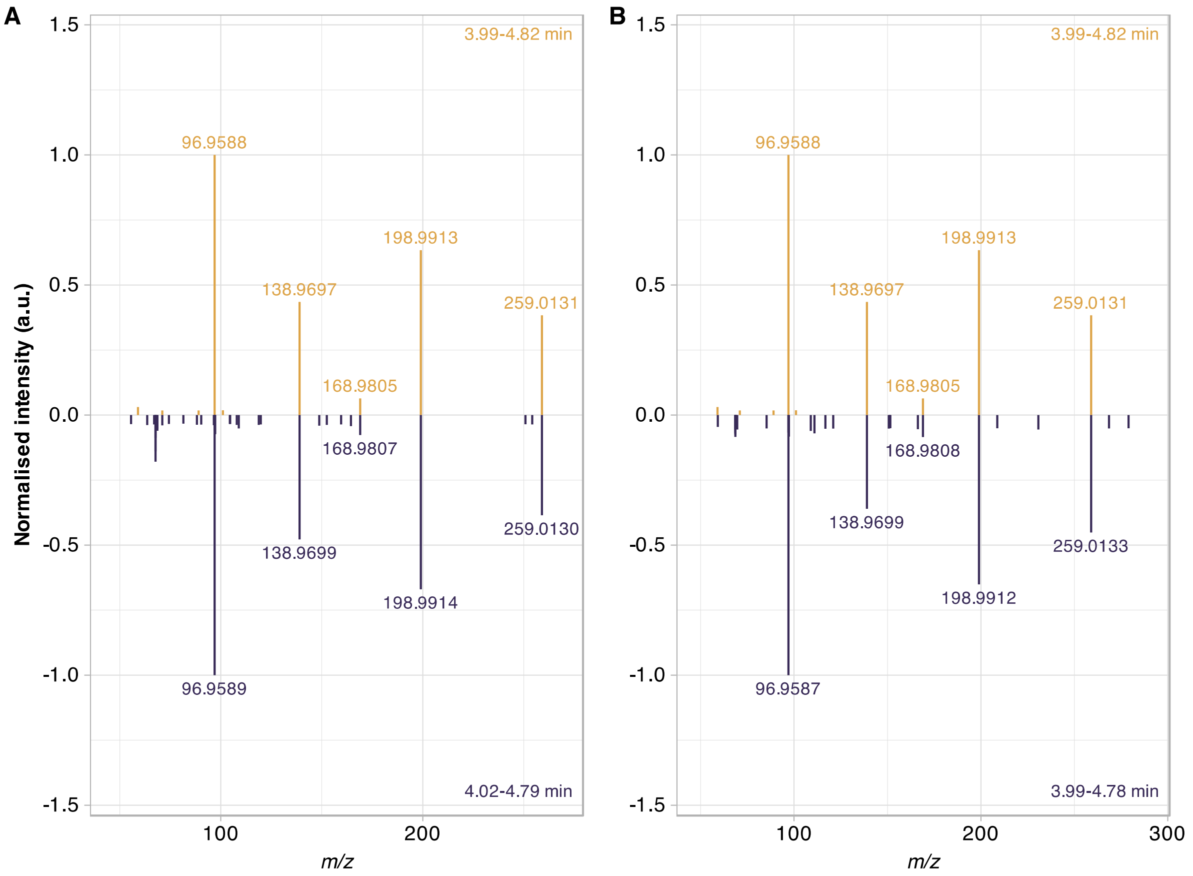


**Fig. S11.**
Comparison of fragmentation patterns confirms that digestion with PbSulf1-GHx and GH92_1 yields mannose-6-sulfate. MS/MS spectra from fragmentation of mannose-6-sulfate (upper panels, yellow) and duplicate digests (lower panels, purple). Retention times of spectra used for averaging are in the corners.

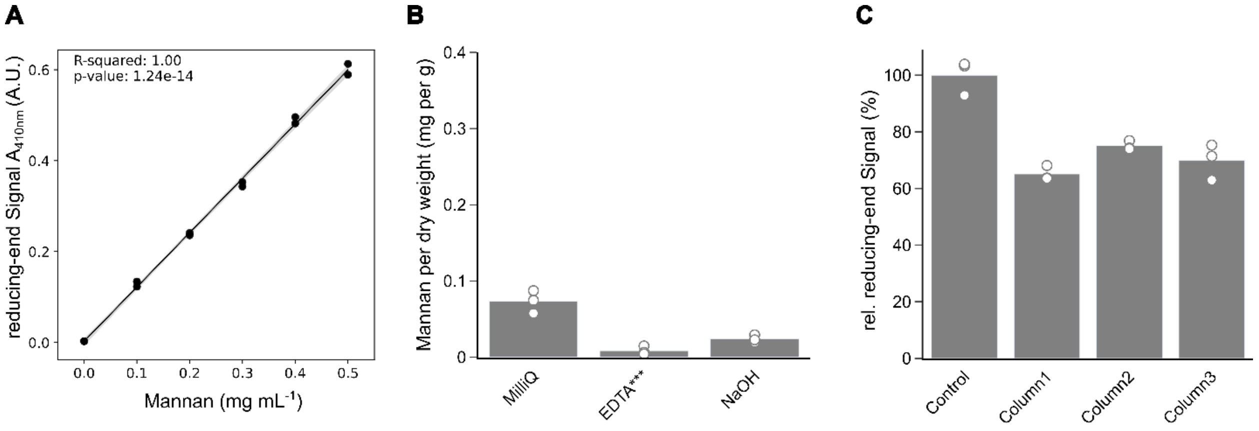


**Fig. S12.**

Enzymatic quantification of mannan. (**A**) Product formation of PbSulf1-GHx and GH92_1 follows a linear trend. Net reducing end signal is calculated as absorbance difference between digested and undigested samples. (**B**) *T. weissflogii* biomass was extracted sequentially using hot water, EDTA or NaOH as described in (Vidal). Mannan content in dialyzed extracts were enzymatically quantified after AEX enrichment. ***Dry weight of dialyzed and dried EDTA extracts was below 0.1 mg. (**C**) Processing control to determine loss of mannan during the enrichment procedure with anion exchange chromatography (AEX). Defined amounts of mannan were loaded on three different AEX columns of the same type and recovery was calculated using reducing end signal after enzyme digestion (n=3).

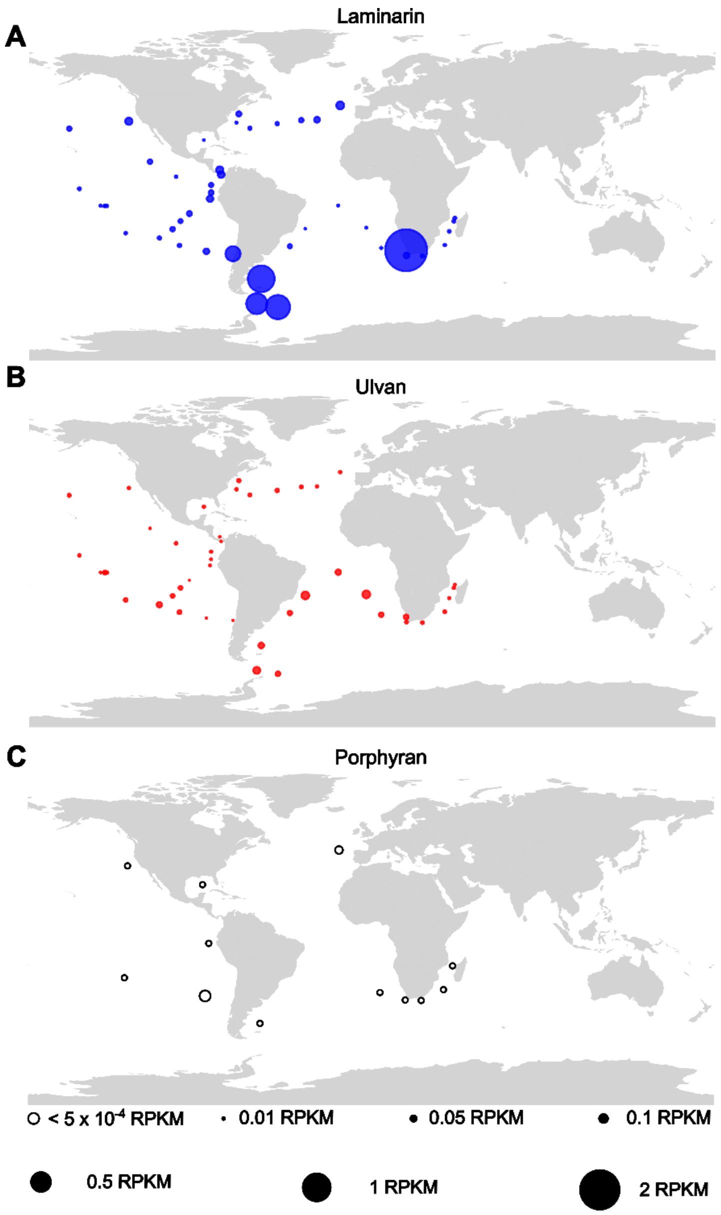


**Fig. S13.**

Global abundance of the mannan polysaccharide utilization locus (PUL) from *Polaribacter* sp. Hel1_33_49 metagenome data from the *TARA* Oceans database. (**A-D**) The sequences from the respective PUL were mapped against the raw reads from *TARA* Oceans at the respective stations. Counts were normalized by the length of the PUL and are displayed as filled circles according to resulting RPKM. (**A**) Laminarin PUL from *Aurantivirga* MAG C_MB344. (**B**) Ulvan PUL from *Formosa agariphila* KMM 3901$^{T}$. (**C**) Porphyran PUL from *Zobellia galactanivorans* Dsij$^{T}$. Porphyran PUL was detected at 12 out of 45 stations with low read abundances. For better visibility these are displayed at lower scale with empty circles. Values ranged from 1-5 × $10^{-4}$ RPKM.

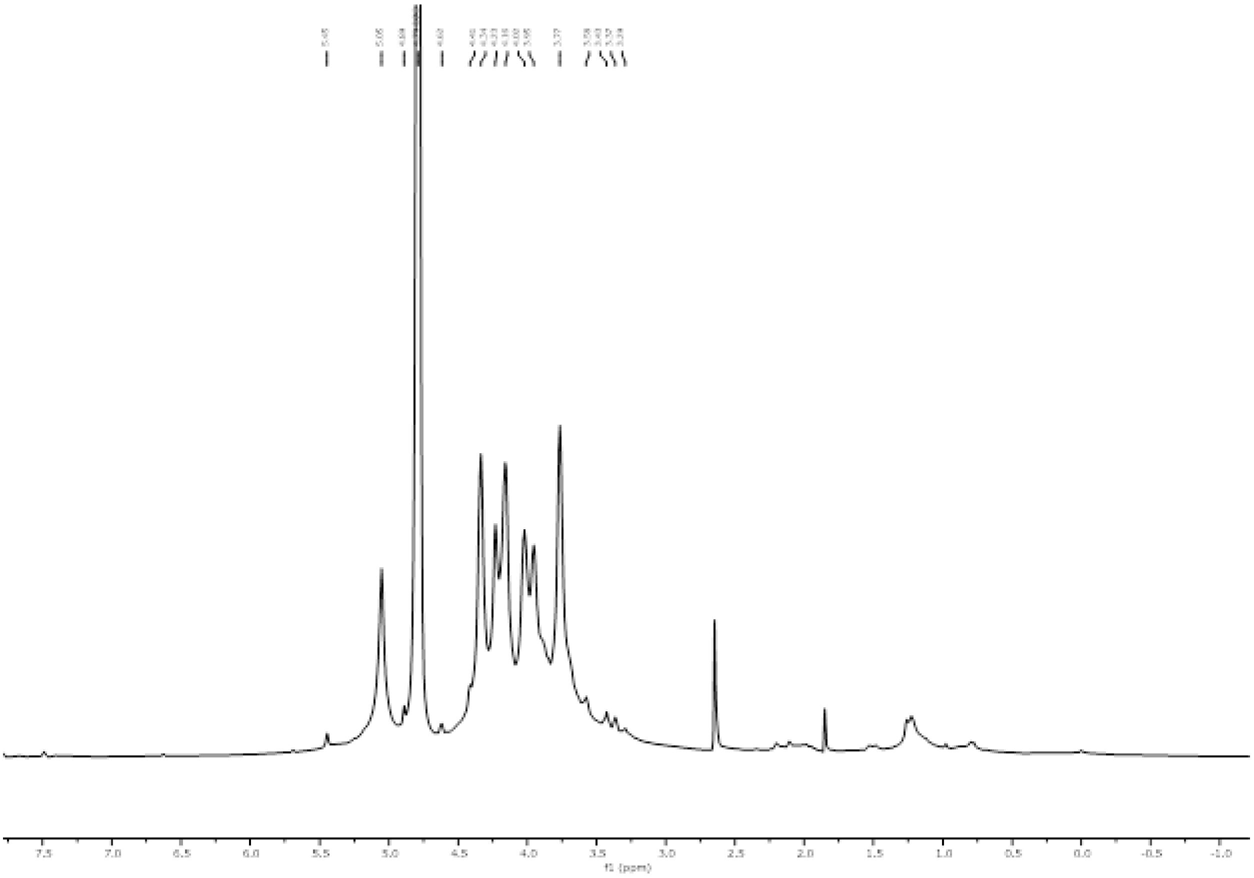


**Fig. S14.** $^{1}$H NMR of the mannan.

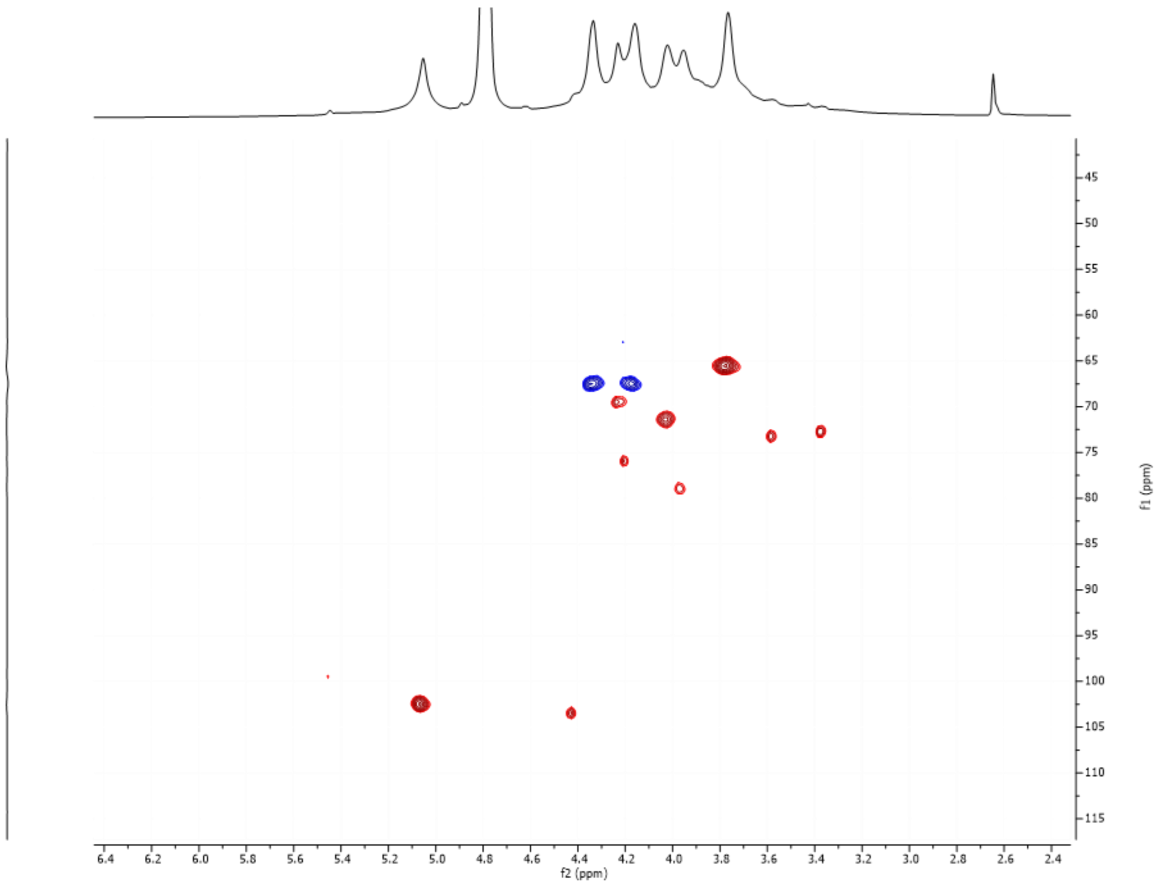

**Fig. S15** $^{1}H$-$^{13}C$ HSQC NMR of the mannan from *T. weissflogii*.

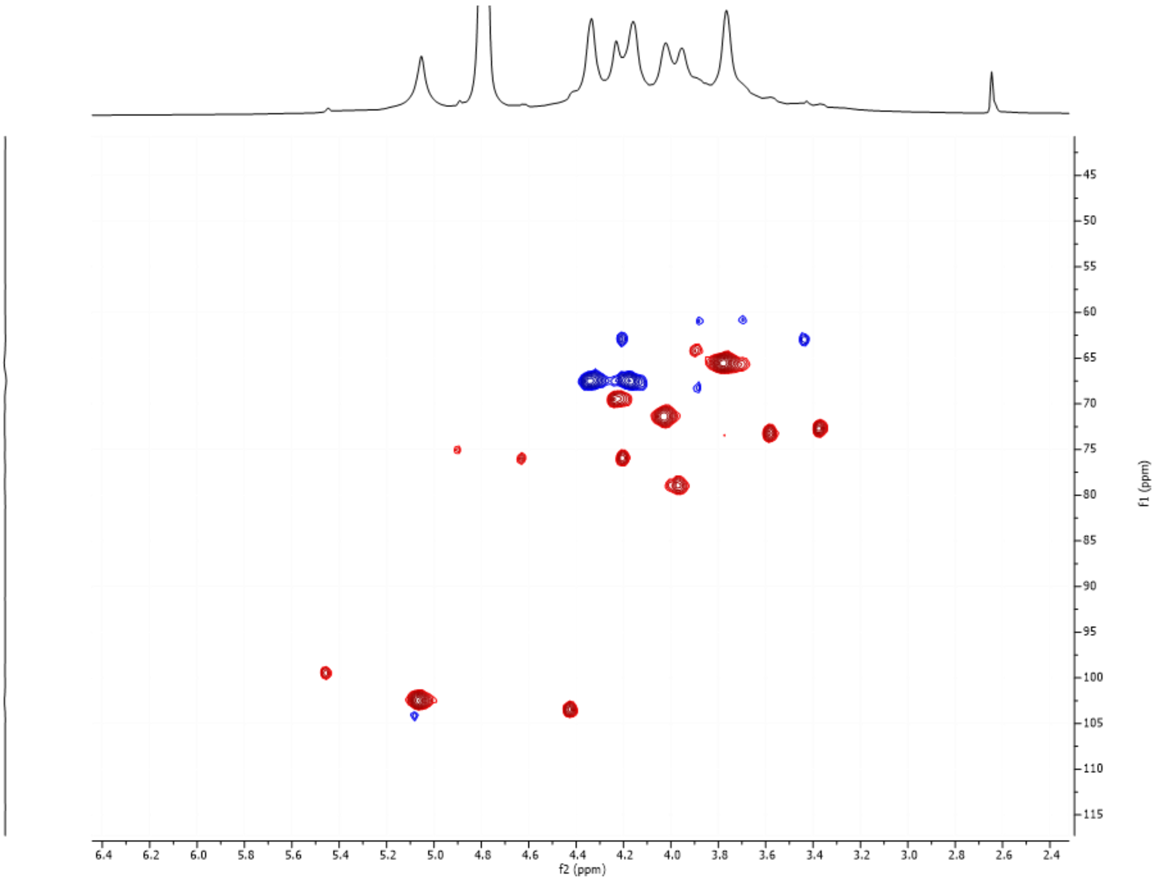


**Fig. S16.** $^{1}H$-$^{13}C$ HSQC NMR mannan from *T. weissflogii* displayed at higher intensity to show minor peaks.

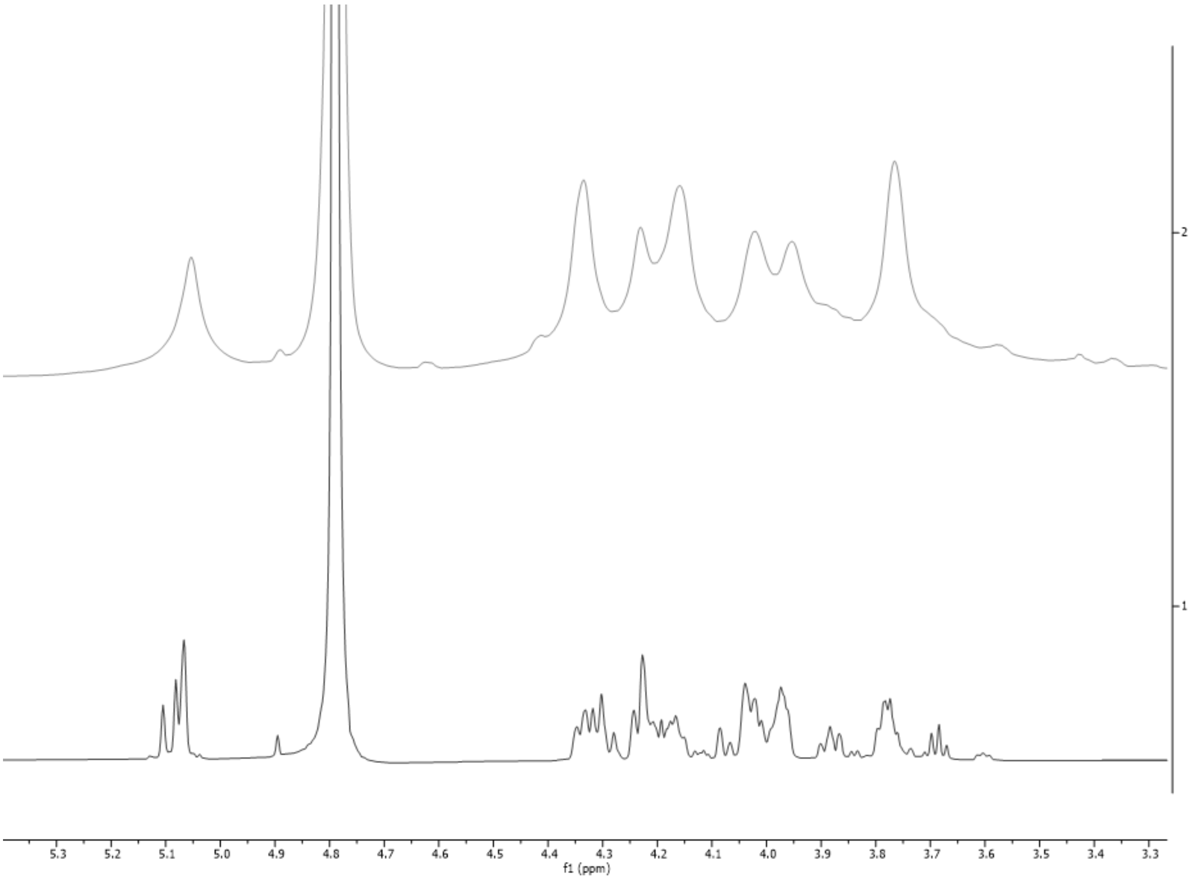


**Fig. S17.** Stacked $^{1}$H NMR of *T. weissflogii* mannan from *T. weissflogii* (top) and synthetic α-1,3 6-*O*-sulfated mannan (bottom).

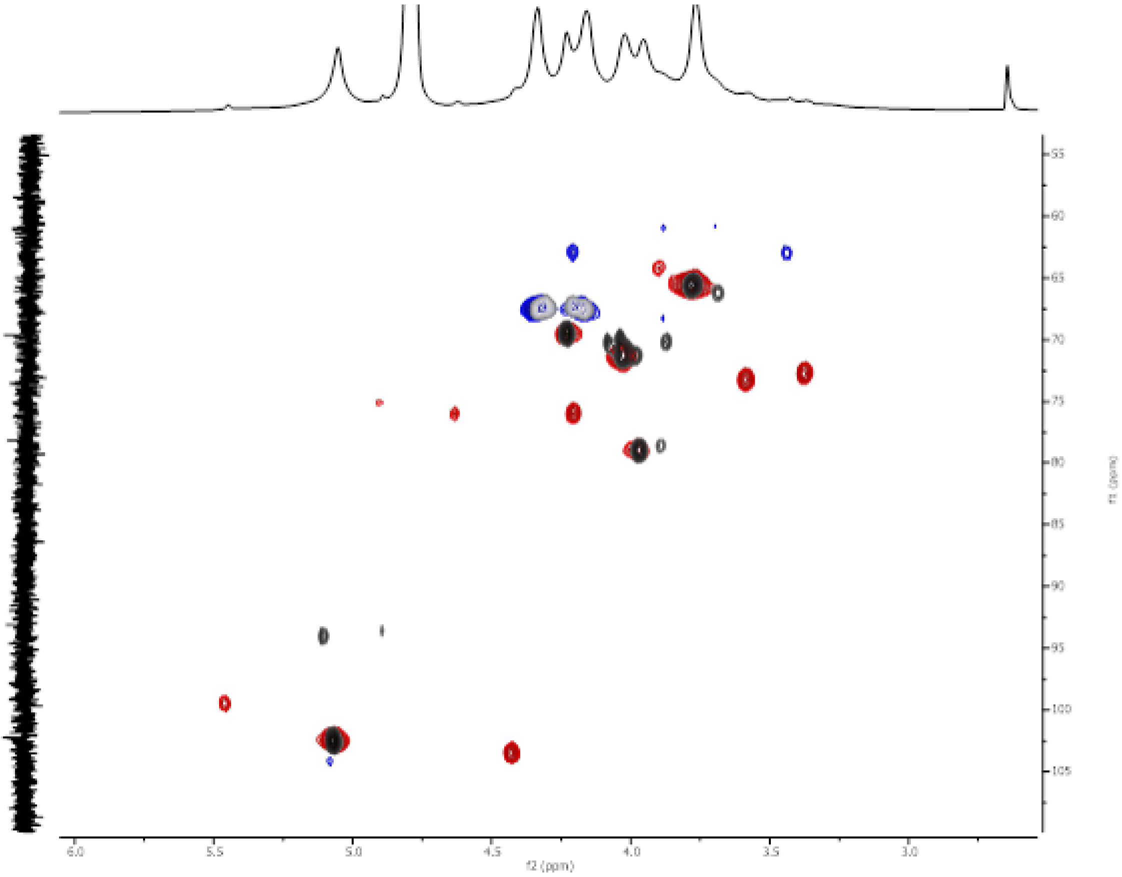

**Fig. S18.** Superimposed $^{1}H$-$^{13}C$ HSQC NMRs of mannan from *T. weissflogii* (red blue) and synthetic α-1,3-6-*O*-sulfated mannan (greyscale).

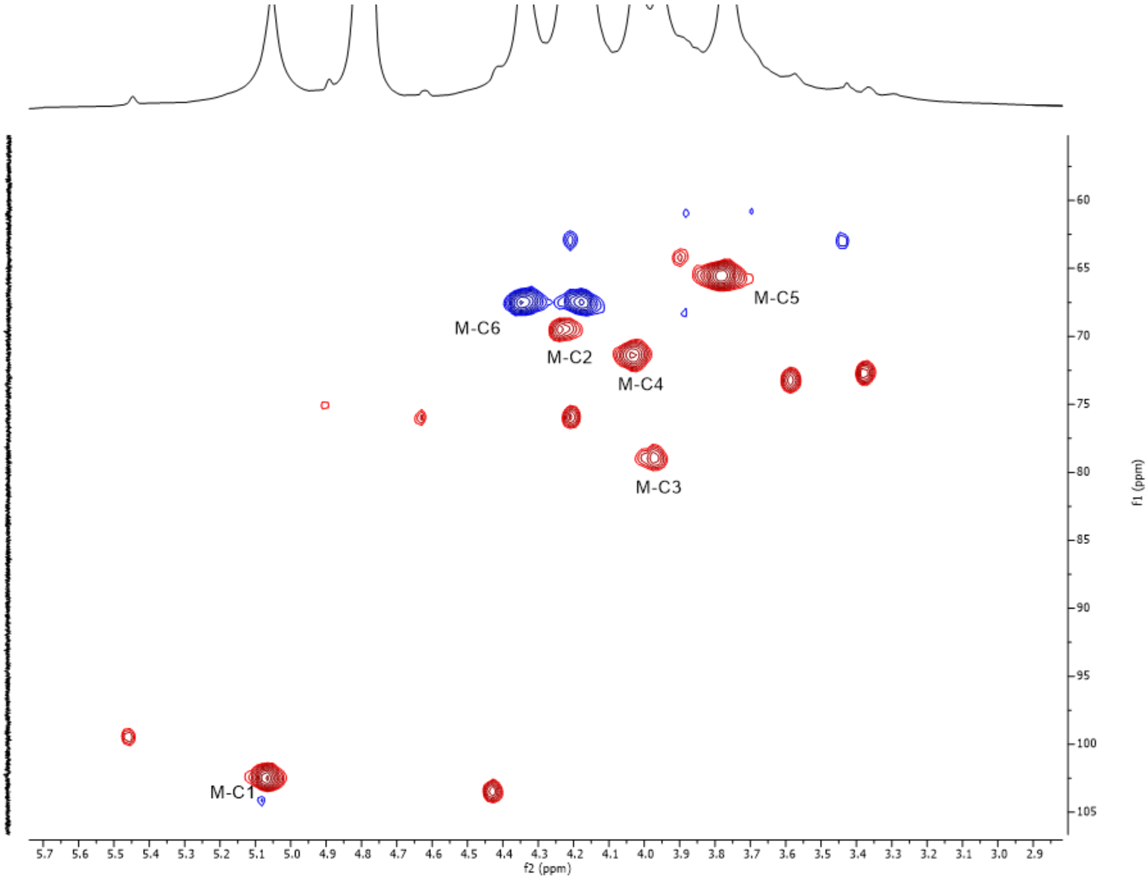


**Fig. S19.** $^{1}H$-$^{13}C$ HSQC NMR of mannan from *T. weissflogii* with major mannose peaks labelled.

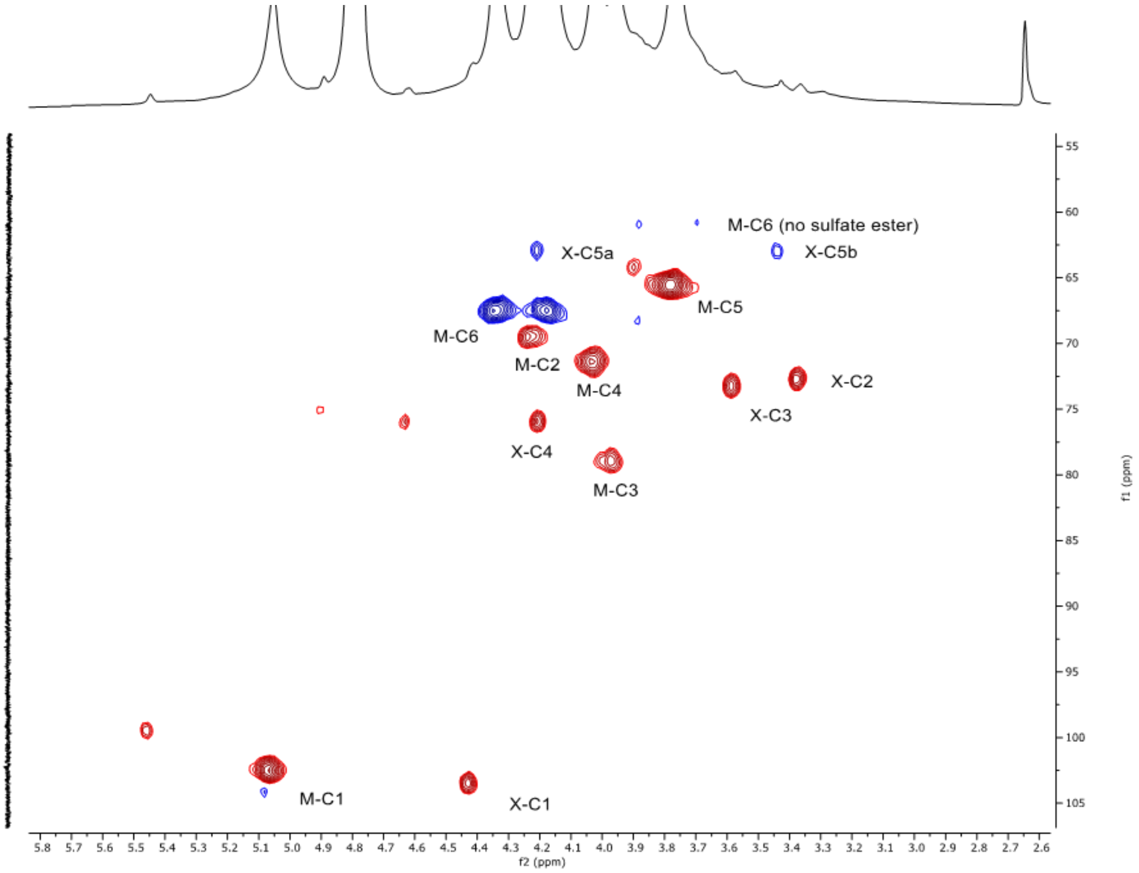


**Fig. S20.** $^{1}H$-$^{13}C$ HSQC NMR mannan with mannose from *T. weissflogii* (M) and possible xylose (X) branch peaks labelled.

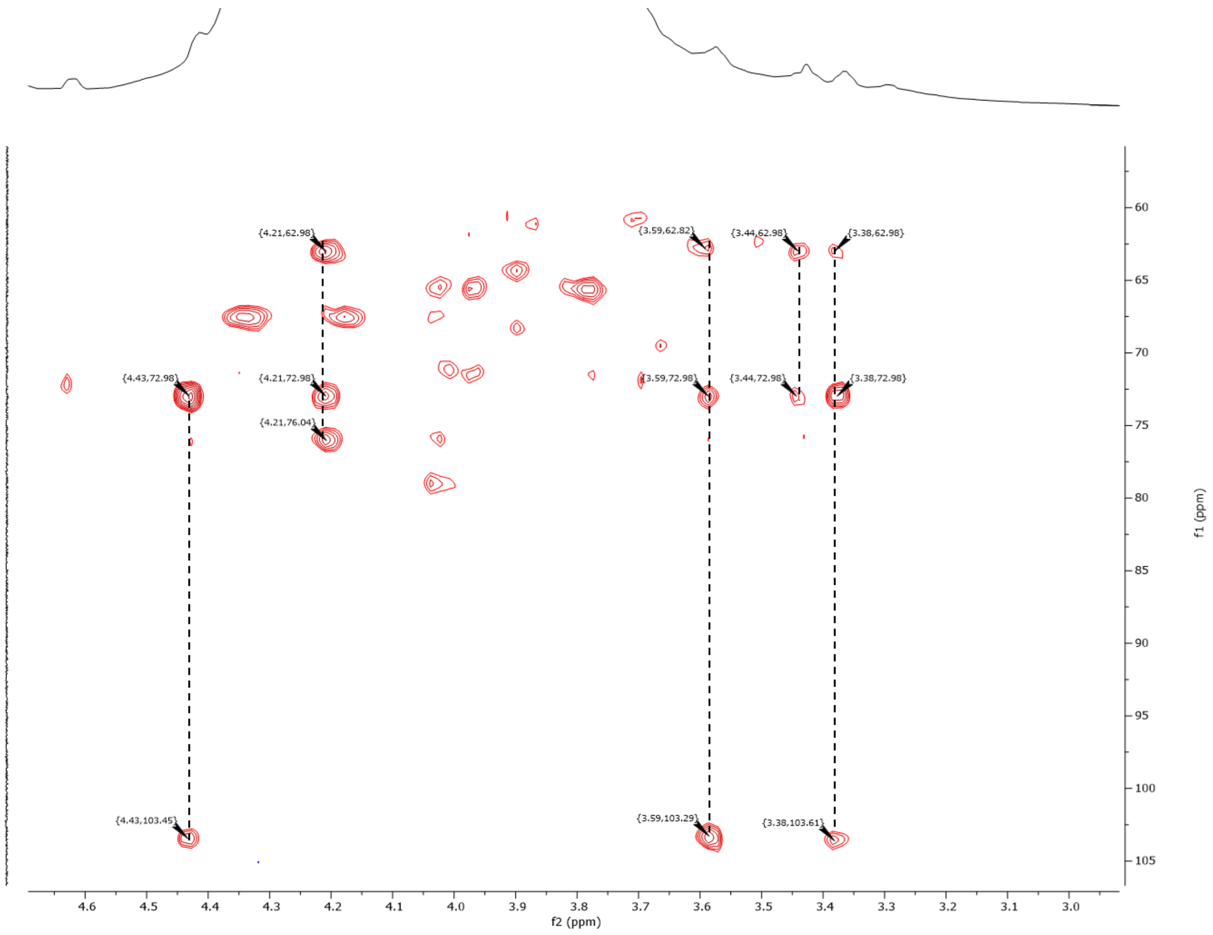


**Fig. S21**. $^{1}H$-$^{13}C$ HSQC TOCSY of mannan from *T. weissflogii*. Highlighted are putative xylose cross-peaks

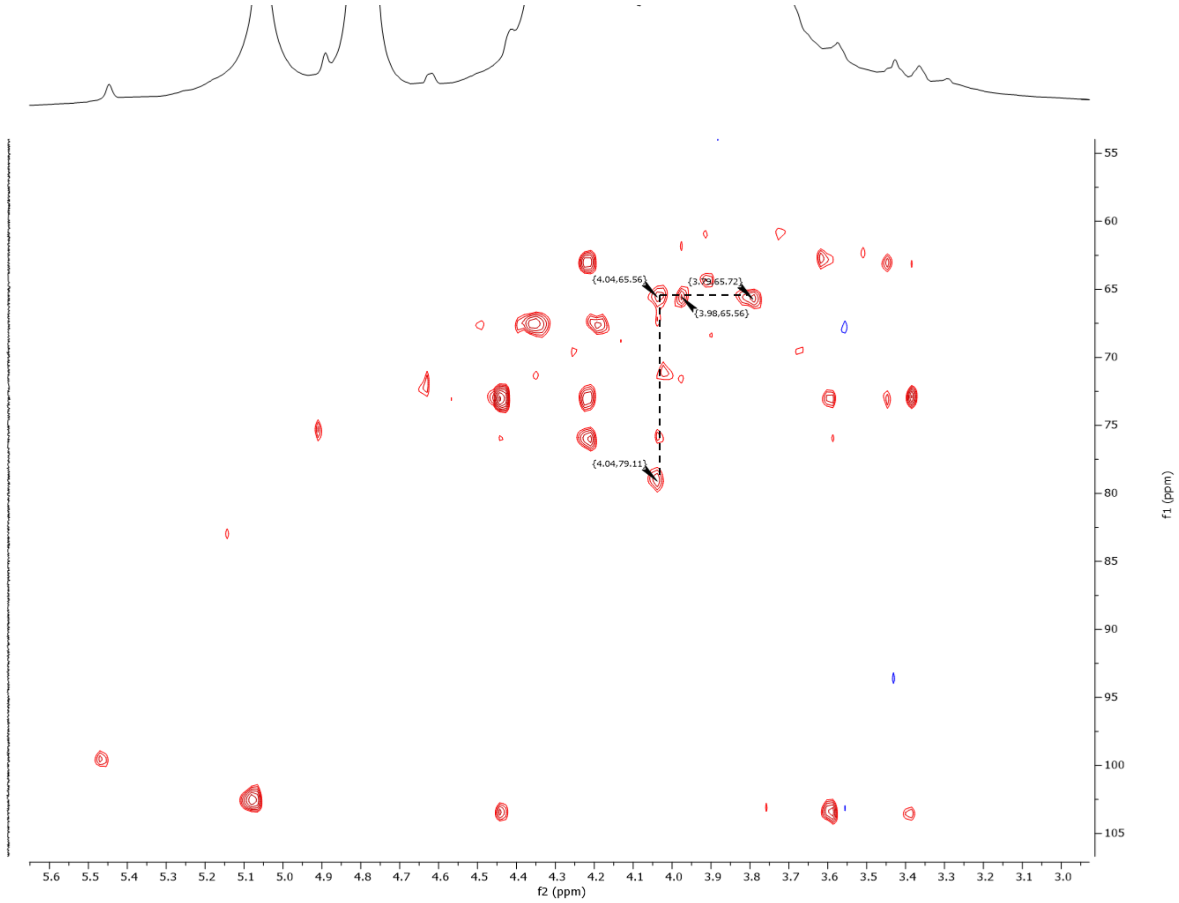


**Fig. S22.** $^{1}H$-$^{13}C$ HSQC TOCSY of *T. weissflogii* mannan. Highlighted are mannose cross-peaks

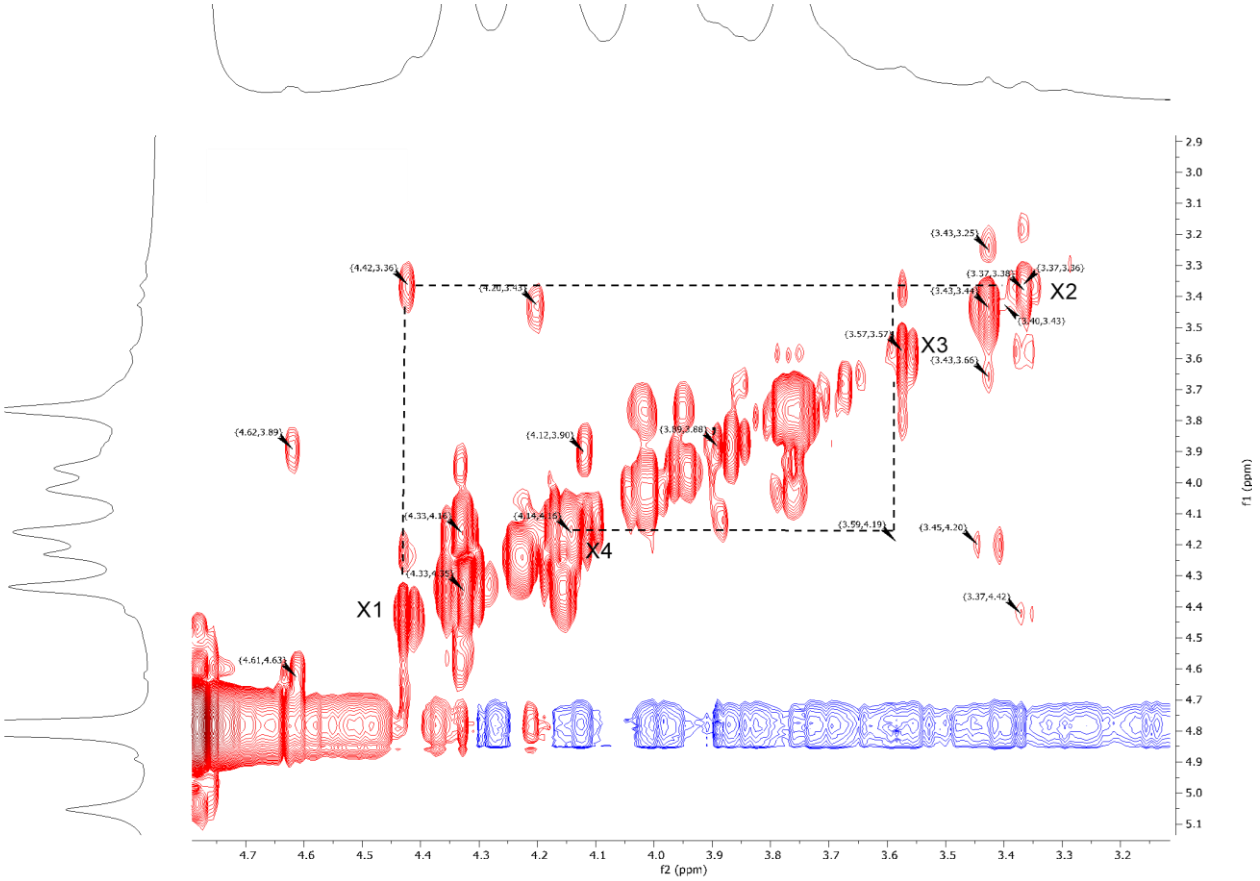

**Fig. S23.** $^1$H-$^1$H COSY of *T. weissflogii* mannan. 4.42 ppm (H-1 xylose-like) vicinal coupling to 3.37 ppm (H-2 xylose-like).

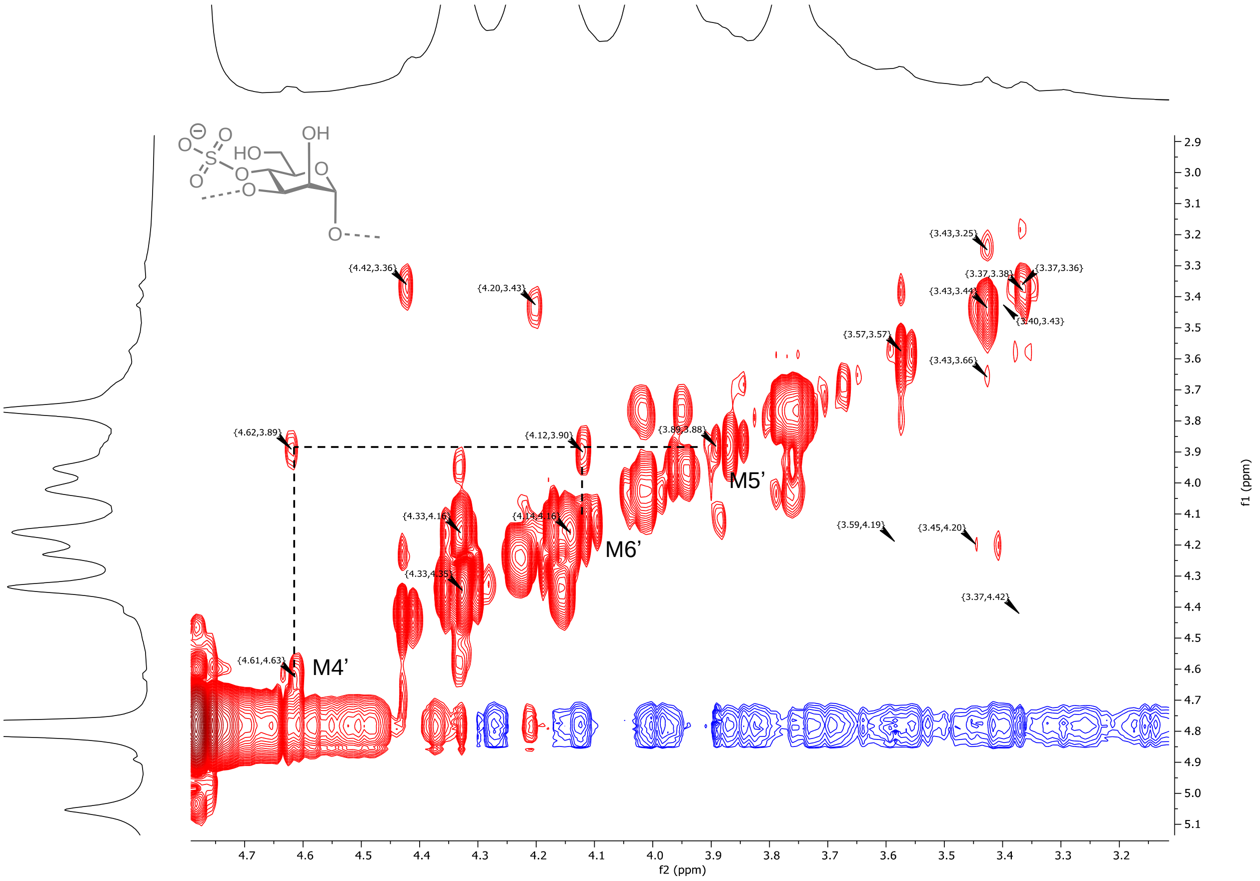


**Fig. S24.** $^1$H-$^1$H COSY of mannan. Mannose (cross-peak 4-*O*-sulfation) coupled a H-5, 4.64 ppm to 3.89 ppm suggesting a minor quantity of 4-*O*-sulfation.

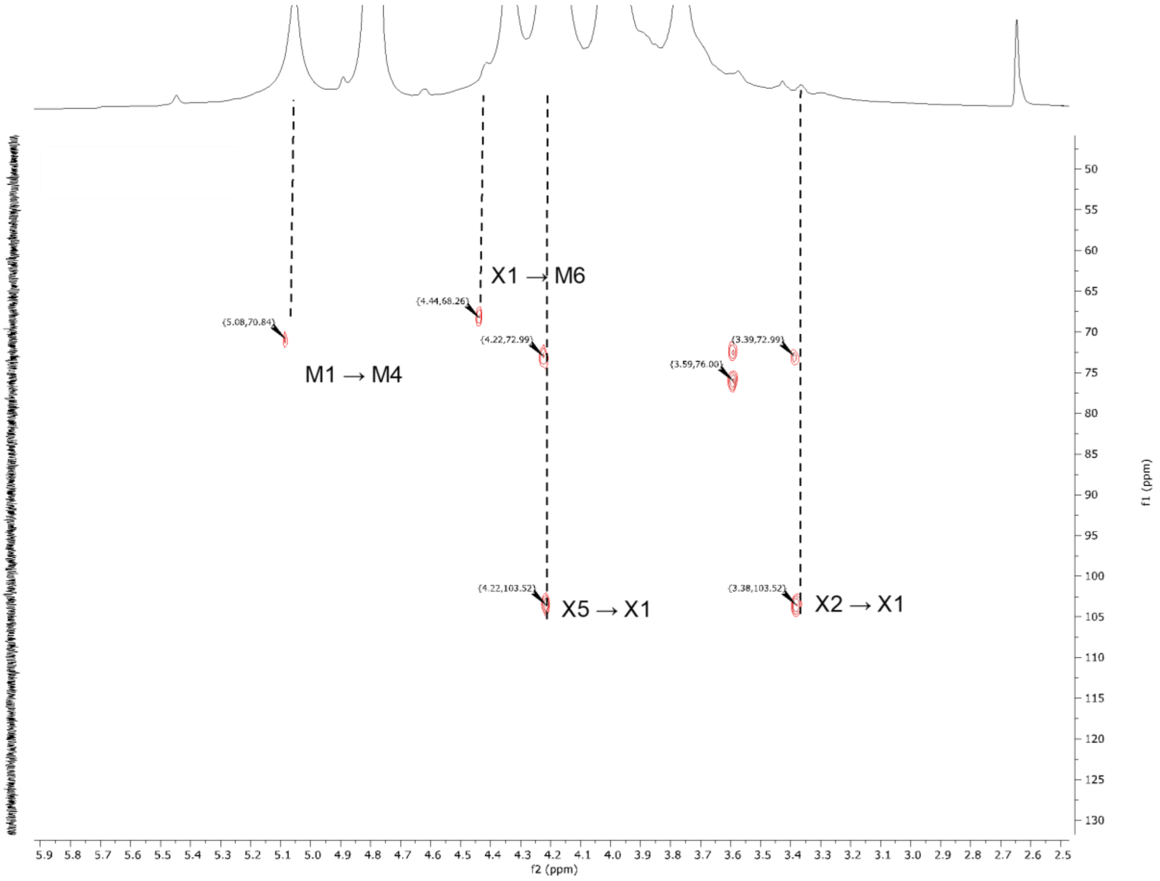


**Fig. S25.** $^{1}H$-$^{13}C$ HMBC of sulfated mannan.

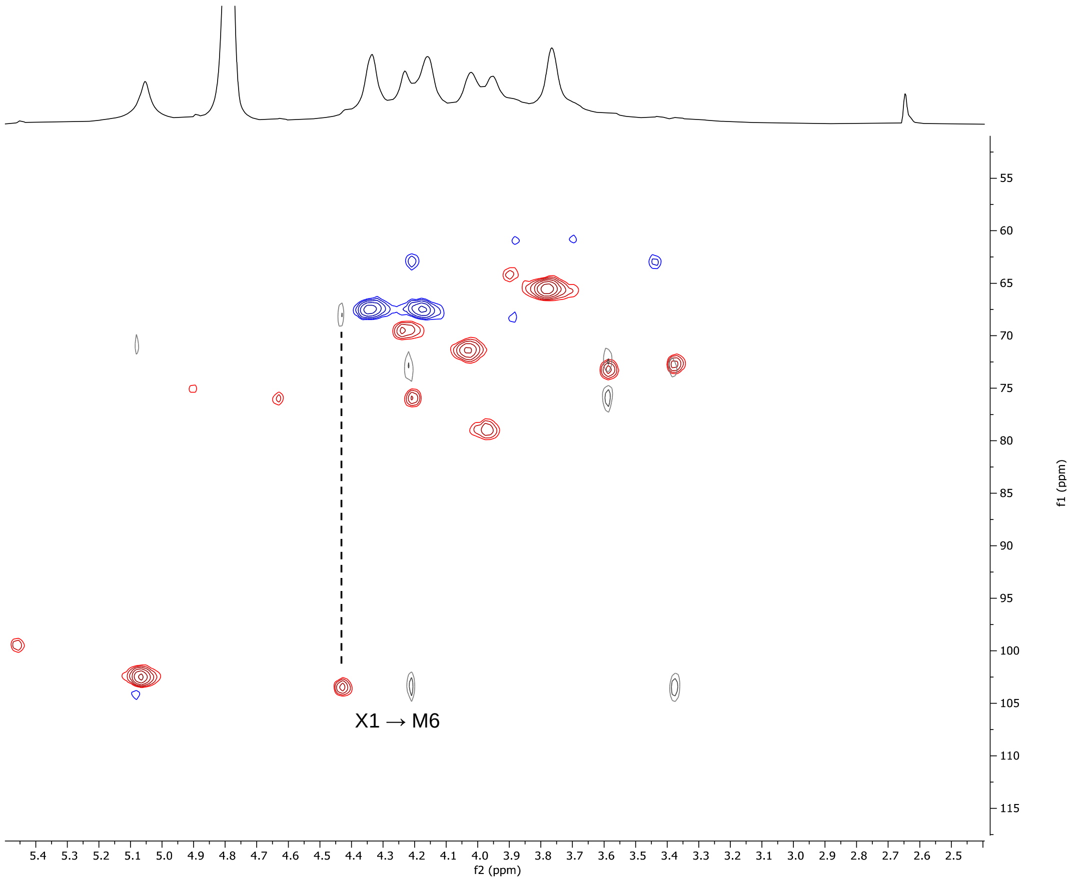

**Fig. S26.** Superimposed $^{1}H$-$^{13}C$ HMBC (greyscale) and $^{1}H$-$^{13}C$ HSQC (red/blue) NMR spectra.

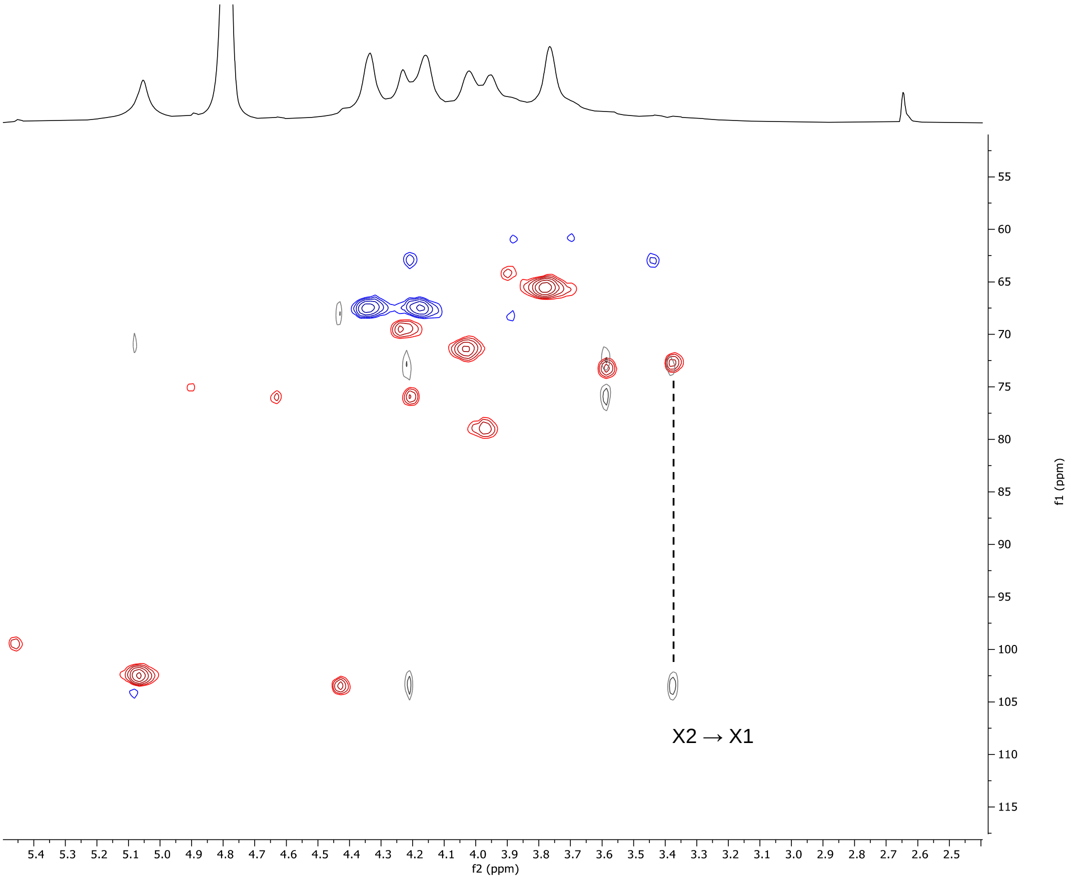


**Fig. S27.** Superimposed $^{1}H$-$^{13}C$ HMBC (greyscale) and $^{1}H$-$^{13}C$ HSQC (red/blue) spectra.

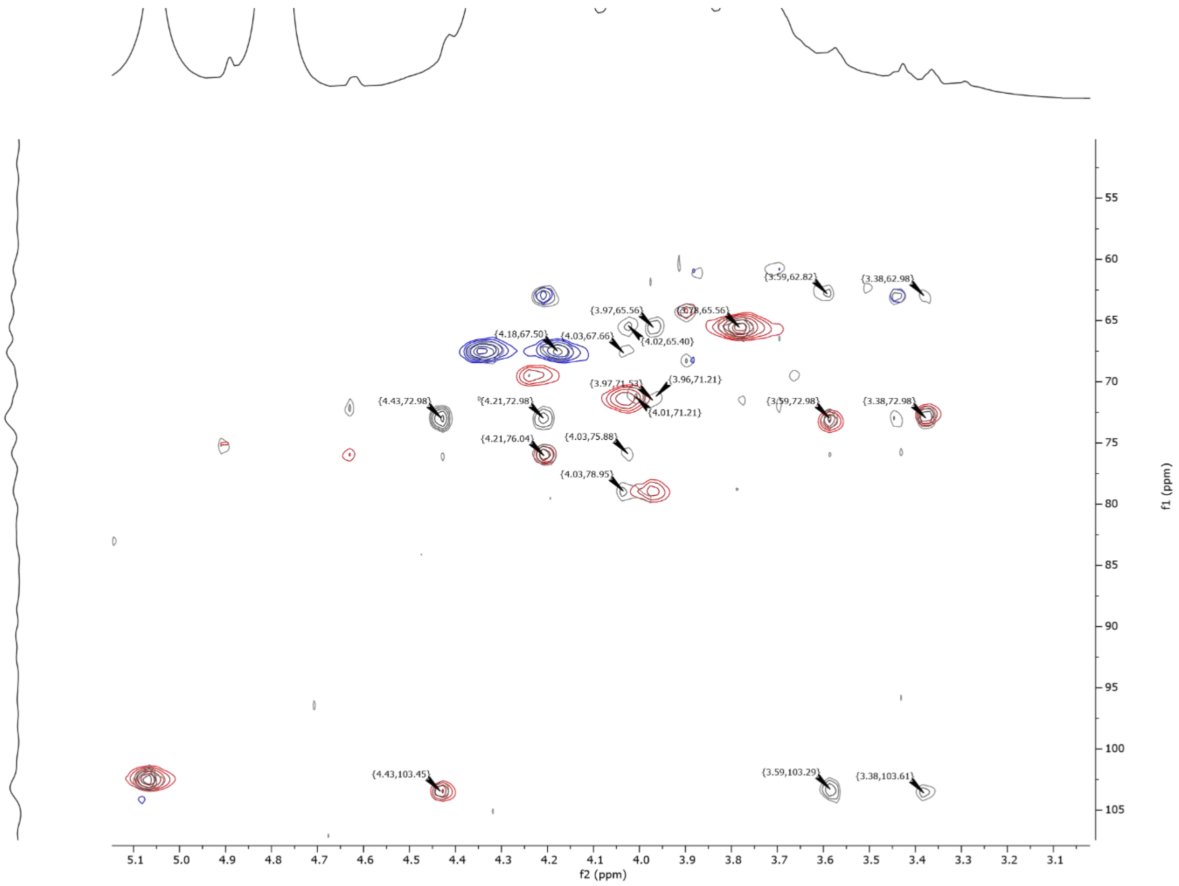


**Fig. S28.** Superimposed $^{1}H$-$^{13}C$ TOSCY (greyscale) and $^{1}H$-$^{13}C$ HSQC (red/blue) spectra.

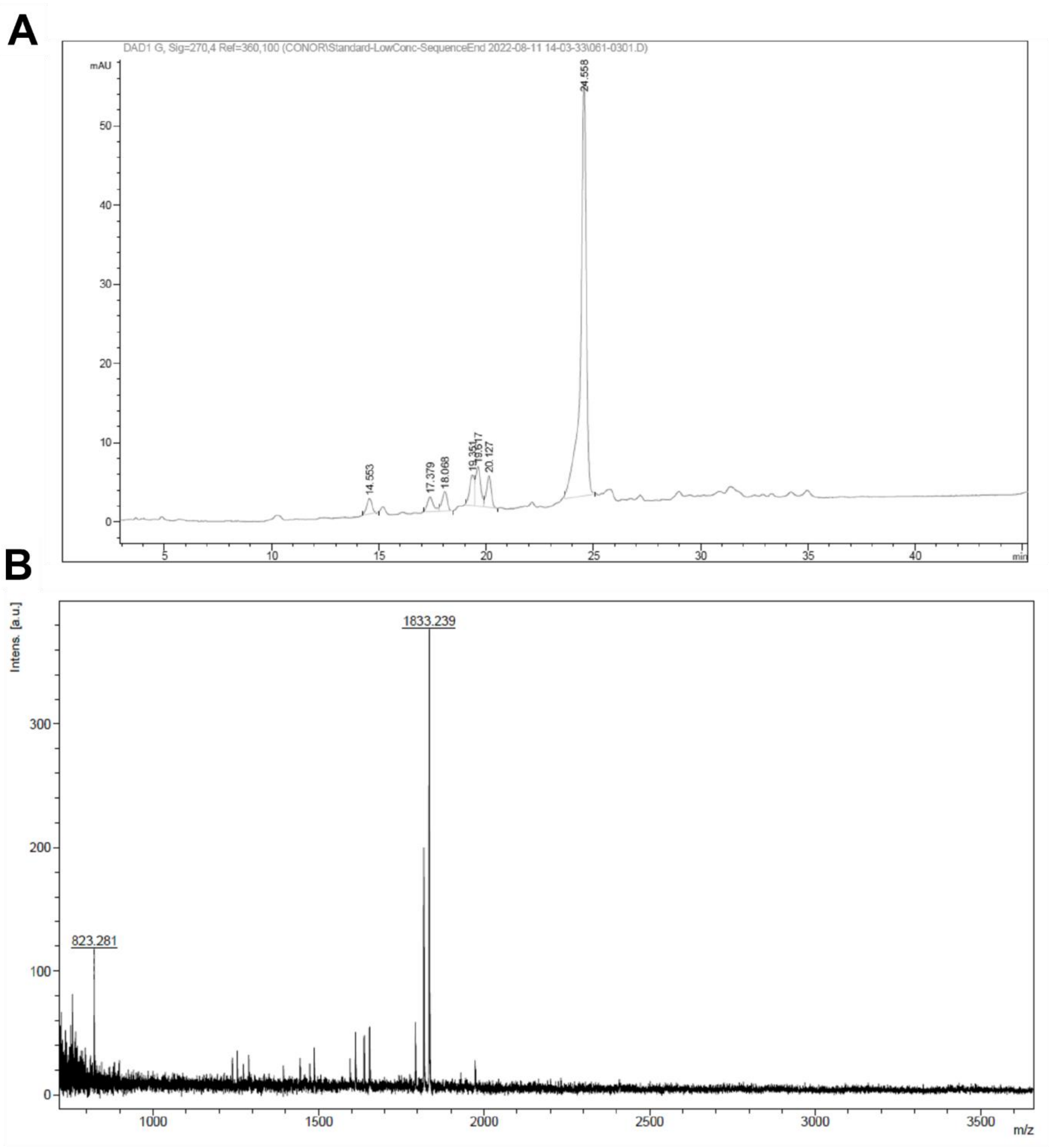


**Fig. S29. Automated assembly of mannan trisaccharide.** (**A**) Crude NP-HPLC of automated assembly up to trisaccharide. Trace is UV 280nm. HPLC Method 1 (20 to 55EA). (**B**) MALDI-TOF spectrum of mannotriose. Chemical Formula: $C_{107}H_{94}KO_{26}$. Expected mass, 1833.5670, observed mass 1833.239.

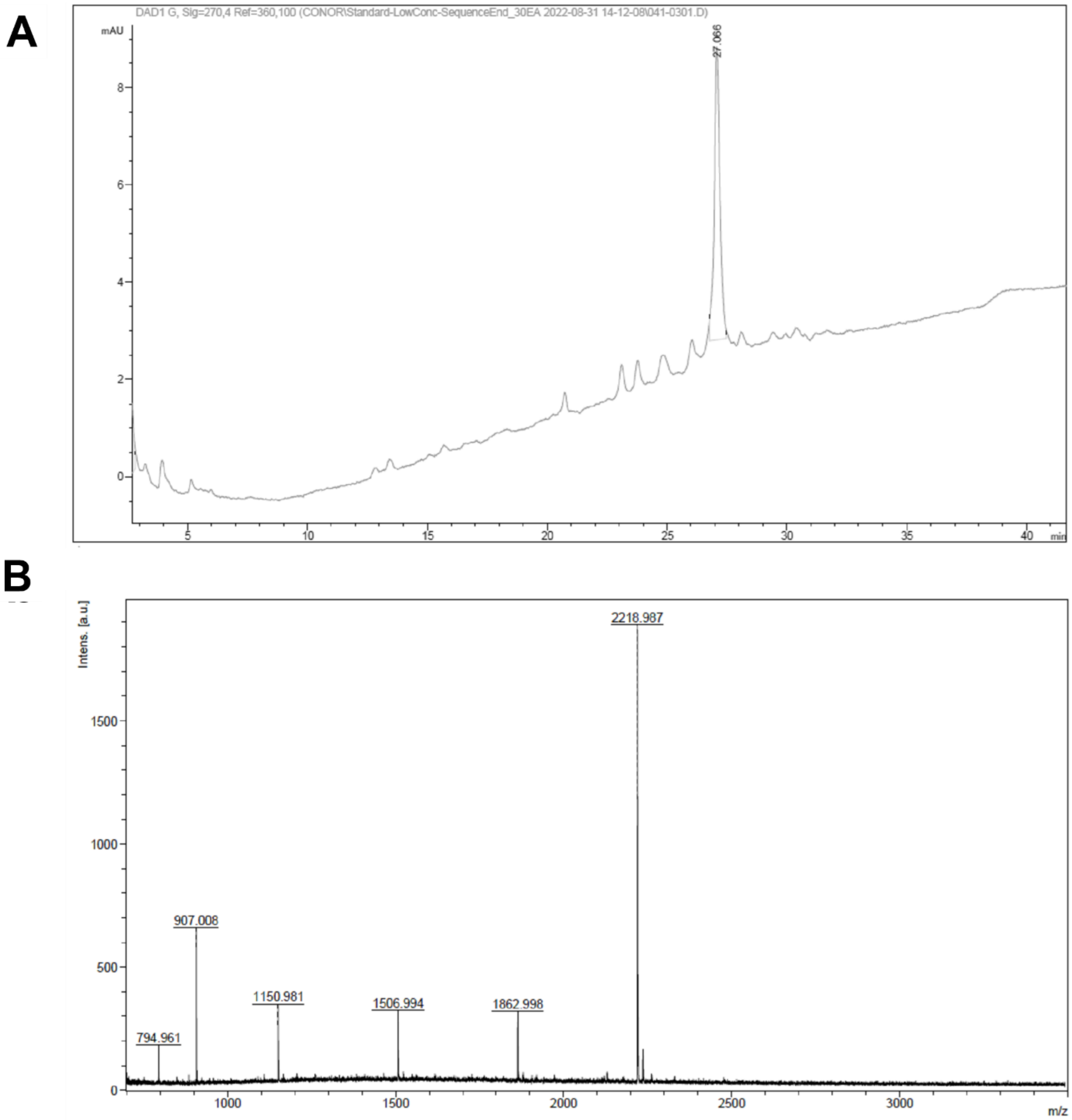


**Fig. S30. a** NP-HPLC of automated assembly of hexasaccharide**.** (**A**) NP-HPLC of 6-mer-t-weiss with no-fmoc. Signal is UV 270nm and HPLC method 2 (30 to 90EA). (**B**) MALDI-TOF of mannose-1,3-hexasacccaride. Note Fmoc protecting groups are removed. Chemical Formula: $C_{122}H_{124}NaO_{38}$. Expected mass, 2219,7668, Observed, 2218.987.

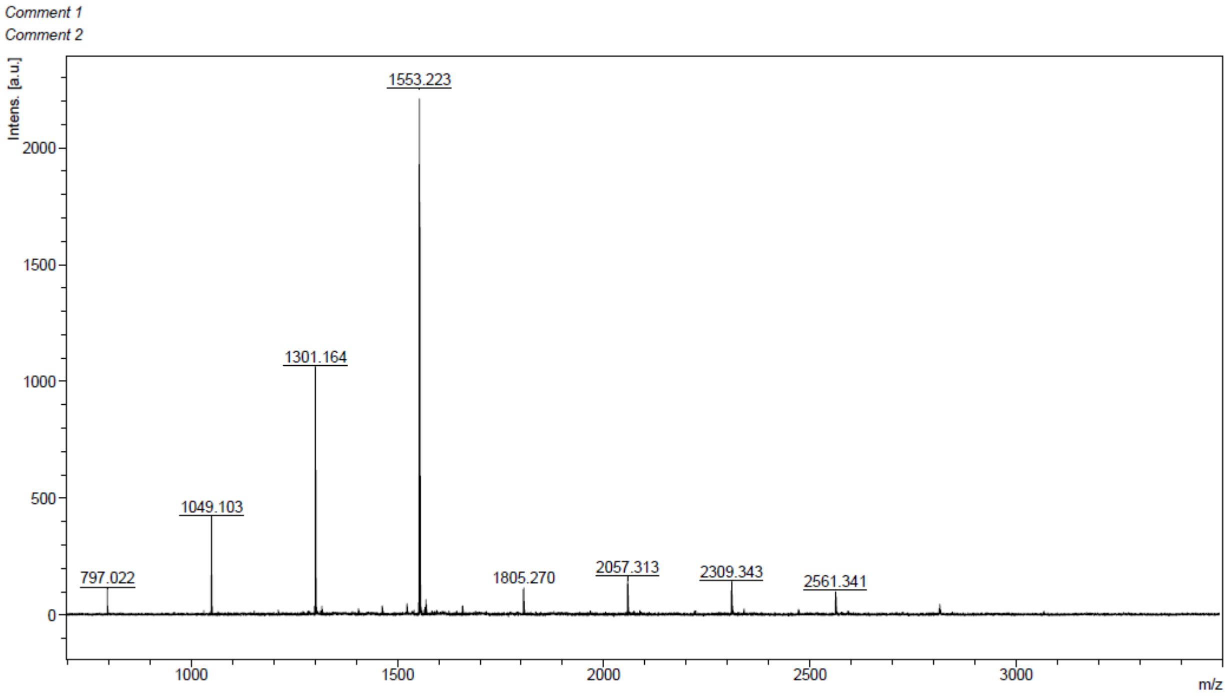


**Fig. S31.** MALDI-TOF of mannose-1,3-hexasacccaride with ester protecting groups removed. Chemical Formula: $C_{78}H_{98}NaO_{31}$. Expected mass, 1553,5990; Observed, 1553,223.

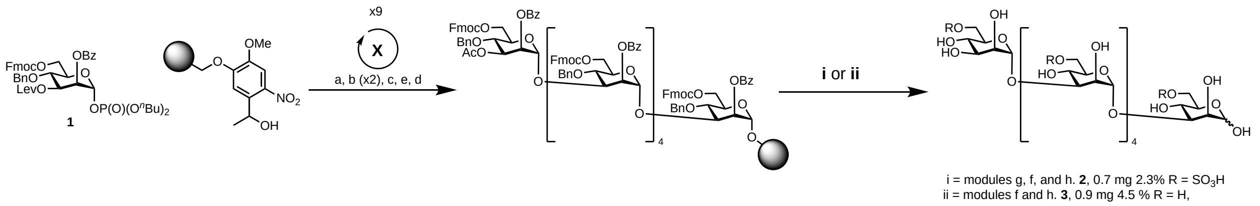


**Scheme S1.** Automated glycan assembly of sulfated mannan oligosaccharide.

**Table S1.**

Comparison of the *T. weissflogii* mannan to a synthetic standard. *difference is *T. weissflogii* mannan minus synthetic glycan.

| Mannan (Synthetic) | $^{1}H$ | $^{13}C$ | 1H Δδ (ppm) | 13C Δδ (ppm) |
|---|---|---|---|---|
| **H-1** | 5.07 (5.07) | 102.49 (102.49) | - | - |
| **H-2** | 4.24 (4.23) | 69.43 (69.43) | 0.01ppm | - |
| **H-3** | 3.97 (3.97) | 78.95 (78.85) | - | 0.1 ppm |
| **H-4** | 4.03 (4.03) | 71.37 (71.21) | - | 0.16 ppm |
| **H-5** | 3.78 (3.78) | 65.56 (65.56) | - | - |
| **H-6a** | 4.34 (4.31) | 67.50 (67.50) | 0.03 ppm | - |
| **H-6b** | 4.18 (4.17) | 67.50 (67.50) | 0.01 ppm | - |

**Table S2.**

*Polaribacter* spp. grown in HaHa 100V medium with or without mannan from *T. weissflogii*

| **Growth of strain ($OD_{600nm}$)** | Medium | Mannan |
|---|---|---|
| *Polaribacter* sp. Hel1_33_49 | 0.168 | 0.410 |
| *Polaribacter* sp. Hel1_33_78 | 0.187 | 0.411 |
| *Polaribacter* sp. Hel1_33_96 | 0.149 | 0.383 |
| *Polaribacter* sp. Hel1_88 | 0.109 | 0.212 |

**Table S3.**

Activity assays of enzymes on mannan from *T. w.* (0.5-5M fraction after AEX), pure mannan or on synthetic substrates. n.d.: no activity detected. -/-: not tested.

| Source Organism | Locus Tag | Family | Predicted localization | Activity on anionic exudates form *T. weissflogii* | Activity on pure mannan | Activity on pnp-substrate |
|---|---|---|---|---|---|---|
| *Polaribacter* sp. Hel1_33_78 | Pb1068 | S1_15 | SPI (75%) | | n.t. | n.t. |
| | Pb1069 | S1_N.C. | SPII (96%) | | n.t. | n.t. |
| | Pb1070 | S1_51 | SPI (89%) | | n.t. | n.t. |
| | Pb1072 | S1_53 | SPII (99%) | | n.t. | n.t. |
| | Pb1095 | S1_71 | SPII (99%) | | n.t. | n.t. |
| | Pb1096 | S1_11 | SPII (99%) | | n.t. | n.t. |
| | Pb1097 | S1_71 | SPII (85%) | | n.t. | n.t. |
| | PbSulf2 (Pb1078) | S1_11 | SPII (99%) | ✓ | ✓ | n.t. |
| | PbSulf1-GHx (Pb1059) | S1_51 GHx | T9SS | ✓ | ✓ | n.t. |
| | Pb1065 | GH92 | SPII (95%) | | n.t. | n.d |
| | Pb1066 | GH92 | none | | n.t. | pnp-α-D-Mannose |
| | PbGH92_1 (Pb1075) | GH92 | SPI (94%) | ✓ | ✓ | n.t. |
| | Pb1076 | GH3 | SPI (96%) | | n.t. | pnp-β-D-Galactose |
| | Pb1077 | GH3 | SPI (89%) | | n.t. | n.d |
| | Pb1081 | GH92 | SPI (93%) | | n.t. | pnp-α-D-Mannose |
| | Pb1091 | GH88 | none | | n.t. | n.d |
| | Pb1093 | GH92 | SPII (99%) | | n.t. | n.d |
| | Pb1094 | GH2 | SPI (85%) | | n.t. | n.d |
| | Pb1063 | GH99-like | none | | n.t. | n.t. |
| | Pb1064 | GH99-like | none | | n.t. | n.d |
| | Pb1071 | CE | SPI (88%) | | n.t. | n.t. |
| | Pb1100 | PLx | T9SS | ✓ | n.d. | n.t. |
| | Pb1053 | Hypothetical | SPI (91%) | | n.t. | n.t. |
| | Pb1054 | Hypothetical | SPII (81%) | | n.t. | n.t. |
| | Pb1055 | Hypothetical | T9SS | | n.t. | n.t. |
| | Pb1057 | Hypothetical | T9SS | | n.t. | n.t. |
| | Pb1058 | Hypothetical | T9SS | | n.t. | n.t. |
| *Ochrovirga pacifica* S85 | OpSulf1 | S1_51 | SPI (50%) SPII (50%) | ✓ | ✓ | n.t. |
| | OpGHx | GHx | SPI (75%) SPII (25%) | ✓ | ✓ | n.t. |

**Table S4.**

Data collection and refinement statistics for OpSulf1.

| | |
|---|---|
| **X-ray source** | DESY P11 |
| **Wavelength (Å)** | 1.0332 |
| **Space group** | P 43 |
| **Unit cell** | |
| **a, b, c (Å)** | 115.27 115.27 169.83 |
| **α, β, γ (º)** | 90 90 90 |
| **Resolution range, (Å)** | 44.07 - 1.489 (1.51 - 1.49) |
| **R-merge** | 0.08627 (1.916) |
| **Completeness (%)** | 99.9 (98.06) |
| **Multiplicity** | 13.7 (13.2) |
| **Mean I/sigma(I)** | 15.64 (1.54) |
| **No. of reflections** | 4948693 (156085) |
| **No. of unique reflections** | 359919 (11806) |
| **Mosaicity** | 0.060 |
| ***Refinement*** | |
| **$R_{work}/R_{free}$** | 0.1719/0.1925 |
| **Number of non-hydrogen atoms** | 18258 |
| **Macromolecules** | 16836 |
| **Water** | 1418 |
| **Ligands** | 4 |
| **Protein residues** | 2076 |
| **B factors** | |
| **Overall** | 23.44 |
| **Protein** | 23.01 |
| **Water** | 28.64 |
| **Ligands** | 22.36 |
| **R.m.s deviations** | |
| **Bond lengths (Å)** | 0.018 |
| **Bond angles (º)** | 1.83 |
| **Ramachandran statistics (%)** | |
| **Favored** | 96.52 |
| **Allowed** | 3.48 |
| **Outliers** | 0.0 |
| **Rotamer outliers (%)** | 0.61 |
| **Clashscore** | 3.00 |
| **PDB accession code** | 9FVT |

Statistics for the highest-resolution shell are shown in parentheses.

**Table S5.**

Confirmation of sulfated mannose position based on retention time and MS/MS. Fragmentation ions generated from HCD fragmentation of 259.0129 *m/z* with a window of 0.4 Da. Mannan was digested with PbSulf1-GHx and PbGH92_1.

| **Sample** | **Retention time (min)** | ***m/z*** | **Relative abundance** | **Ion formula** | **Annotation** |
|---|---|---|---|---|---|
| Synthetic mannose-2-sulfate | 3.0514 | 80.9637 | 0.11 | $HSO_3$ | $HSO_3$ |
| | | 96.9588 | 1.00 | $HSO_4$ | $HSO_4$ |
| | | 108.9007 | 0.37 | $CHSO_4$ | $^{1,2}A$ |
| | | 138.9697 | 0.94 | $C_2H_3O_5S$ | $^{1,3}A$, $^{0,2}X$ |
| | | 241.0037 | 0.11 | $C_6H_9O_8S$ | $[M-H_2O-H]^-$ |
| | | 259.0131 | 0.59 | $C_6H_{11}O_9S$ | $[M-H]^-$ |
| Synthetic mannose-3-sulfate | 2.5346 | 96.9587 | 1.00 | $HSO_4$ | $HSO_4$ |
| | | 108.9006 | 0.03 | $CHSO_4$ | $^{3,4}A$ |
| | | 168.9803 | 0.01 | $C_3H_5O_6S$ | $^{0,3}X$, $^{1,4}A$ |
| | | 259.0131 | 0.31 | $C_6H_{11}O_9S$ | [M-H]- |
| Synthetic mannose-4-sulfate | 1.3593 | 96.959 | 1.00 | $HSO_4$ | $HSO_4$ |
| | | 138.9699 | 0.35 | $C_2H_3O_5S$ | $^{2,4}A$ |
| | | 180.9804 | 0.08 | $C_4H_5O_6S$ | $^{0,4}X-H_2O$ |
| | | 198.9915 | 0.68 | $C_4H_7O_7S$ | $^{0,4}X$ |
| | | 241.0026 | 0.02 | $C_6H_9O_8S$ | $[M-H_2O-H]^-$ |
| | | 259.0132 | 0.73 | $C_6H_{11}O_9S$ | [M-H]- |
| Synthetic mannose-6-sulfate | 4.138 | 96.9588 | 1.00 | $HSO_4$ | $HSO_4$ |
| | | 138.9697 | 0.43 | $C_2H_3O_5S$ | $^{0,4}A$ |
| | | 168.9805 | 0.06 | $C_3H_5O_6S$ | $^{0,3}A$, $^{1,4}X$ |
| | | 198.9913 | 0.63 | $C_4H_7O_7S$ | $^{0,2}X$, $^{1,3}A$, $^{2,4}X$ |
| | | 259.0131 | 0.38 | $C_6H_{11}O_9S$ | [M-H]- |
| Digest replicate 1 | 4.2803 | 96.9587 | 1.00 | $HSO_4$ | $HSO_4$ |
| | | 138.9699 | 0.36 | $C_2H_3O_5S$ | $^{0,4}A$ |
| | | 168.9808 | 0.08 | $C_3H_5O_6S$ | $^{0,3}A$, $^{1,4}X$ |
| | | 198.9912 | 0.65 | $C_4H_7O_7S$ | $^{0,2}X$, $^{1,3}A$, $^{2,4}X$ |
| | | 259.0133 | 0.45 | $C_6H_{11}O_9S$ | [M-H]- |
| Digest replicate 1 | 4.2105 | 96.9589 | 1.00 | $HSO_4$ | $HSO_4$ |
| | | 138.9699 | 0.48 | $C_2H_3O_5S$ | $^{0,4}A$ |
| | | 168.9807 | 0.08 | $C_3H_5O_6S$ | $^{0,3}A$, $^{1,4}X$ |
| | | 198.9914 | 0.67 | $C_4H_7O_7S$ | $^{0,2}X$, $^{1,3}A$, $^{2,4}X$ |
| | | 259.013 | 0.39 | $C_6H_{11}O_9S$ | [M-H]- |

**Table S6.** Primers used for cloning of PUL enzymes. Genes marked with an asterisk were cloned using Gibson-assembly into a regular pet28A vector.

| Locus Tag | Primer Forward (5‘ → 3‘) | Primer Reverse (5‘ → 3‘) |
|---|---|---|
| **pet28A_P1** | AGCGTTAAUGTCTGGCTTCTGATAAAG | ATTCCTCTCCCUATAGTGAGTCGTATTAATTTC |
| **pet28A_P2** | AGGGAGAGGAAUTGTGAGCGGATAAC | ATTAACGCUCTCTGGAGAAACTCAACG |
| **pet28A** | ATCCGAAUTCGAGCTCCGTCG | ACCGCTGCUGTGATGATGATGATGATGG |
| **Pb1053** | AGCAGCGGUTGTTTAATCTCCTCTCTTG | ATTCGGAUTTACTTATTTTTATTTTCTAAG |
| **Pb1054** | AGCAGCGGUTCTAAACACGCTAAAACTAG | ATTCGGAUTTATTTTCGATTATTTACTTTTC |
| **Pb1055** | AGCAGCGGUCAACAAACAGTTTATGTTTC | ATTCGGAUTTAACTAATGTTTTCTGACAC |
| **Pb1057** | AGCAGCGGUCAAACTACCGTTTATGTTTC | ATTCGGAUTTAAACAGAAAGTTTAGTTGC |
| **Pb1058** | AGCAGCGGUATGAAAACAATCCAATTAATATC | ATTCGGAUTTACATACTAATGTTTTCTAAGAC |
| **Pb1059** | AGCAGCGGUCAAACCCAAAAACC | ATTCGGAUTTAGCTAAATTTCTCATTTG |
| **Pb1060** | AGCAGCGGUCAAGAAACTATAAAAGTAGAGGG | ATTCGGAUCAGTAAAGCAACTTAGATTGC |
| **Pb1062** | AGCAGCGGUTGCACAAACAAATTTGAAG | ATTCGGAUCTGAAGTTACAGATACAATAGGG |
| **Pb1063** | AGCAGCGGUATGATATCTTTTTCATGTAC | ATTCGGAUGAGATTGATTTGATTTTC |
| **Pb1064** | AGCAGCGGUATGCTAACGCTTTTC | ATTCGGAUGGATTTTTATTTCGTTG |
| **Pb1065** | AGCAGCGGUTGTAGTAATTCAGAAAACAG | ATTCGGAUTTCATAGGTAATCTGTTG |
| **Pb1066** | AGCAGCGGUAAAATCACAAAAACAAC | ATTCGGAUTGTTACTCTTTAATTATTTAGG |
| **Pb1068*** | CTGGTGCCGCGCGGCAGCCATATGGCTAGCAAA AAACCAAATGTCATTGTAT | ATCTCAGTGGTGGTGGTGGTGGTGCTCGAGTTAA TTATTTTTCCAAATTTC |
| **Pb1069*** | CTGGTGCCGCGCGGCAGCCATATGGCTAGC AAGTCTAAAAACGAACTTGTA | ATCTCAGTGGTGGTGGTGGTGGTGCTCGAGTTA TTTTTTTGTTTTATTTTG |
| **Pb1070*** | CTGGTGCCGCGCGGCAGCCATATGGCTAGC CAAAACACTAAAAAGCCAAAC | ATCTCAGTGGTGGTGGTGGTGGTGCTCGAGTTAC TTTACAAACTCTGCGTAAG |
| **Pb1071** | AGCAGCGGUCAAAAAACAATAGATACAATAG | ATTCGGAUGAATGATTTTTAATAGATGC |
| **Pb1072*** | CTGGTGCCGCGCGGCAGCCATATGGCTAGCACA CAAGACACTGCTAAAAAG | ATCTCAGTGGTGGTGGTGGTGGTGCTCGAGTTAA ATTAATTTTCGTTTTGA |
| **Pb1075** | AGCAGCGGUCAAGAAACTGATGACAACAC | ATTCGGAUTTATTTTGGTATTACTGC |
| **Pb1076** | AGCAGCGGUCAAGAAAAATTACCGTATC | ATTCGGAUTGTTAACATACGCATCTC |
| **Pb1077** | AGCAGCGGUCAAGAAAAATTACCGTATC | ATTCGGAUGGCTTTTGAGAAATGG |
| **Pb1078*** | CTGGTGCCGCGCGGCAGCCATATGGCTAGCAAA ACTGAAAAAAAAGTGGAA | ATCTCAGTGGTGGTGGTGGTGGTGCTCGAGTTAA TTCGCTGGCTCAACTCT |
| **Pb1081** | AGCAGCGGUCAAAATAAATTAACCGATAAAG | ATTCGGAUTTAATTTACTTTACCGTTTGC |
| **Pb1082** | AGCAGCGGUCAAGGAATTATTGAAAATTCG | ATTCGGAUGCTTCAACCATTAGAAAAATAG |
| **Pb1086** | AGCAGCGGUCAAAACTCATTTACCACAAG | ATTCGGAUTTATTCTACAGCGTGCAC |
| **Pb1091** | AGCAGCGGUATGATACTAAGTGATATAAATATTC | ATTCGGAUTTAATGAATCATATCAAAG |
| **Pb1093** | AGCAGCGGUTGTAATACTCAAGGACAAAC | ATTCGGAUCTATTTGTATTTTTTTAGACTC |
| **Pb1094** | AGCAGCGGUCAAGATAGGTTACTCTCTAAAG | ATTCGGAUCCATTTTATAAATGCAAG |
| **Pb1095** | AGCAGCGGUTGCTTTTCATCGAAGTC | ATTCGGAUCGGGTTTTGGTTTAGC |
| **Pb1096** | AGCAGCGGUTGTTCTAGTGCTAAACC | ATTCGGAUTTAAGATTTACTATCCCAATAC |
| **Pb1097*** | CTGGTGCCGCGCGGCAGCCATATGGCTAGCCAA AAAACACAAGTCACTCAA | ATCTCAGTGGTGGTGGTGGTGGTGCTCGAGTTAA TTTAATCTAGTATTTAA |
| **Pb1098** | AGCAGCGGUCAAACAAAAATAATTGCGC | ATTCGGAUCGCCTTATTTAGTTGCGT |

| | | |
|---|---|---|
| **OpSulf1** | AGCAGCGGUCAAAATAATAAAAAACCAAATATTG | ATTCGGAUCTATAGTTAATAGGAAAAAAATTG |
| **OpGHx** | AGCAGCGGUCAAGACCTATTAAATCATGTAC | ATTCGGAUTTATTTTAATGGAAGTTTTTG |

### Chemical Synthesis

#### Automated glycan assembly

Solvents were taken from an anhydrous solvent system (JC Meyer-solvent systems) to prepare activator, acid wash (TMSOTf), and capping solutions. The building blocks were co-evaporated once with toluene and dried under a high vacuum before use. All solutions were freshly prepared and kept under argon during the automation run. Final yields were calculated based on the resin loading. Resin loading was determined by performing a double glycosylation followed by DBU-promoted Fmoc-cleavage and determination of dibenzofulvene formation by measuring its UV absorbance.

#### Preparation of reagent solutions

**Building block solution**: Building block (0.09 mmol) was dissolved in DCM (1 mL).

**Fmoc deprotection solution:** Either a solution of 20% piperidine in DMF (v/v) (module E1) or a solution of 20% triethylamine in DMF (v/v) was prepared (module E2).

**Acid wash solution/phosphate donor activator:** TMSOTf (0.45 mL, 2.49 mmol) was dissolved in DCM (40 mL).

**Capping solution**: A 50 mL solution of 10% acetic anhydride and 2% methanesulfonic acid in DCM (v/v) was prepared.

**Lev deprotection solution:** A solution of hydrazine acetate (725 mg) in the mixture of pyridine (40 mL), acetic acid (10 mL) and water (2.5 mL) was prepared.

### Modules for Automated Solid-Phase Synthesis

#### Resin preparation for synthesis

The traceless linker resin (50 mg, resin loading 0.4 mmol/g) was placed in the reaction vessel and swollen in DCM for 20 min at room temperature prior to synthesis. During this time, all reagent lines needed for the synthesis were washed and primed. Before the first glycosylation, the resin was washed with the DMF, THF, and DCM (three times each with 2 mL for 25 s).

**TMSOTf acidic wash solution (Module a):** The resin was swollen in DCM (2 mL) and the temperature of the reaction vessel was adjusted to -30 °C. Upon reaching the low temperature, TMSOTf solution (1 mL, 0.06 mmol) was added dropwise to the reaction vessel. After bubbling for 3 min, the acidic solution was drained and the resin was washed with DCM (2 mL) for 25 s.

| ACTION | CYCLES | SOLUTION | AMOUNT | T (°C) | INCUBATION TIME |
|---|---|---|---|---|---|
| **COOLING** | - | - | - | -30 | - |
| **DELIVER** | 1 | DCM | 2 mL | -30 | - |
| **DELIVER** | 1 | TMSOTf solution | 1 mL | -30 | 3 min |
| **WASH** | 1 | DCM | 1 mL | -30 | 25 s |

**Phosphate glycosylation (Module b):** The building block solution (0.1 mmol of BB in 1 mL of DCM per glycosylation) was delivered to the reaction vessel. After the set temperature was reached, the reaction was started by dropwise addition of the activator solution (1.0 mL, excess). After completion of the reaction, the solution is drained and the resin was washed with DCM, DCM/dioxane (1:2, v/v, 2 mL for 20 s), and DCM (twice, each with 2 mL for 25 s). The temperature of the reaction vessel is increased to 25°C for the next module.

| ACTION | CYCLES | SOLUTION | AMOUNT | T (°C) | INCUBATION TIME |
|---|---|---|---|---|---|
| **COOLING** | - | - | - | -30 | - |
| **DELIVER** | 1 | BB solution | 1 mL | -30 | - |
| **DELIVER** | 1 | activator solution | 1 mL | -30 | - |
| **REACTION TIME** | 1 | | | -30<br>to -10 | 10 min<br>30 min |
| **WASH** | 1 | DCM | 2 mL | 0 | 25 sec |
| **WASH** | 1 | DCM:Dioxane | 2 mL | 0 | 20 sec |
| **HEATING** | - | - | - | 25 | - |
| **WASH** | 1 | DCM | 2 mL | >0 | 25 sec |

**Capping (Module c):** The resin was washed twice with DMF (2 mL, 25 s) and the temperature of the reaction vessel was adjusted to 25 °C. Pyridine solution (2 mL, 10% in DMF) was delivered into the reaction vessel. After 1 min, the reaction solution was drained and the resin was washed with DCM (three times with 3 mL for 25 s). Capping solution (4 mL) was delivered into the reaction vessel. After 20 min, the reaction solution was drained and the resin was washed with DCM (three times with 3 mL for 25 s).

| ACTION | CYCLES | SOLUTION | AMOUNT | T (°C) | INCUBATION TIME |
|---|---|---|---|---|---|
| **HEATING** | - | - | - | 25 | - |
| **WASH** | 2 | DMF | 2 mL | 25 | 25 s |
| **DELIVER** | 1 | 10% Py./DMF | 2 mL | 25 | 1 min |
| **WASH** | 3 | DCM | 2 mL | 25 | 25 s |
| **DELIVER** | 1 | Capping solution | 4 mL | 25 | 20 min |
| **WASH** | 3 | DCM | 2 mL | -20 | 25 s |

**Fmoc deprotection with $NEt_3$ (Module d):** The resin was washed with DMF (three times with 2 mL for 25 s) and the temperature of the reaction vessel was adjusted to 25 °C. Fmoc deprotection solution was delivered to the reaction vessel and kept under Ar bubbling (three times with 2 mL). After 5 min, the reaction solution was drained and the resin was washed with DMF (three times with 2 mL for 25 s) and DCM (five times each with 2 mL for 25 s). The temperature of the reaction vessel was decreased to -20 °C for the next module.

| ACTION | CYCLES | SOLUTION | AMOUNT | T (°C) | INCUBATION TIME |
|---|---|---|---|---|---|
| **WASH** | 3 | DMF | 2 mL | 25 | 25 s |
| **DELIVER** | 3 | Fmoc depr. | 2 mL | 25 | 5 min |
| **WASH** | 3 | DMF | 2 mL | 25 | 25 s |
| **WASH** | 5 | DCM | 2 mL | 25 | 25 s |
| **COOLING** | 1 | - | - | -20 | - |

**Levulinoyl ester deprotection (Module e):** The resin was washed with DCM (three times with 2 mL for 25 s) and the temperature of the reaction vessel was adjusted to 25 °C. 2 mL of Levulinoyl ester deprotection solution was delivered to the reaction vessel and kept under Ar bubbling. After 30 min, the reaction solution was drained and the resin was washed with DMF, THF and DCM (six times each with 2 mL for 25 s).

| ACTION | CYCLES | SOLUTION | AMOUNT | T (°C) | INCUBATION TIME |
|---|---|---|---|---|---|
| **HEATING** | - | - | - | 25 | - |
| **WASH** | 3 | DCM | 2 mL | 25 | 25 s |
| **DELIVER** | 1 | Lev solution | 1 mL | 25 | 30 min |
| **WASH** | 6 | DCM | 2 mL | 25 | 25 s |
| **DELIVER** | 1 | Lev solution | 1 mL | 25 | 30 min |
| **WASH** | 9 | DCM | 2 mL | 25 | 25 s |
| **DELIVER** | 1 | Lev solution | 1 mL | 25 | 30 min |
| **WASH** | 3 | DCM | 2 mL | -20 | 25 s |
| **WASH** | 6 | DMF | 2mL | <25 | 25 s |
| **WASH** | 6 | THF | 2mL | <25 | 25 s |
| **WASH** | 6 | DCM | 2mL | <25 | 25 s |

### Solid-phase synthesis

**Methanolysis (Module f):** The resin was suspended in anhydrous THF (4.8 mL). Then, 0.2 mL of a solution of NaOMe in MeOH (0.5 M) was added and the resin was shaken at room temperature for 16 h. The resin was then washed successively with THF, DCM, methanol, and DCM.

**Sulfation (Module g):** The resin was suspended in a silanized glass microwave vial in 1 mL DMF. Then, a solution of $Py\cdot SO_3$ (20 eq. per OH) in DMF:Py (4 mL, 80:20 v/v) was added.

The microwave vial was sealed and the temperature was raised to 50 °C with gentle stirring for 16 h. Thereafter, the reaction was cooled, and the resin was washed successively with DMF, $CH_2Cl_2$, methanol, and $CH_2Cl_2$.

**Photocleavage from the solid support (Module h):** Glycans were cleaved from the solid support using a batch-flow photoreactor. The resin-bound glycan (~40 mg) was suspended in DMF (4 mL) under the irradiation of an LED lamp (370nm), with stirring for 24 hours. The solution was separated from the resin using a fritted syringe and concentrated under vacuum.

### Solution-phase synthesis

**Hydrogenolysis (Module i1), oligosaccharides:** The crude compound was dissolved in 4 mL of THF: *t*BuOH: $H_2O$ (60:10:30). 5% Pd-C (200 mg) was added and the reaction was stirred under $H_2$ atmosphere for 12 h. The reaction was filtered through a pad of celite and washed with *t*BuOH and $H_2O$. The filtrates were concentrated *in vacuum*, and dissolved in 3.0 mL of water for RP-HPLC purification.

**Hydrogenolysis (Module i2), sulfated oligosaccharides:** The crude compound was dissolved in 4 mL of THF: *t*BuOH: $H_2O$ (50:20:30). 5% Pd-C (200 mg) was added and the reaction was stirred under $H_2$ atmosphere for 12 h. The reaction was filtered through a pad of celite and washed with *t*BuOH and $H_2O$. The filtrates were concentrated *in vacuum*, and dissolved in 3.0 mL of water for RP-HPLC purification.

### HPLC analysis and purification

Analytical traces of crude and pure compounds were collected using an analytic RP-HPLC Agilent 1200 Series (**Methods 1, 3, 5, 6, 7**). Purification of the crudes was conducted using a preparative RP-HPLC Agilent 1200 Series (**Methods 2, 4**).

**Method 1, analytic NP-HPLC:** (YMC-Diol-300 column, 150 x 4.6 mm) flow rate of 1.0 mL / min with Hex – 20% EtOAc as eluent [isocratic 20% EtOAc (5 min), linear gradient to 55% EtOAc (35 min), linear gradient to 100% EtOAc (5 min)].

**Method 2 analytic NP-HPLC:** (YMC-Diol-300 column, 150 x 4.6 mm) flow rate of 1.0 mL / min with Hex – 30% EtOAc as eluent [isocratic 30% EtOAc (5 min), linear gradient to 90% EtOAc (35 min), linear gradient to 100% EtOAc (5 min)].

**Method 3, analytic RP-HPLC (non-sulfated oligosaccharide):** (Hypercarb column, 150 x 4.6 mm, 3 µm) flow rate of 0.7 mL/min with ACN/$H_2O$ (0.1% formic acid) as eluents [isocratic 100 % $H_2O$ (0.1% formic acid) (5 min), linear gradient to 100% ACN (30 min)].

**Method 4, prep RP-HPLC (non-sulfated oligosaccharide):** (Hypercarb column, 150 x 10 mm, 5 µm), flow rate of 3.5 mL /min with $H_2O$ (0.1% formic acid) as eluents [isocratic 100 % $H_2O$ (0.1% formic acid) (5 min), linear gradient to 100% ACN (30 min)].

**Method 5 analytic RP-HPLC (sulfated oligosaccharide):** (Hypercarb column, 150 x 4.6 mm, 3 µm) flow rate of 0.7 mL/min with ACN/$H_2O$ (0.1mM $(NH_4)_2CO_3$) as eluents [isocratic 100 % $H_2O$ (0.1mM $(NH_4)_2CO_3$) (5 min), linear gradient to 100% ACN (30 min)].

**Method 6 prep RP-HPLC (sulfated oligosaccharide):** (Hypercarb column, 150 x 10 mm, 5 µm), flow rate of 3.5 mL /min with ACN/$H_2O$ (0.1mM $(NH_4)_2CO_3$) as eluents [isocratic 100 % $H_2O$ (0.1mM $(NH_4)_2CO_3$) (5 min), linear gradient to 100% ACN (30 min)].

Following purification, all products were lyophilized on a Christ Alpha 2-4 LD plus freeze dryer before characterization.

### Compound characterisation

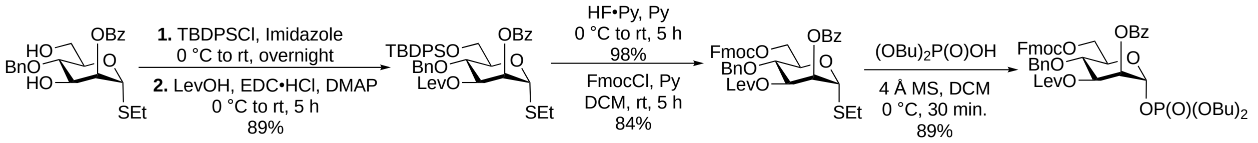


**SI Scheme 1.** Ethyl 2-*O*-benzoyl-4-*O*-benzyl-1-thio-α-D-mannopyranoside was prepared following literature procedures.[6]

#### Ethyl 2-*O*-benzoyl-4-*O*-benzyl-6-*tert*-butyl diphenylsilyl-3-*O*-levulinoyl-1-thio-α-D-mannopyranoside (4)

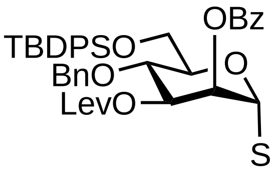


Ethyl 2-*O*-benzoyl-4-*O*-benzyl-1-thio-α-D-mannopyranoside (3.42 g, 8.18 mmol) was dissolved in anhydrous DMF (25 mL). Subsequently, imidazole (1.67 mL, 24.55 mmol, 3 eq.) and TBDPSCl (6.37 mL, 24.55 mmol, 3 eq.) were added at room temperature. The reaction was monitored by TLC and once completed was quenched by the addition of 10 % aqueous citric acid solution. The aqueous layers were extracted with $CH_2Cl_2$, and then the combined organic layers were dried over $Na_2SO_4$, filtered, and concentrated *in vacuo*. The crude material was passed through silica plug, concentrated and dried under vacuum. The intermediate was dissolved in $CH_2Cl_2$ (30 mL) and 4-dimethylaminopyridine (199 mg, 1.64 mmol, 0.2 eq.), levulinic acid (1.67 mL, 16.36 mmol, 2 eq.) and EDC•HCl (3.14 g, 16.36 mmol, 2 eq.) were added. Once the reaction reached completion (16 h), the organic layer was extracted with saturated aqueous $NaHCO_3$ and brine. The organic layer was dried over $Na_2SO_4$, filtered and concentrated. TLC. The material was purified by flash chromatography ($SiO_2$, Hexane/EtOAc) to give compound **4** (5.49 g, 89% over two steps) as a white solid.

**$^1$H NMR** (400 MHz, $CDCl_3$) δ 8.20 – 8.12 (m, 2H), 7.78 (ddt, *J* = 17.3, 6.7, 1.5 Hz, 4H), 7.69 – 7.58 (m, 1H), 7.50 – 7.43 (m, 4H), 7.38 (ddd, *J* = 8.2, 6.6, 2.9 Hz, 4H), 7.34 – 7.24 (m, 5H), 5.69 – 5.63 (m, 1H), 5.50 – 5.42 (m, 2H), 4.82 – 4.68 (m, 2H), 4.34 (t, *J* = 9.7 Hz, 1H), 4.23 (ddd, *J* = 9.7, 3.5, 1.7 Hz, 1H), 4.12 (dd, *J* = 11.4, 3.4 Hz, 1H), 3.96 (dd, *J* = 11.4, 1.7 Hz, 1H), 2.84 – 2.40 (m, 6H), 2.13 (s, 3H), 1.31 (t, *J* = 7.4 Hz, 3H), 1.16 (s, 9H). **$^{13}$C NMR** (101 MHz, $CDCl_3$) δ 206.3, 171.8, 165.6, 138.1, 136.0, 135.6, 135.6, 133.5, 133.4, 133.0, 130.0, 129.8, 129.7, 129.7, 128.6, 128.5, 128.4, 127.9, 127.8, 127.8, 127.7, 127.6, 81.9, 77.4, 77.3, 77.1, 76.8, 75.1, 73.1, 73.0, 72.8, 72.3, 62.6, 37.9, 29.8, 28.0, 26.9, 25.2, 19.4, 14.8. **HRMS** QTOF-MS: calcd. $C_{43}H_{51}O_8SSi$ for $[M+H]^+$ 755.3074, found 755.3098.

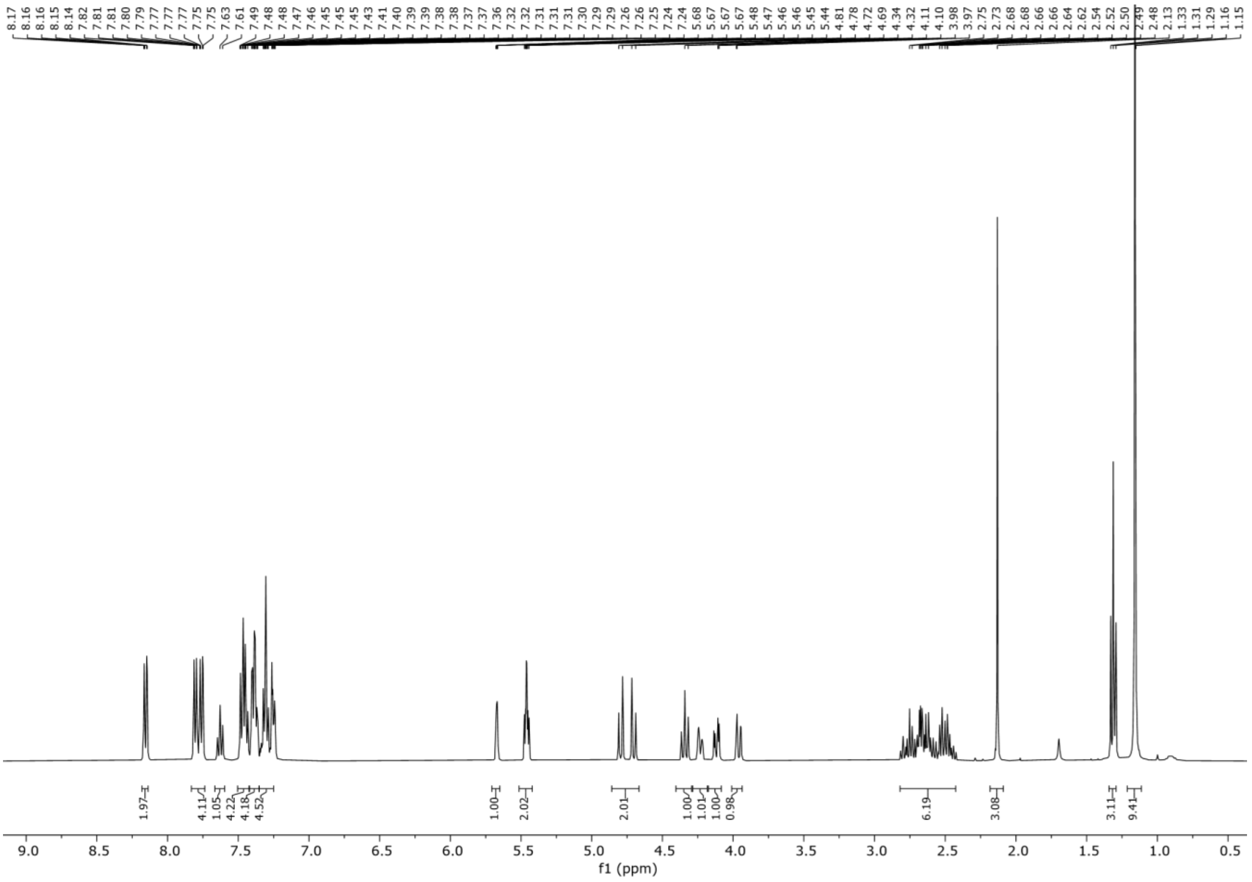
8.17 8.16 8.15 8.14 7.82 7.81 7.81 7.80 7.79 7.77 7.77 7.77 7.75 7.75 7.63 7.61 7.49 7.48 7.48 7.47 7.46 7.45 7.45 7.45 7.43 7.41 7.40 7.39 7.39 7.38 7.38 7.37 7.37 7.36 7.32 7.32 7.31 7.31 7.31 7.30 7.29 7.29 7.26 7.26 7.25 7.24 7.24 5.68 5.67 5.67 5.67 5.48 5.47 5.46 5.46 5.45 5.44 4.81 4.78 4.72 4.69 4.34 4.32 4.11 4.10 3.98 3.97 2.75 2.73 2.68 2.68 2.66 2.66 2.64 2.62 2.54 2.52 2.50 2.49 2.48 2.13 1.33 1.31 1.29 1.16 1.15
1.97 4.11 1.05 4.22 4.18 4.52 1.00 2.02 2.01 1.00 1.01 1.00 0.98 6.19 3.08 3.11 9.41
9.0 8.5 8.0 7.5 7.0 6.5 6.0 5.5 5.0 4.5 4.0 3.5 3.0 2.5 2.0 1.5 1.0 0.5
f1 (ppm)

**$^1$H NMR**

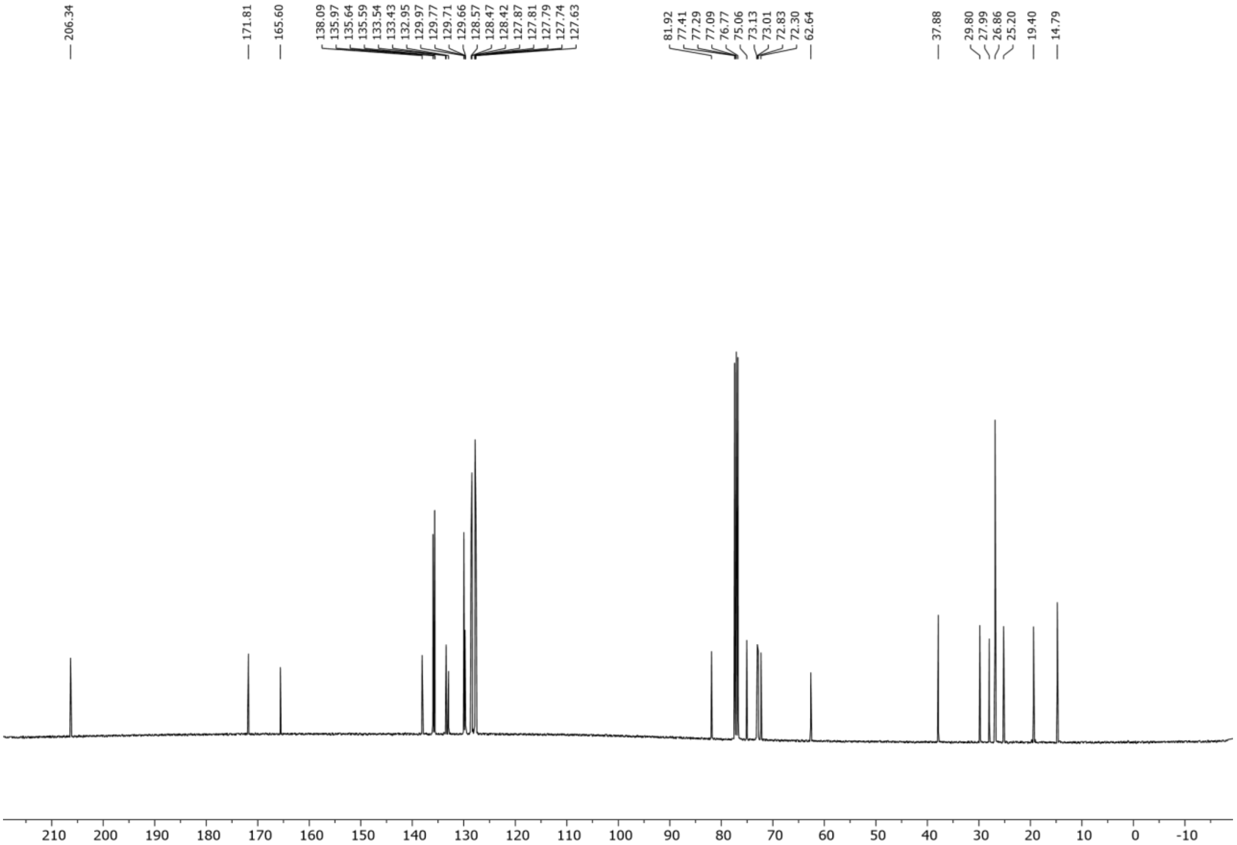
206.34
171.81
165.60
138.09 135.97 135.64 135.59 133.54 133.43 132.95 129.97 129.77 129.71 129.66 128.57 128.47 128.42 127.87 127.81 127.79 127.74 127.63
81.92 77.41 77.29 77.09 76.77 75.06 73.13 73.01 72.83 72.30 62.64
37.88
29.80 27.99 26.86 25.20 19.40 14.79
210 200 190 180 170 160 150 140 130 120 110 100 90 80 70 60 50 40 30 20 10 0 -10
f1 (ppm)

### $^{13}C$ NMR

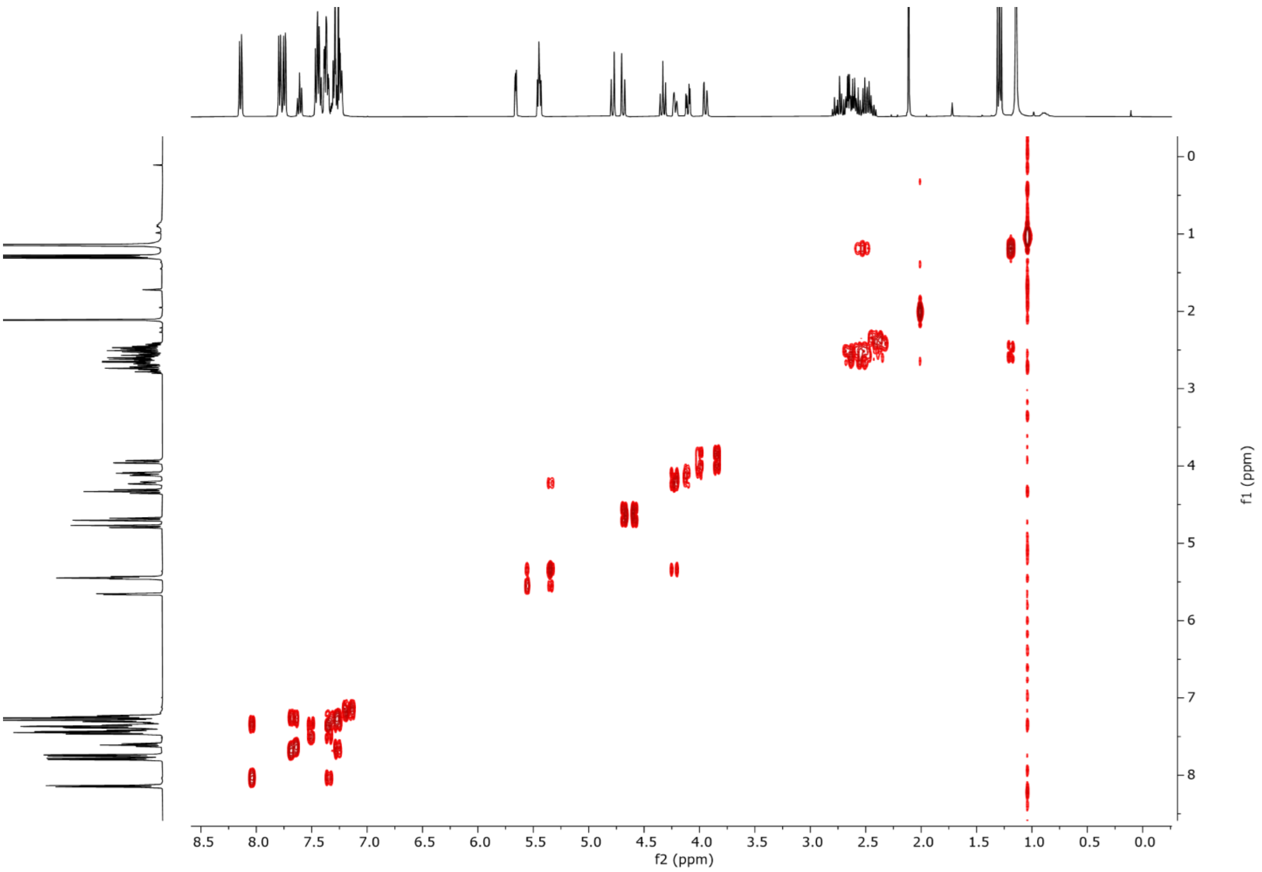


### $^{1}H$-$^{1}H$ COSY NMR

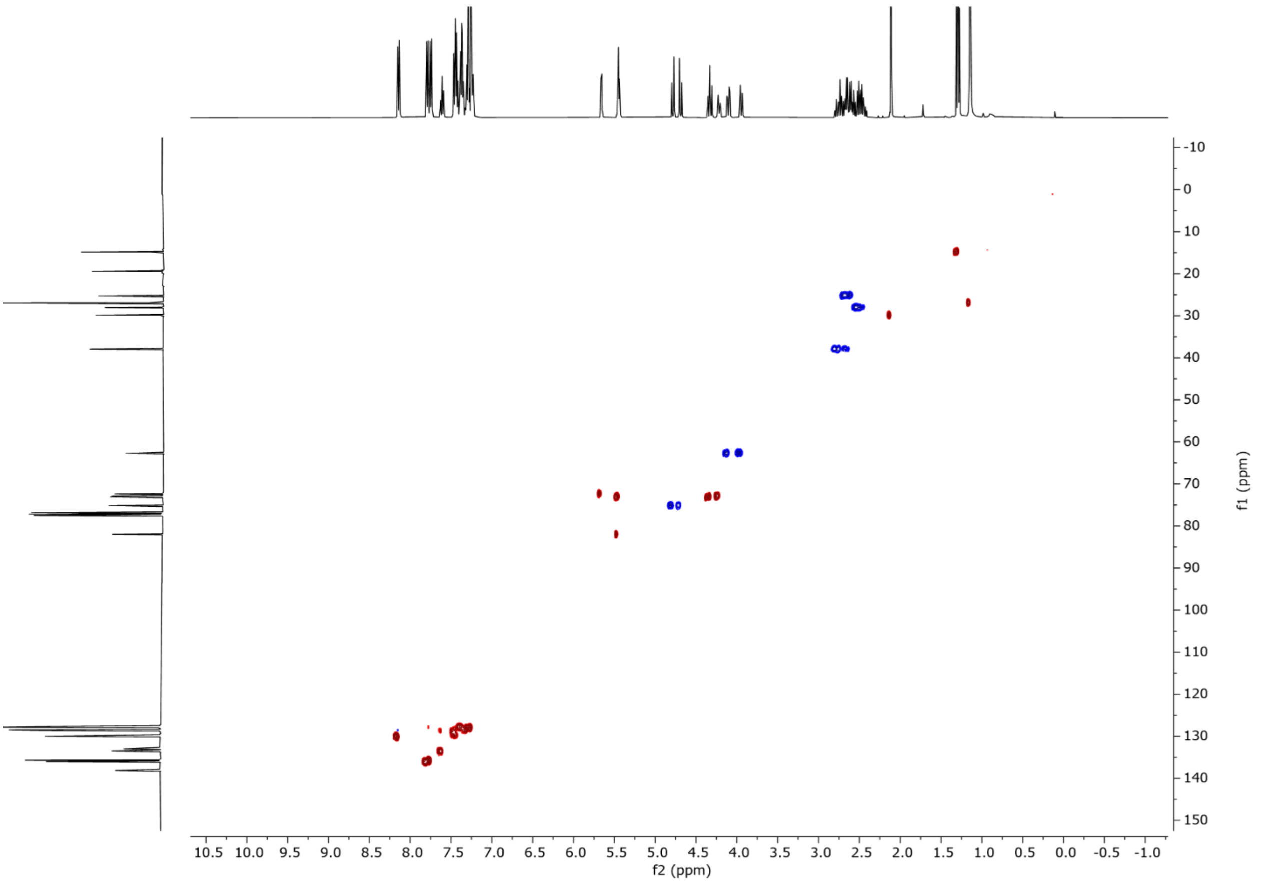

$^{1}H$-$^{13}C$ **HSQC NMR**

**Ethyl 2-*O*-benzoyl-4-*O*-benzyl-6-*O*-(9-fluorenylmethoxycarbonyl)-3-*O*-levulinoyl-1-thio-α-D-mannopyranoside (5)**

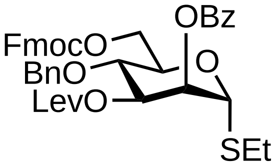


Ethyl 2-*O*-benzoyl-4-*O*-benzyl-6-*tert*-butyl diphenylsilyl-3-*O*-levulinoyl-1-thio-α-D-mannopyranoside (10.97 g, 14.53 mmol) was dissolved in pyridine (70 mL) and HF•Py (70% HF, 9.3 mL, 72.65 mmol, 5eq) was added. Once TLC analysis indicated the reaction completion (4 hours), the reaction was quenched with $Et_3N$ (2 mL). The mixture was concentrated, dissolved in ethyl acetate and subsequently washed with saturated aqueous $NaHCO_3$ and brine. The aqueous layers were extracted with EtOAc and the combined organic layers were dried over $Na_2SO_4$, filtered, and concentrated *in vacuo*. The crude was passed through silica, concentrated and dried under vacuum. The intermediate residue was dissolved in anhydrous $CH_2Cl_2$ (75 mL), pyridine (11 mL,) and fluorenylmethoxycarbonylchloride (Fmoc-Cl) (5.64 g, 21.78 mmol, 1.5 eq) was added. The reaction was tracked by TLC analysis and once completed (16h) was quenched by the addition of 10 % aqueous citric acid solution. The aqueous layers were then extracted with $CH_2Cl_2$, combined organic layers were dried over $Na_2SO_4$, filtered and concentrated *in vacuo*. The residue was purified by flash chromatography ($SiO_2$, Hexane/EtOAc) to yield compound **5** (9.00 g, 84% over two steps).

**$^1$H NMR** (400 MHz, $CDCl_3$) δ 8.14 – 8.07 (m, 2H), 7.82 – 7.75 (m, 2H), 7.68 – 7.52 (m, 3H), 7.48 – 7.37 (m, 4H), 7.35 – 7.24 (m, 7H), 5.62 (dd, $J$ = 3.2, 1.7 Hz, 1H), 5.45 – 5.37 (m, 2H), 4.77 (d, $J$ = 11.2 Hz, 1H), 4.62 (d, $J$ = 11.2 Hz, 1H), 4.49 (d, $J$ = 3.4 Hz, 2H), 4.44 – 4.39 (m, 3H), 4.28 (d, $J$ = 7.5 Hz, 1H), 4.13 – 4.08 (m, 1H), 2.83 – 2.57 (m, 4H), 2.57 – 2.37 (m, 2H), 2.11 (s, 3H), 1.29 (t, 3H). **$^{13}$C NMR** (101 MHz, $CDCl_3$) δ 206.3, 171.7, 165.4, 155.1, 143.4, 143.2, 141.3, 137.5, 133.6, 129.9, 129.5, 128.6, 128.5, 128.2, 128.0, 128.0, 127.2, 127.2, 125.2, 125.2, 120.1, 82.2, 77.4, 77.3, 77.1, 76.7, 74.9, 72.9, 72.7, 71.9, 70.1, 70.1, 66.5, 60.5, 46.8, 37.8, 29.8, 27.9, 25.5, 21.1, 14.9, 14.2. **HRMS** QTOF-MS: calcd. $C_{42}H_{42}NaO_{10}S$ for $[M+Na]^+$ 761.2396, found 761.2390.

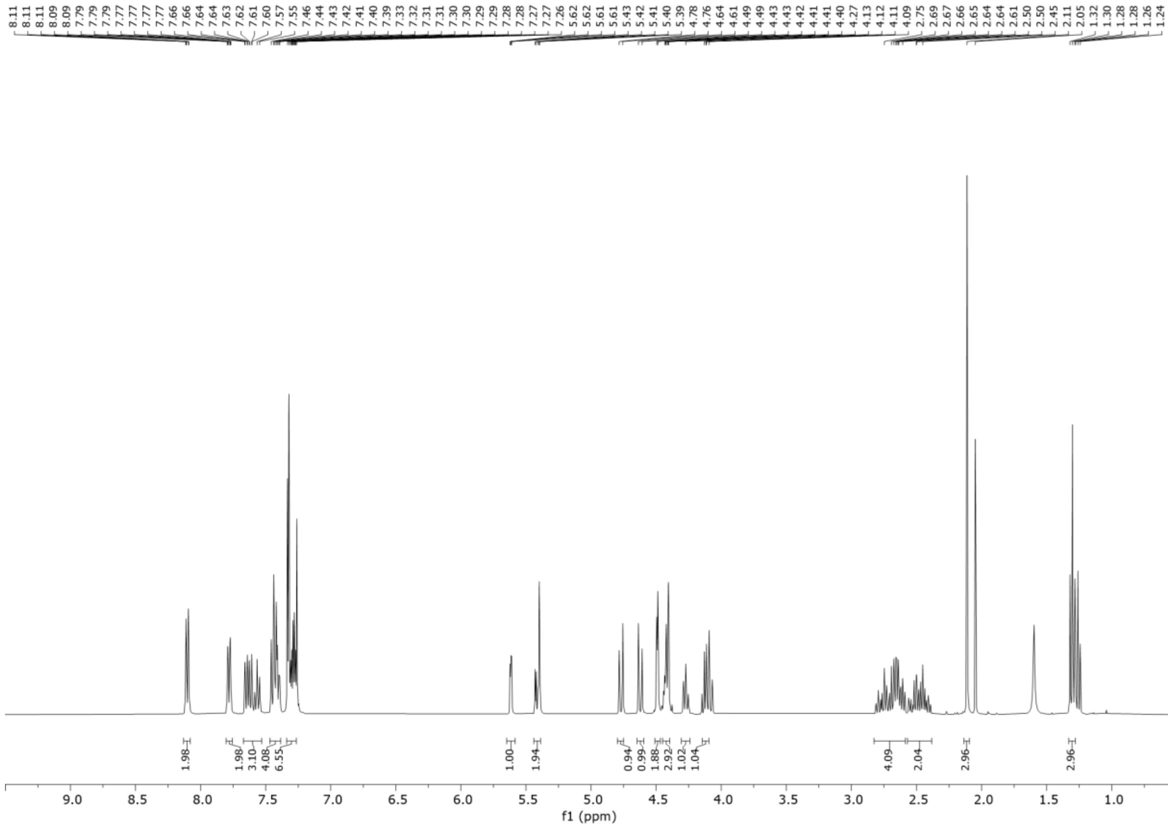

507

508 **[1]H NMR**

509

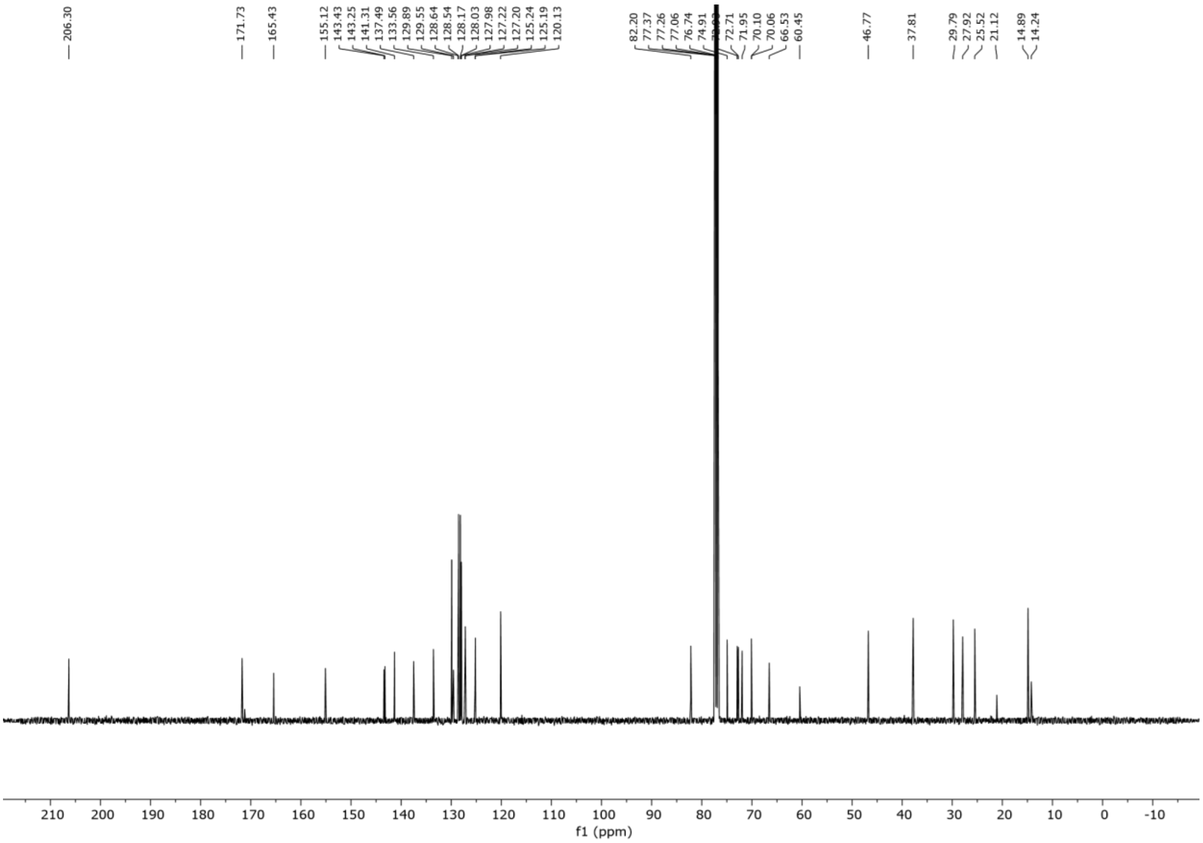
206.30
171.73
165.43
155.12
143.43
143.25
141.31
137.49
133.56
129.89
129.55
128.64
128.54
128.17
128.03
127.98
127.22
127.20
125.24
125.19
120.13
82.20
77.37
77.26
77.06
76.74
74.91
72.71
71.95
70.10
70.06
66.53
60.45
46.77
37.81
29.79
27.92
25.52
21.12
14.89
14.24
f1 (ppm)

**$^{13}$C NMR**

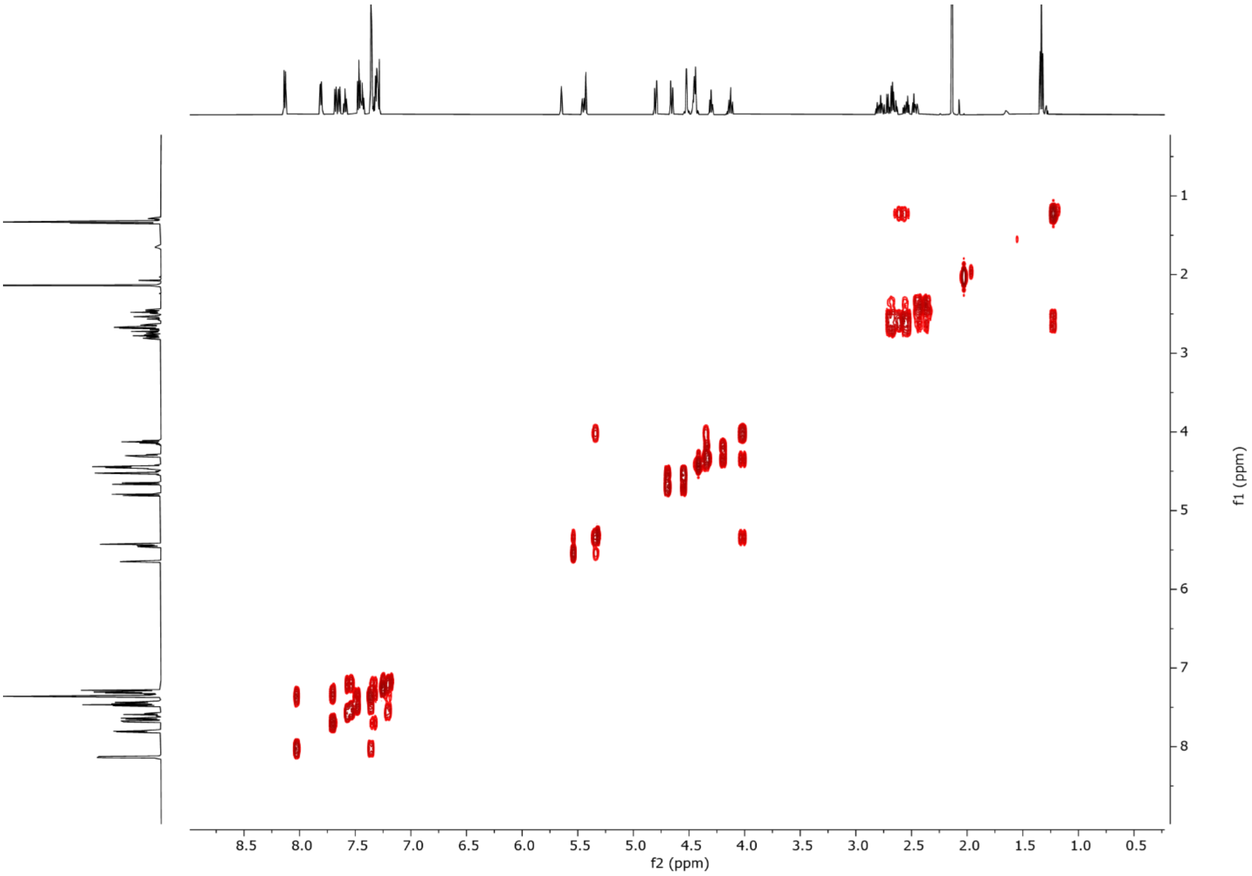
f2 (ppm)
f1 (ppm)

### $^{1}H$-$^{1}H$ COSY NMR

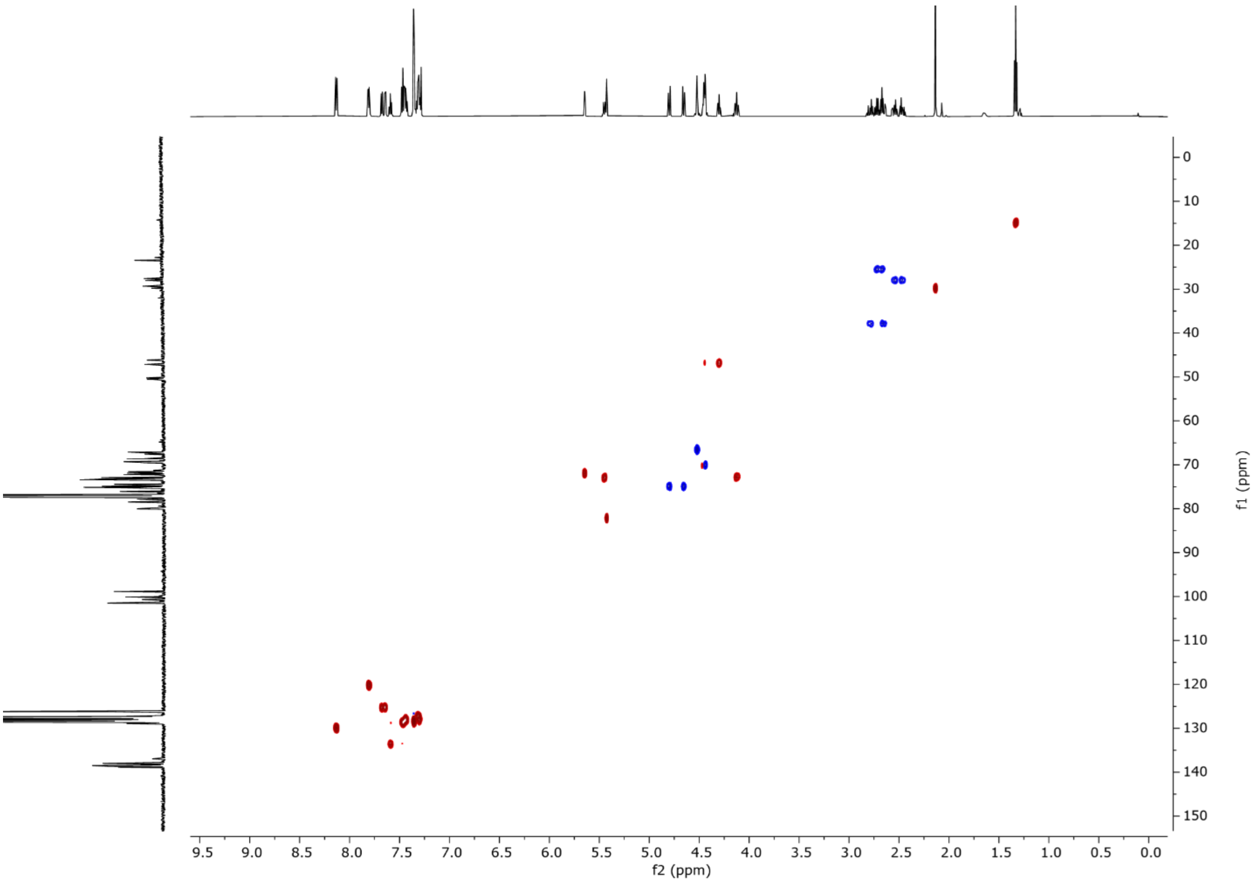


### $^{1}H$-$^{13}C$ HSQC NMR

**Dibutyl 2-*O*-benzoyl-4-*O*-benzyl-6-*O*-(9-fluorenylmethoxycarbonyl)-3-*O*-levulinoyl-1-phosphate-α-D-mannopyranoside (1)**

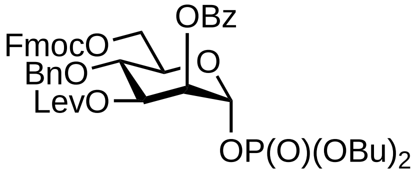


Donor **X** (2.2 g, 2.55 mmol) and dibutylphosphate (2.0 g, 3.57 mmol, 1.4 eq.) were dissolved in $CH_2Cl_2$ (20 mL) and 4Å molecular sieves were added. After 30 minutes, the mixture was cooled to 0 °C and *N*-iodosuccinimide (0.57 g, 2.55 mmol, 1 eq.) and TMSOTf (100 µL, 0.51 mmol, 0.2 eq.) were added. Once TLC analysis indicated complete consumption of the glycosyl donor (2 hours), the reaction was quenched with pyridine (2 mL) and diluted with DCM. After filtration through a plug of Celite®, the mixture was washed with 10 % aqueous $Na_2S2O_3$, saturated aqueous $NaHCO_3$, brine, and dried over $Na_2SO_4$ and concentrated *in vacuo*. The material was purified by flash chromatography ($SiO_2$, Hexane/EtOAc) to yield compound **X** (2.61 g, 98%).

**$^1$H NMR** (600 MHz, $CDCl_3$) δ 8.12 – 8.07 (m, 2H), 7.78 (dd, *J* = 7.6, 3.1 Hz, 2H), 7.63 (dd, *J* = 13.9, 7.5 Hz, 2H), 7.58 – 7.54 (m, 1H), 7.46 – 7.38 (m, 4H), 7.33 – 7.25 (m, 7H), 5.78 (dd, *J* = 6.7, 2.2 Hz, 1H), 5.59 (t, *J* = 2.7 Hz, 1H), 5.51 (dd, *J* = 9.5, 3.2 Hz, 1H), 4.78 (d, *J* = 11.1 Hz, 1H), 4.62 (d, *J* = 11.1 Hz, 1H), 4.52 – 4.38 (m, 4H), 4.30 – 4.19 (m, 2H), 4.12 (th, *J* = 8.1, 2.6 Hz, 5H), 2.83 – 2.74 (m, 1H), 2.68 – 2.60 (m, 1H), 2.57 – 2.49 (m, 1H), 2.48 – 2.40 (m, 1H), 2.12 (s, 3H), 1.73 – 1.63 (m, 4H), 1.47 – 1.36 (m, 4H), 0.93 (dt, *J* = 12.3, 7.4 Hz, 6H). **$^{13}$C NMR** (151 MHz, $CDCl_3$) δ 206.2, 171.7, 165.1, 155.0, 143.4, 143.2, 141.3, 141.3, 137.3, 133.7, 129.9, 129.1, 128.7, 128.6, 128.2, 128.1, 128.0, 127.3, 127.2, 125.2, 125.1, 120.1, 95.0, 95.0, 77.2, 77.0, 76.8, 75.1, 71.9, 71.7, 71.4, 70.1, 69.6, 69.5, 68.3, 68.2, 68.1, 68.1, 66.1, 46.8, 37.8, 32.3, 32.2, 32.2, 29.7, 27.9, 18.6, 13.6. **$^{31}$P NMR** (243 MHz, $CDCl_3$) δ -2.9. **HRMS** QTOF-MS: calcd. $C_{43}H_{49}O_{10}P$ for $[M+H]^+$ 772.3012, found 772.3012.

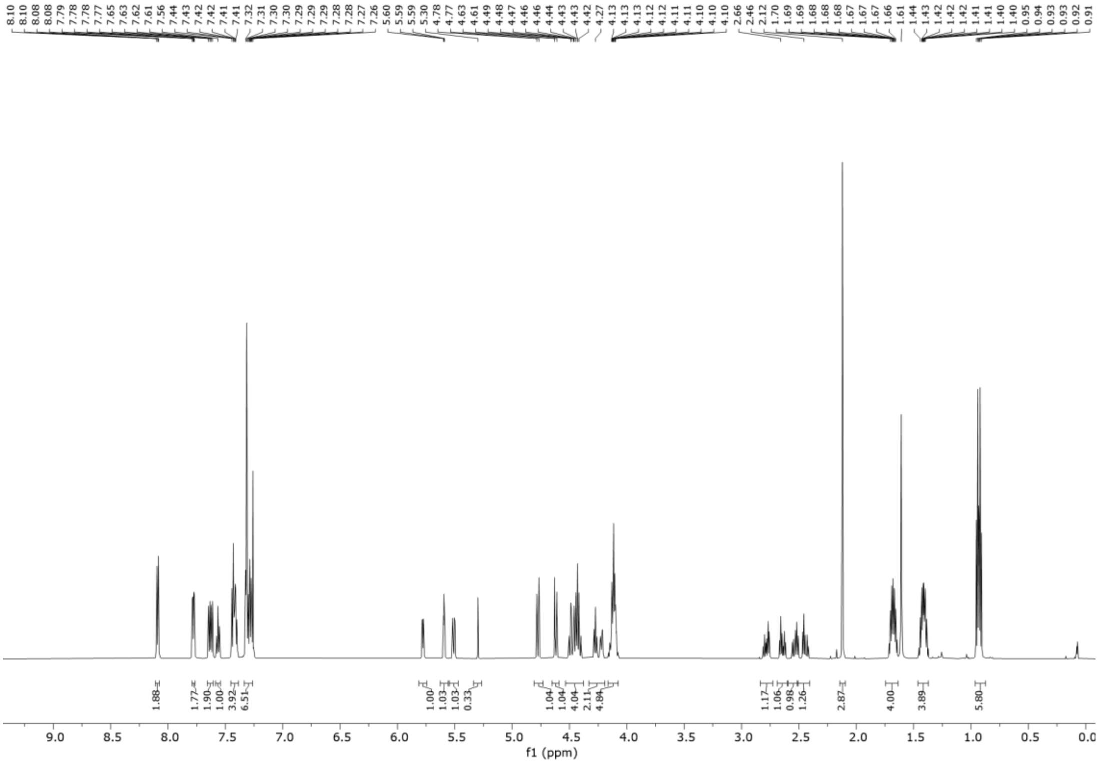


**$^1$H NMR**

547

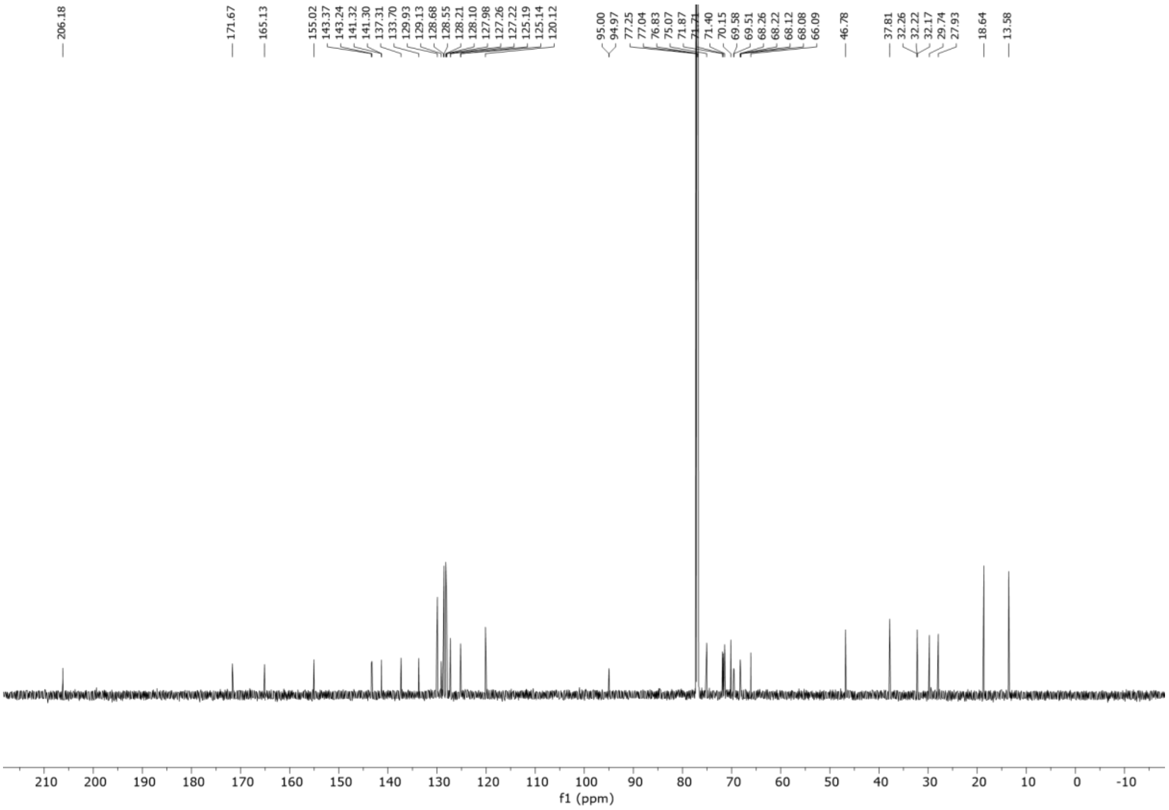
206.18
171.67
165.13
155.02
143.37
143.24
141.32
141.30
137.31
133.70
129.93
129.13
128.68
128.55
128.21
128.10
127.98
127.26
127.22
125.19
125.14
120.12
95.00
94.97
77.25
77.04
76.83
75.07
71.87
71.71
71.40
70.15
69.58
69.51
68.26
68.22
68.12
68.08
66.09
46.78
37.81
32.26
32.22
32.17
29.74
27.93
18.64
13.58
210 200 190 180 170 160 150 140 130 120 110 100 90 80 70 60 50 40 30 20 10 0 -10
f1 (ppm)

**$^{13}$C NMR**

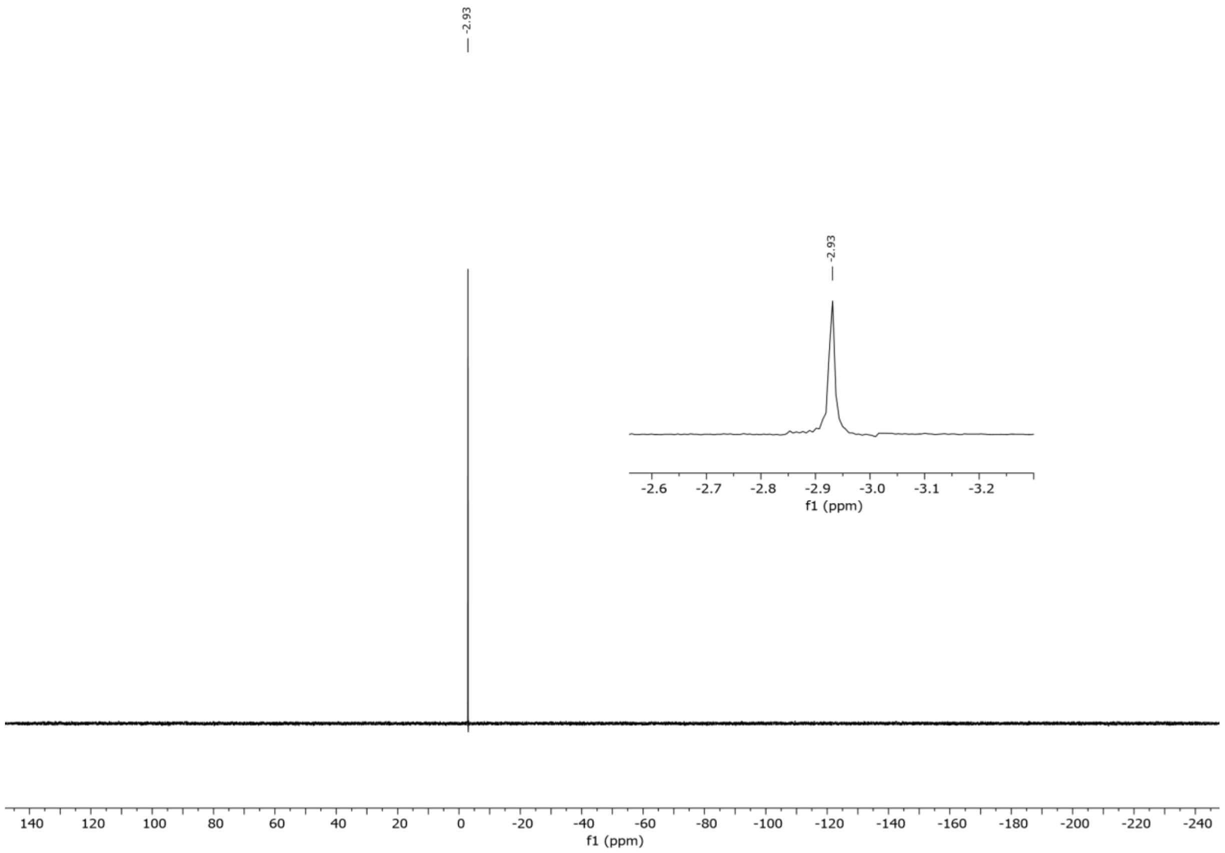
-2.93
-2.93
-2.6
-2.7
-2.8
-2.9
-3.0
-3.1
-3.2
f1 (ppm)
140
120
100
80
60
40
20
0
-20
-40
-60
-80
-100
-120
-140
-160
-180
-200
-220
-240
f1 (ppm)

**$^{31}$P NMR**

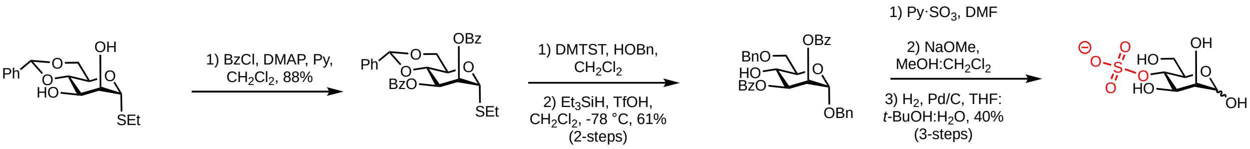


**SI Scheme X. Synthesis of 4-*O*-sulfated mannose.**

### Ethyl 4,6-*O*-benzylidene-2,3-di-*O*-benzoyl-1-thio-α-D-mannopyranoside (6)

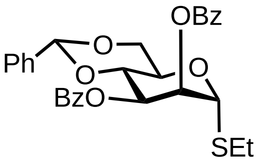


Ethyl 4,6-*O*-benzylidene-1-thio-α-D-mannopyranoside (1.5 g, 4.8 mmol, 0.2 eq.) was dissolved in $CH_2Cl_2$ (0.1 M, 20 mL) at 0 °C. Subsequently, benzoyl chloride (1.12 mL, 9.6 mmol, 2 eq.), pyridine (1.9 mL, 24.0 mmol, 5 eq.) and 4-dimethylaminopyridine (12 mg, 0.1 mmol, 0.2 eq.) were added. The mixture was stirred at room temperature under argon atmosphere. Once complete, as indicated by TLC analysis (≈16 h), the reaction was quenched with citric acid (10% w/v) and extracted. The organic layer was washed with saturated aqueous $NaHCO_3$, brine, dried over $Na_2SO_4$, and concentrated. The residue was purified by flash chromatography ($SiO_2$, Hexane/Ethyl acetate) to give compound **6** (2.2 g, 4.2 mmol, 88%) as white solid. $R_f$ = 0.91 (Hexane/Ethyl acetate, 50:50, v/v).

**$^1$H NMR** (400 MHz, $CDCl_3$) δ 8.03 – 7.95 (m, 2H), 7.87 – 7.79 (m, 2H), 7.60 – 7.13 (m, 22H), 5.78 (dd, *J* = 10.2, 3.6 Hz, 1H), 5.67 (dd, *J* = 3.6, 1.6 Hz, 1H), 5.57 (s, 1H), 5.00 (d, *J* = 1.6 Hz, 1H), 4.72 (d, *J* = 11.9 Hz, 1H), 4.61 – 4.53 (m, 3H), 4.30 – 4.20 (m, 2H), 4.14 – 3.99 (m, 1H), 3.92 – 3.83 (m, 1H). **$^{13}$C NMR** (101 MHz, $CDCl_3$) δ 192.6, 165.5, 165.5, 141.0, 137.1, 136.5, 133.6, 133.1, 130.0, 129.9, 128.7, 128.7, 128.7, 128.3, 128.2, 127.7, 127.1, 126.2, 102.0, 98.0, 71.1, 69.9, 69.1, 68.9, 65.4, 64.2. **HRMS** QTOF-MS: calcd. $C_{29}H_{30}NaO_6S$ for $[M+H]^+$ 529.1661, found 529.1668.

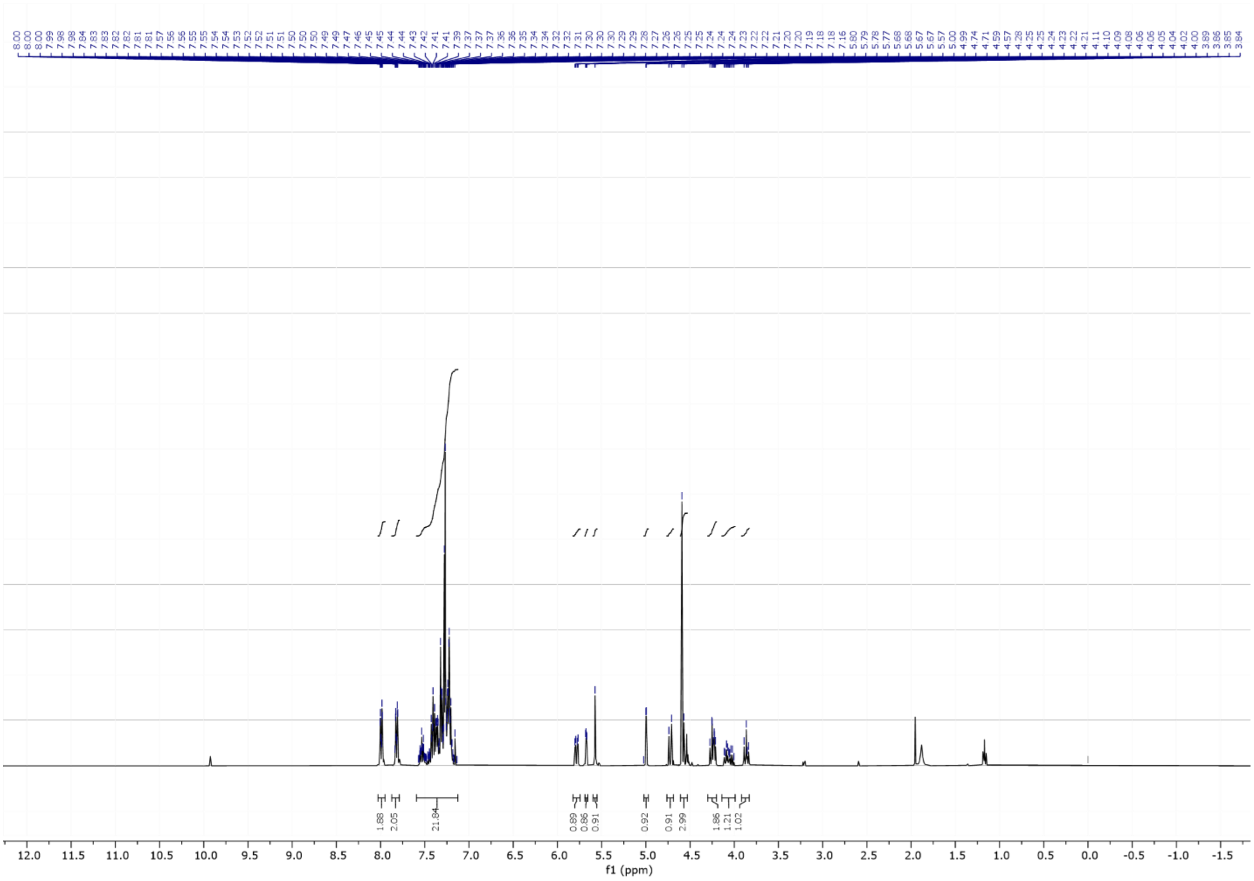
1.88
2.05
21.84
0.89
0.86
0.91
0.92
0.91
2.99
1.86
1.21
1.02
f1 (ppm)

### $^1$H NMR

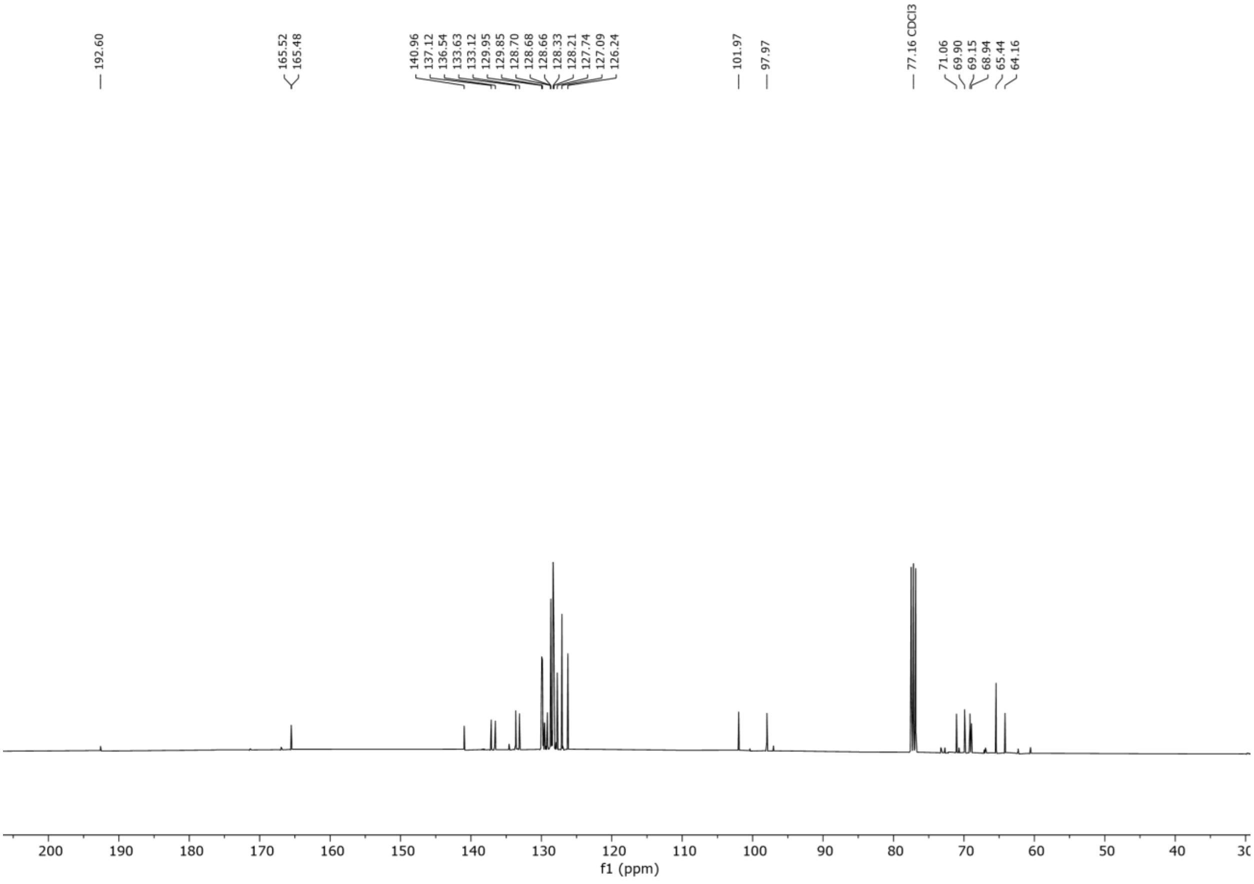
192.60
165.52
165.48
140.96
137.12
136.54
133.63
133.12
129.95
129.85
128.70
128.68
128.66
128.33
128.21
127.74
127.09
126.24
101.97
97.97
77.16 CDCl3
71.06
69.90
69.15
68.94
65.44
64.16
f1 (ppm)

**$^{13}C$ NMR**

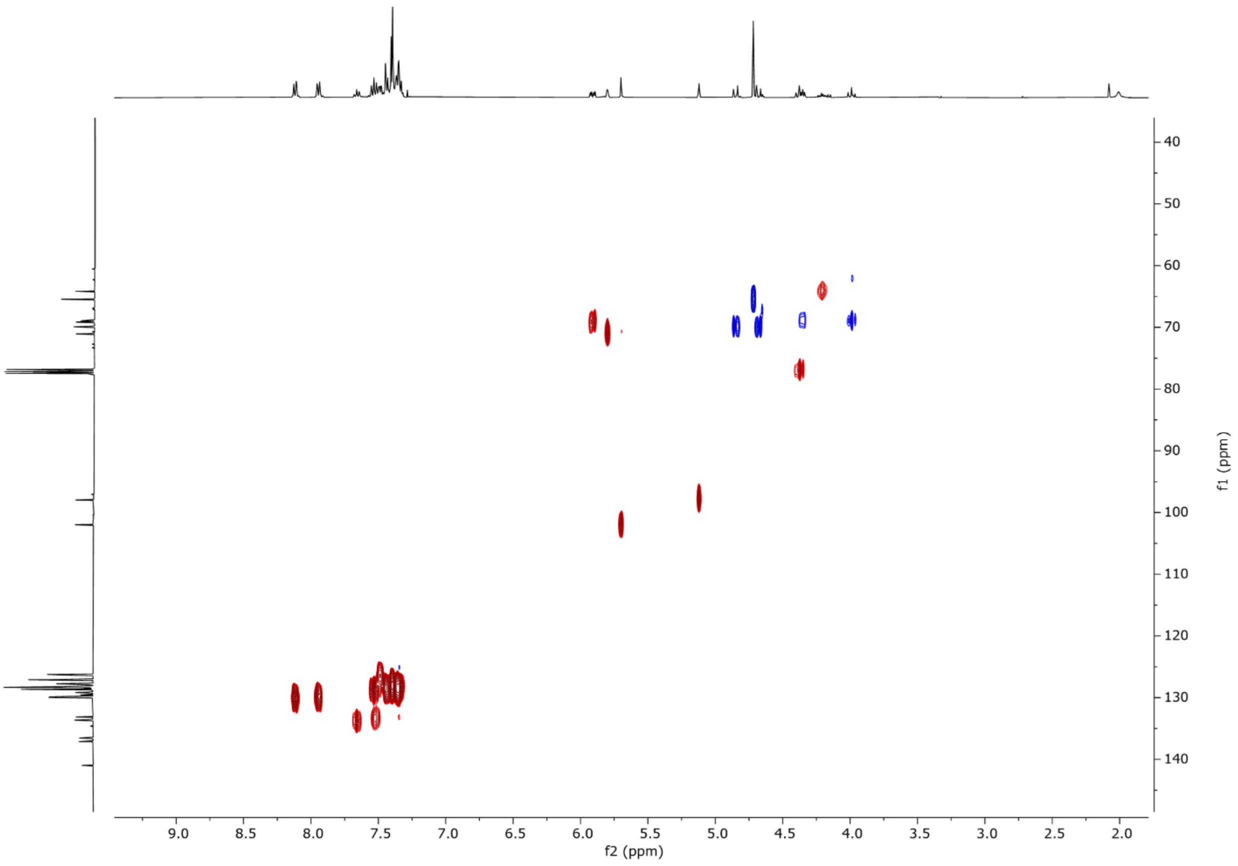


**$^{1}H$-$^{13}C$ HSQC NMR**

### **Benzyl 6-*O*-benzy-2,3-di-*O*-benzoyl-1-α-D-mannopyranoside (7)**

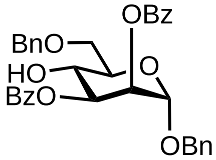


Ethyl 4,6-*O*-benzylidene-2,3-di-*O*-benzoyl-1-thio-α-D-mannopyranoside (1.5 g, 2.64 mmol) was dissolved in anhydrous $CH_2Cl_2$ (0.1M, 20 mL) and benzyl alcohol (0.55 mL, 5.28 mmol, 2 eq.) and 4Å molecular sieves were added. After 30 minutes, the mixture was cooled to 0 °C and DMTST (2.24 g, 8.71 mmol, 3.3 eq.) was added. Once TLC analysis indicated the reaction had reached completion it was quenched with triethylamine (2 mL), diluted with DCM, and filtered through a plug of Celite®. The solution was washed with saturated aqueous $NaHCO_3$, brine, and dried over $Na_2SO_4$ and concentrated. The material was purified by flash chromatography ($SiO_2$, Hexane/EtOAc) to yield the intermediate benzyl 4,6-*O*-benzylidene-2,3-di-*O*-benzoyl-1-α-D-mannopyranoside.

The intermediate compound was dissolved in dissolved in anhydrous $CH_2Cl_2$ (0.2 M, 208 mL) and 4Å molecular sieves were added. The mixture was stirred at room temperature under an argon atmosphere for 20 minutes. Subsequently, the mixture was cooled to -78 °C, and triethylsilane (1.26 mL, 7.92 mmol, 3 eq.) and triflic acid (0.7 mL, 7.92 mmol, 3 eq.) was added. The reaction was maintained at -78 °C until the complete, as indicated by TLC analysis (20 minutes). The reaction was neutralized with $Et_3N$ (1 mL) and the mixture was filtered through a pad of Celite®. The filtrate was washed with saturated aqueous $NaHCO_3$, brine, dried over $Na_2SO_4$, and concentrated. The residue was purified by flash chromatography ($SiO_2$, Hexane/Ethyl acetate) to give compound **7** (0.91 g, 1.61 mmol, 61%) as white solid.

**$^{1}$H NMR** (400 MHz, $CDCl_3$) δ 8.14 – 8.04 (m, 2H), 8.00 – 7.91 (m, 2H), 7.68 – 7.23 (m, 19H), 5.72 – 5.66 (m, 2H), 5.14 (d, *J* = 1.5 Hz, 1H), 4.84 (d, *J* = 11.9 Hz, 1H), 4.79 – 4.62 (m, 4H), 4.48 – 4.39 (m, 1H), 4.10 – 4.01 (m, 1H), 4.01 – 3.94 (m, 1H), 3.88 (dd, *J* = 10.5, 3.6 Hz, 1H). **$^{13}$C NMR** (101 MHz, $CDCl_3$) δ 166.8, 165.6, 138.2, 136.8, 133.5, 133.4, 130.0, 129.9, 129.6, 129.5, 128.7, 128.6, 128.6, 128.5, 128.4, 128.2, 128.2, 128.1, 127.8, 127.7, 127.7, 127.1, 97.0, 77.5, 77.2, 76.8, 73.9, 73.1, 71.8, 70.6, 69.9, 69.7, 67.6. **HRMS** QTOF-MS: calcd. $C_{34}H_{32}NaO_8$ for $[M+Na]^+$ 591.1995, found 591.1989.

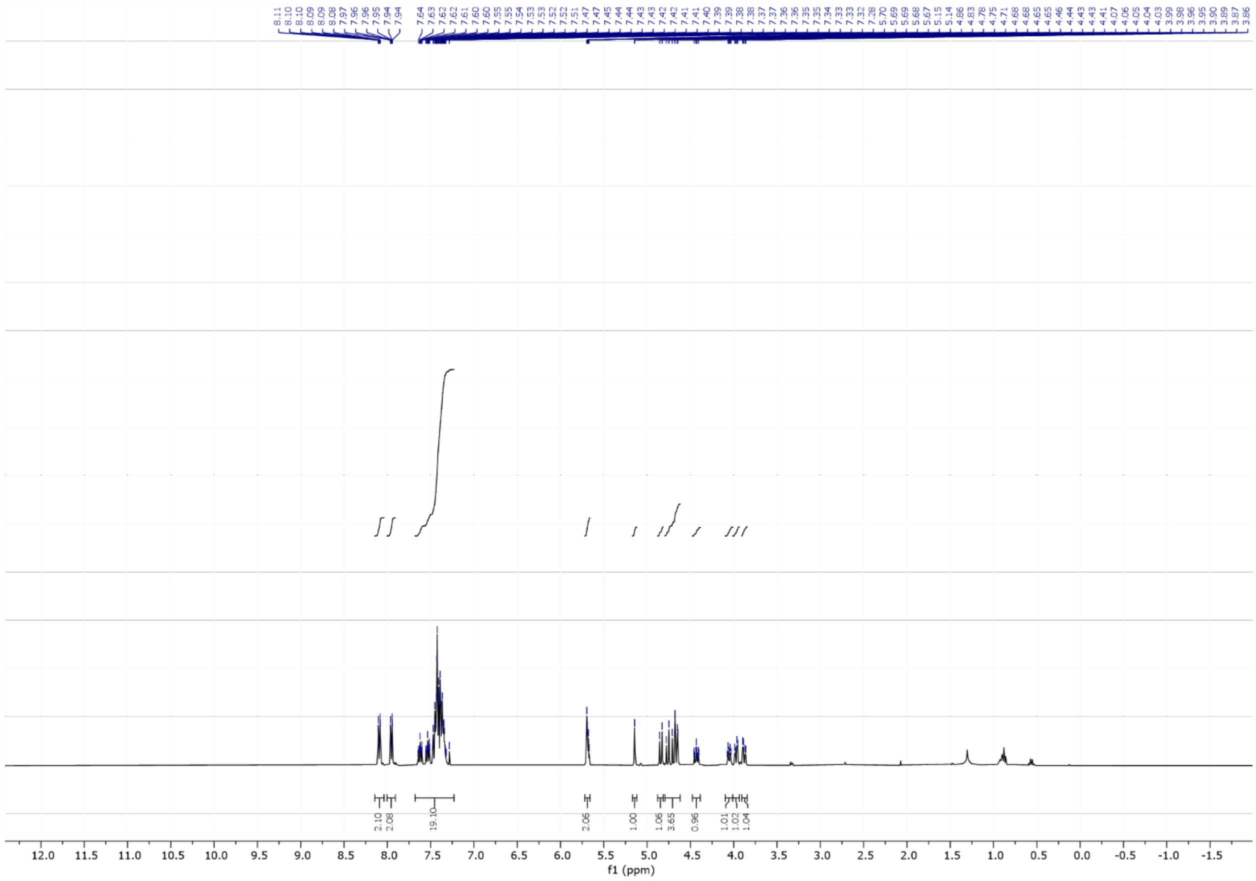


**$^{1}$H NMR**

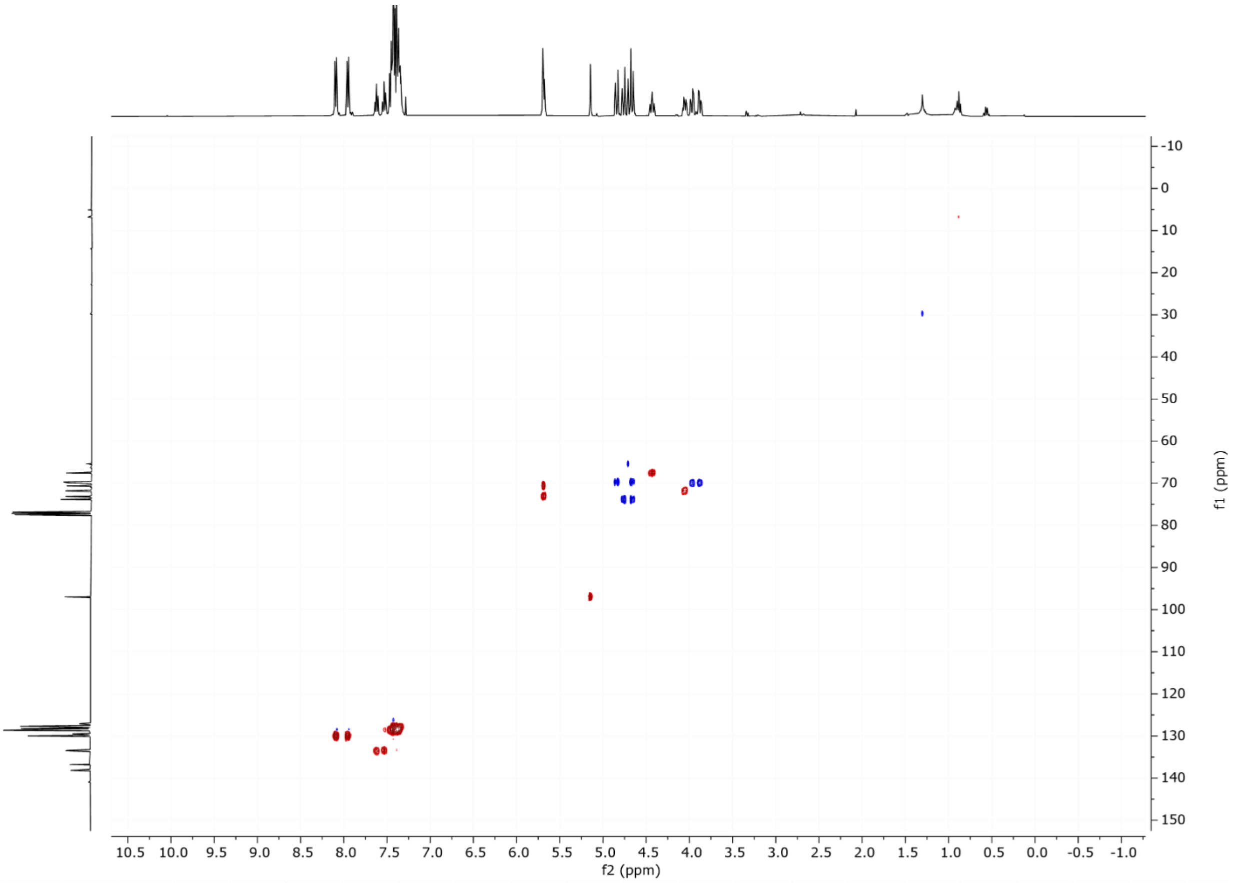
f2 (ppm)
f1 (ppm)

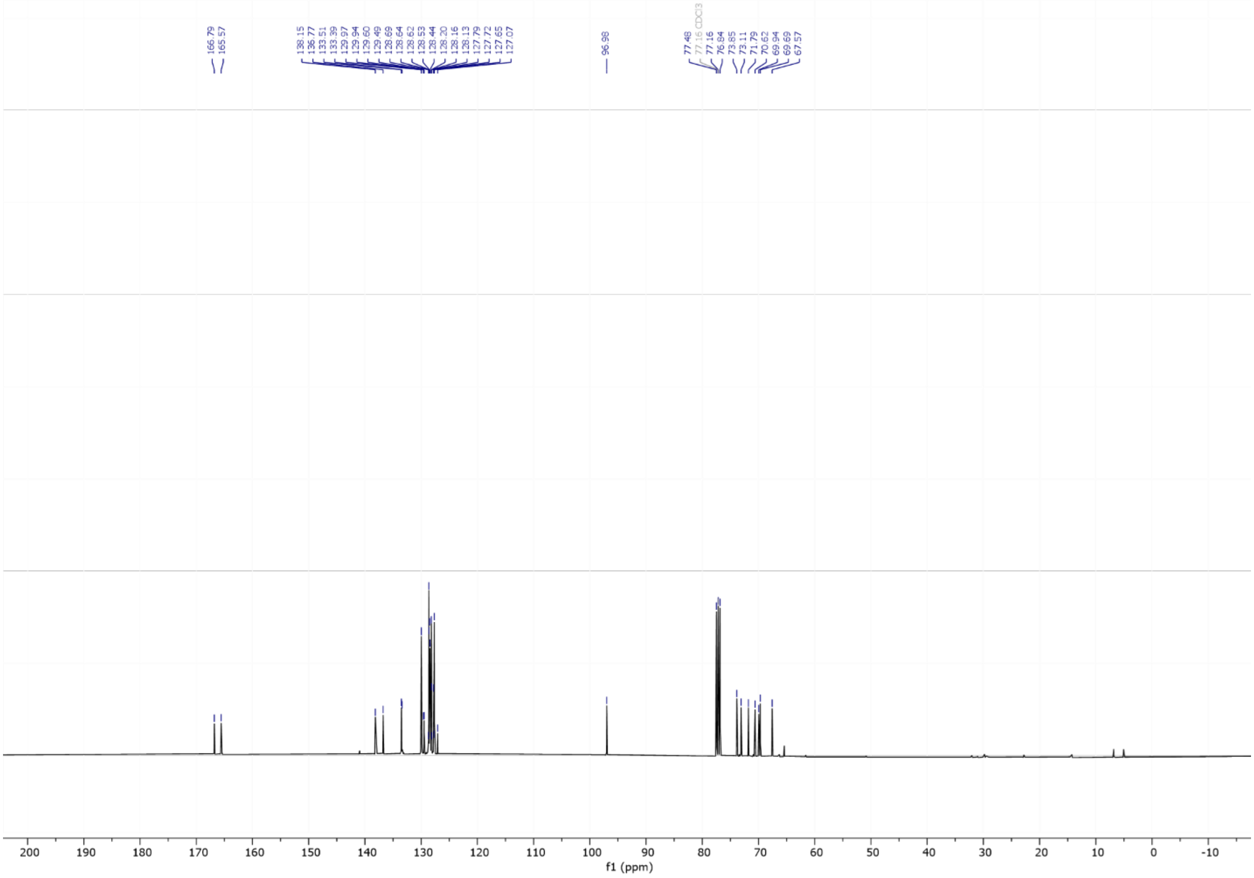
166.79
165.57
138.15
136.77
133.51
133.39
129.97
129.94
129.60
129.49
128.69
128.64
128.62
128.53
128.44
128.20
128.16
128.13
127.79
127.72
127.65
127.07
96.98
77.48
77.16 CDCl3
77.16
76.84
73.85
73.11
71.79
70.62
69.94
69.69
67.57
f1 (ppm)

### $^{13}$C NMR

### $^{1}H$-$^{13}C$ HSQC NMR

### 4-*O*-sulfonate-D-mannopyranose (8)

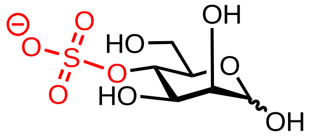


Benzyl 6-*O*-benzy-2,3-di-*O*-benzoyl-1-α-D-mannopyranoside (80 mg, 0.141 mmol) was dissolved in DMF (2 mL) and Py·$SO_3$ (140 mg, 0.70 mmole, 5 eq.) was added. The reaction was stirred at room temperature until completion, as indicated by TLC (16 h). A premixed solution of $Et_3N$/MeOH (1/1, v/v, 1.0 mL) was added to quench the reaction and stirring was continued for another 30 mins. The reaction mixture was concentrated and purified by flash chromatography ($SiO_2$, MeOH/ $CH_2Cl_2$) to give the intermediate benzyl 6-*O*-benzy-2,3-di-*O*-benzoyl-4-*O*-sulfonate-1-α-D-mannopyranoside. This intermediate was then dissolved in THF: *t*BuOH: $H_2O$ (4 mL, 50:20:30, v/v/v) and 5% Pd-C (200 mg) was added. The reaction was stirred under $H_2$ atmosphere for 12 h. The reaction was filtered through a pad of Celite® and washed with *t*BuOH and $H_2O$. The filtrates were concentrated *in vacuum*, and dissolved in 3.0 mL of water for RP-HPLC purification (Method 6) to yield **8** (16 mg, 0.063 mmol, 40%).

**$^1$H NMR** (400 MHz, $D_2O$) δ 5.12 (s, 1H), 4.86 (s, 1H), 4.32 (t, *J* = 9.5 Hz, 1H), 4.25 (t, *J* = 9.7 Hz, 0H), 3.98 (dd, *J* = 9.2, 3.3 Hz, 1H), 3.96 – 3.85 (m, 3H), 3.85 – 3.76 (m, 1H), 3.76 – 3.66 (m, 2H), 3.50 – 3.43 (m, 0H). **$^{13}$C NMR** (101 MHz, $D_2O$) δ 93.5, 75.2, 74.9, 74.4, 71.7, 71.1, 70.6, 70.6, 69.0, 60.6. **HRMS** QTOF-MS: calcd. $C_6H_{11}O_8S$ for $[M-H]^-$ 260.0202, found 260.0209.

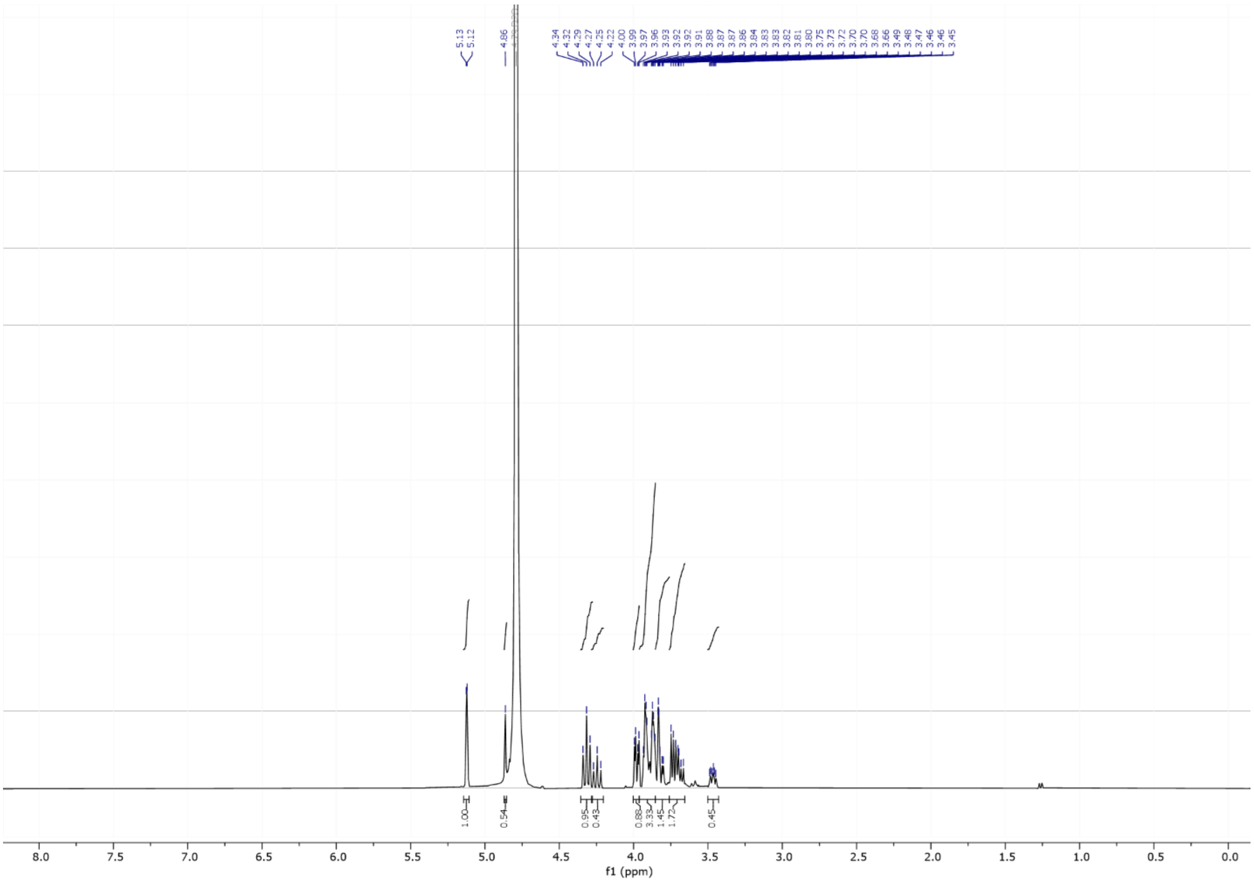


**$^1$H NMR**

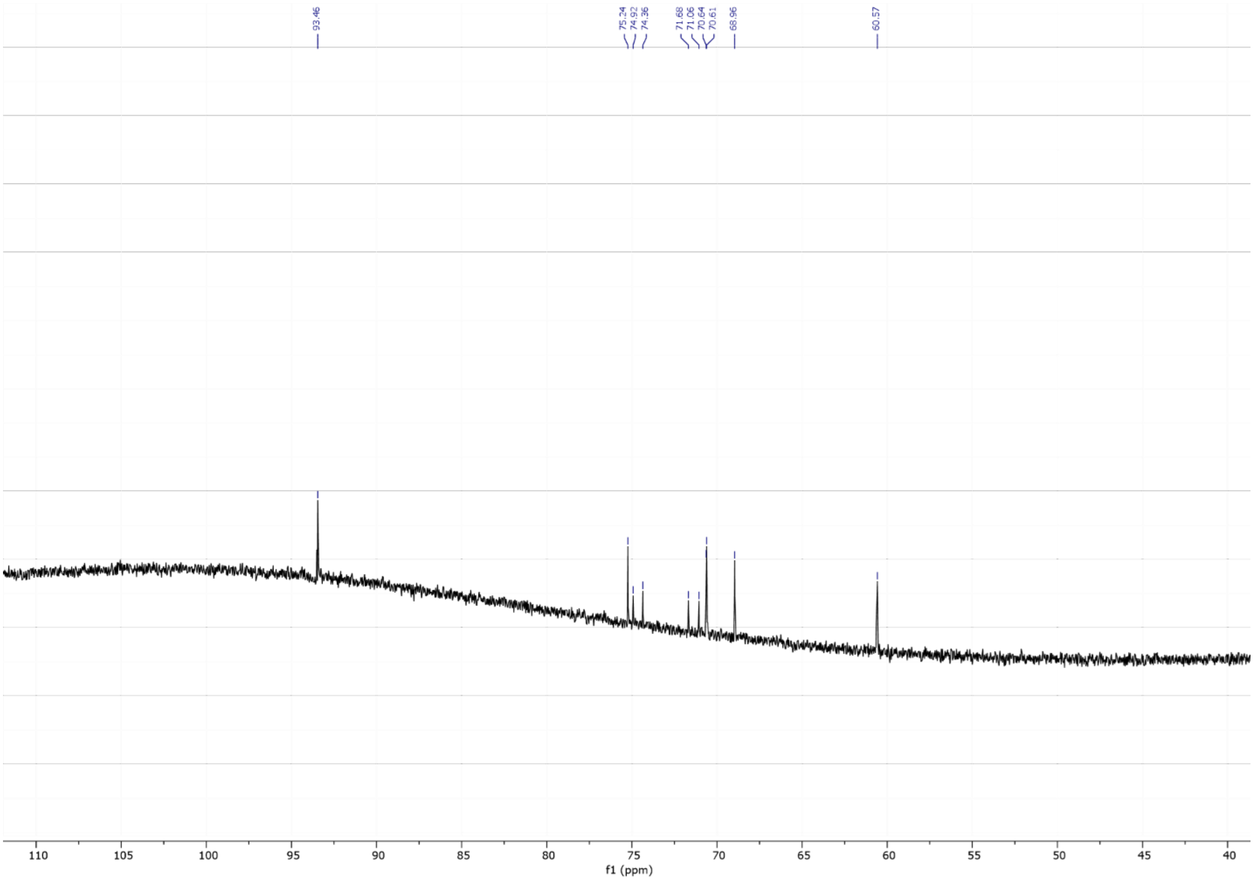
93.46
75.24
74.92
74.36
71.68
71.06
70.64
70.61
68.96
60.57
f1 (ppm)

$^{13}$C NMR

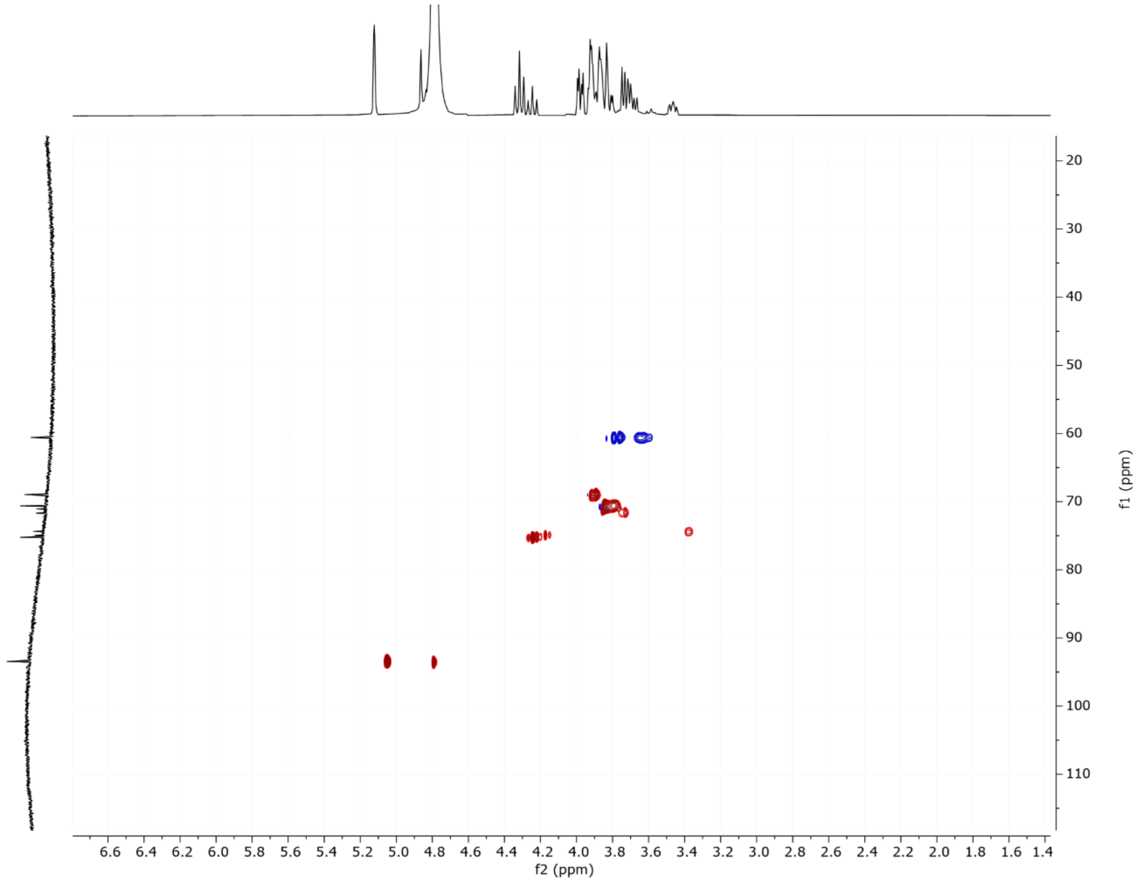
f2 (ppm)
f1 (ppm)

**$^{1}H$-$^{13}C$ HSQC NMR**

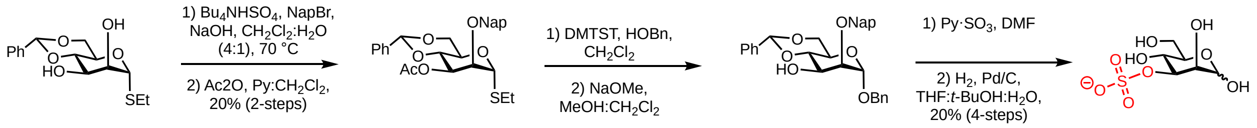


**Scheme 1. Synthesis of 3-*O*-sulfated mannose.**

### Ethyl 4,6-*O*-benzylidene-3-*O*-acetyl-2-*O*-(2-naphthalenylmethyl)-1-thio-α-D-mannopyranoside (9)

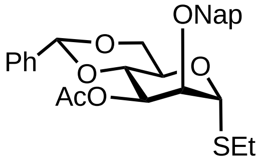


Ethyl 4,6-*O*-benzylidene-1-thio-α-D-mannopyranoside (1.0 g, 3.2 mmol) was dissolved in $CH_2Cl_2$ (50 mL) and $Bu_4NHSO_4$ (0.2 g, 0.6 mmol, 0.2 equiv.) and 2-(bromomethyl)naphthalene (0.78 g, 3.5 mmol, 1.1 eq.) were added. Subsequently, an aqueous solution of NaOH (4.0 g in 50 mL) was added and the biphasic solution was refluxed at 70 °C for 16h. The mixture was allowed to cool down to room temperature and the aqueous phase was extracted with $CH_2Cl_2$. The combined organic phase was washed with a 10% citric acid solution, dried over $Na_2SO_4$, filtered, and concentrated. The intermediate ethyl 4,6-*O*-benzylidene-2-*O*-(2-naphthalenylmethyl)-1-thio-α-D-mannopyranoside was isolated after purification by column chromatography ($SiO_2$, Hex/EtOAc).

The intermediate was dissolved in $CH_2Cl_2$ (0.1 M, 35 mL) and acetic anhydride (0.6 mL, 6.4 mmol, 2 eq.), pyridine (1.3 mL, 16.0 mmol, 5 eq.) and 4-dimethylaminopyridine (78 mg, 0.64 mmol, 0.2 eq.) were added. The mixture was stirred at room temperature. Once complete, as indicated by TLC analysis (16 h), the reaction was quenched with citric acid (10% w/v) and extracted. The organic layer was washed with saturated aqueous $NaHCO_3$, brine, dried over $Na_2SO_4$, and concentrated. The residue was purified by flash chromatography ($SiO_2$, Hexane/Ethyl acetate) to give compound **9** (0.3 g, 0.64 mmol, 20% over 2 steps) as white solid.

**$^1$H NMR** (400 MHz, $CDCl_3$) δ 7.83 – 7.69 (m, 4H), 7.50 – 7.34 (m, 5H), 7.33 – 7.21 (m, 3H), 5.51 (s, 1H), 5.30 (s, 1H), 5.17 (dd, *J* = 10.2, 3.5 Hz, 1H), 4.78 (d, *J* = 12.1 Hz, 1H), 4.66 (d, *J* = 12.1 Hz, 1H), 4.30 – 4.11 (m, 3H), 4.08 – 4.02 (m, 1H), 3.83 (t, *J* = 10.1 Hz, 1H), 3.09 (s, 2H), 2.59 – 2.45 (m, 3H), 1.90 (s, 3H), 1.16 (t, *J* = 7.4 Hz, 3H). **$^{13}$C NMR** (101 MHz, $CDCl_3$) δ 170.3, 137.4, 135.0, 133.3, 133.2, 129.2, 128.5, 128.4, 128.0, 127.8, 127.1, 126.5, 126.4, 126.3, 126.0, 101.9, 83.1, 78.0, 76.6, 73.5, 70.8, 68.7, 64.6, 25.5, 23.1, 21.1, 14.9. **HRMS**

QTOF-MS: calcd. $C_{28}H_{30}NaO_6S$ for $[M+Na]^+$ 517.1661, found 517.1654.

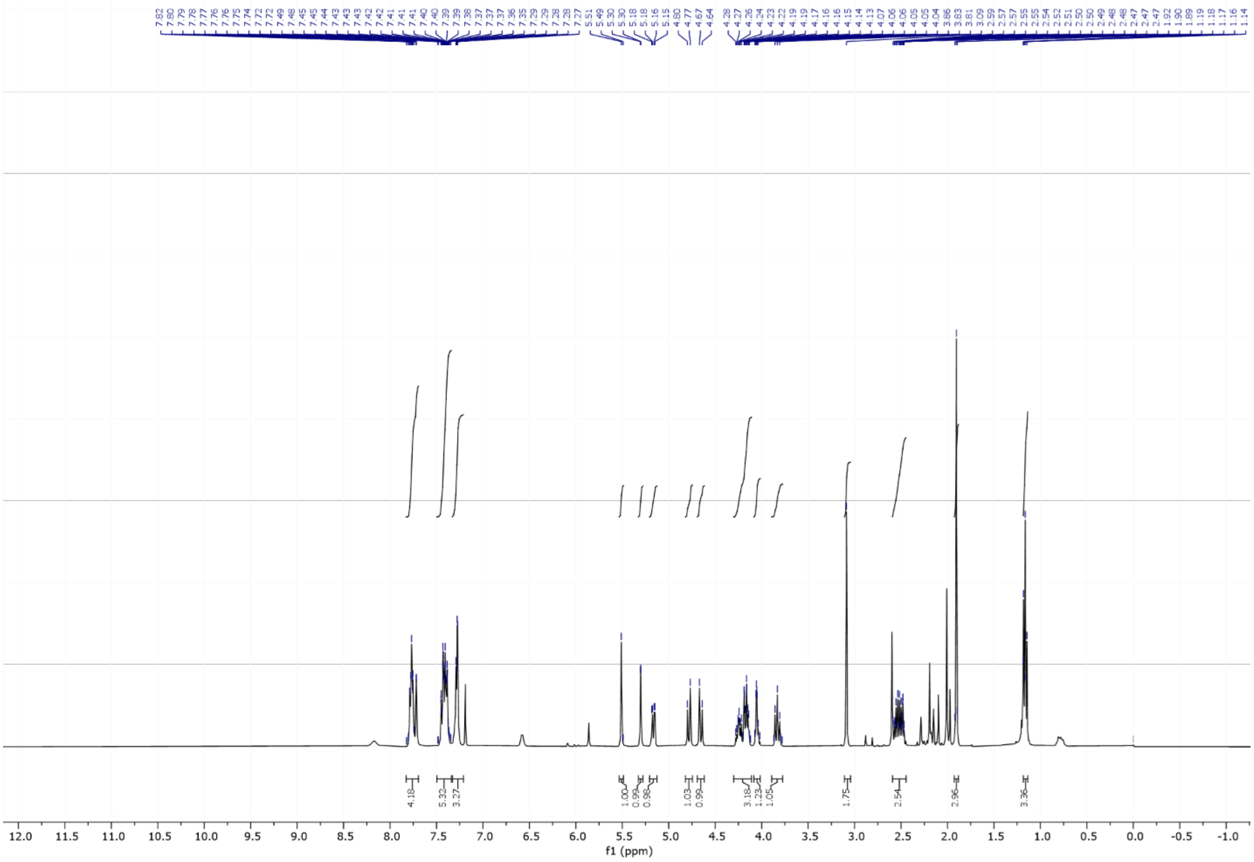


**$^1$H NMR**

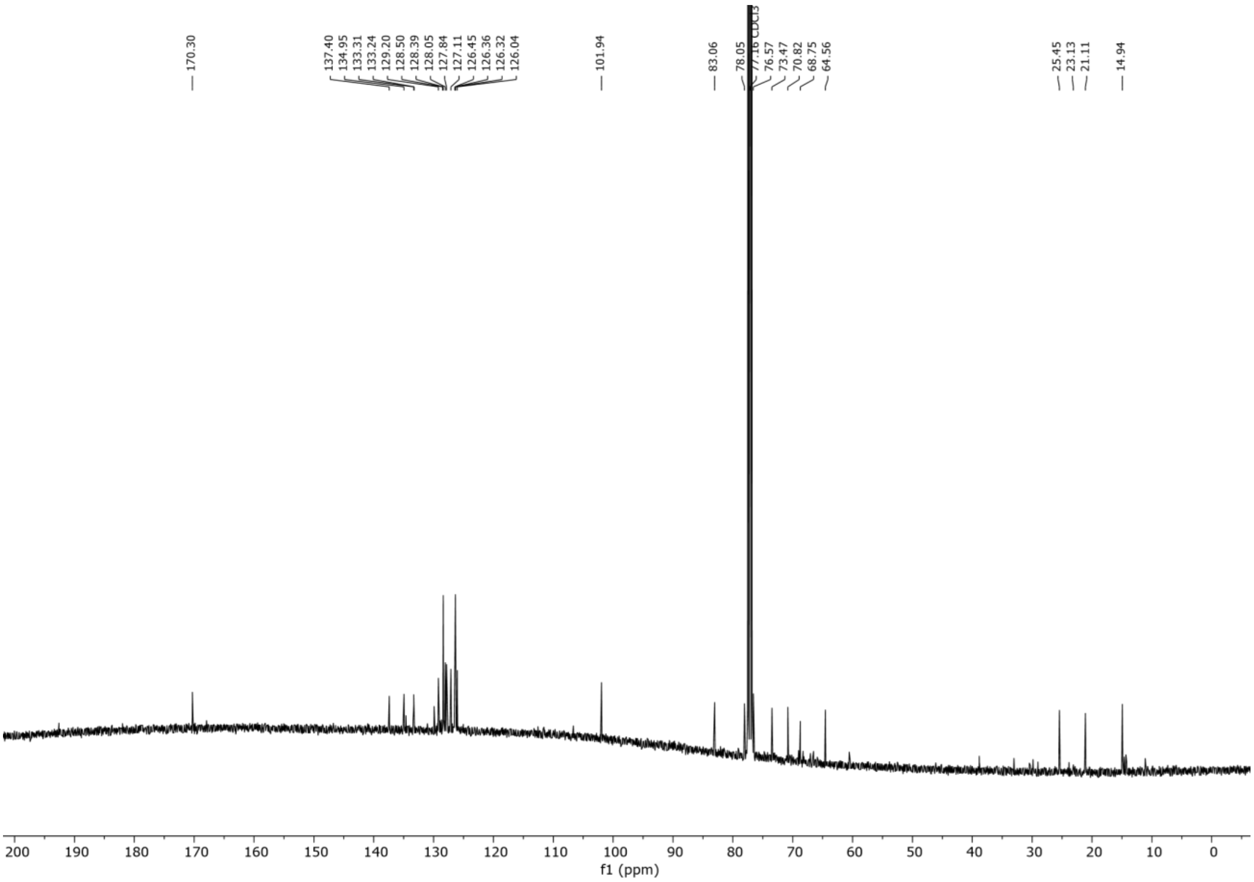

**$^{13}$C NMR**

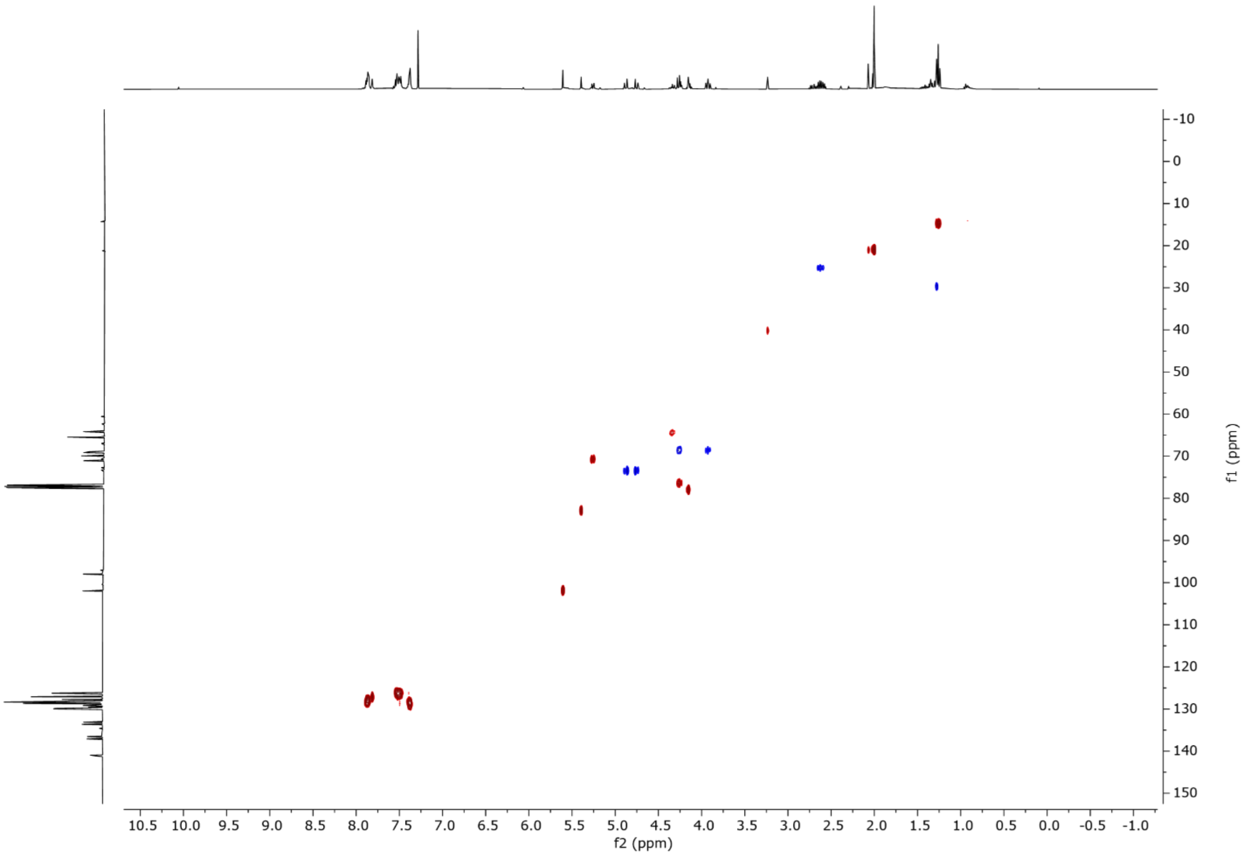


**$^{1}$H-$^{13}$C HSQC**

### 3-*O*-sulfonate-D-mannopyranose (10)

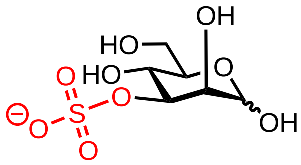

Ethyl 4,6-*O*-benzylidene-3-*O*-acetyl-2-*O*-(2-naphthalenylmethyl)-1-thio-α-D-mannopyranoside (0.3 g, 0.60 mmol) was dissolved in anhydrous $CH_2Cl_2$ (0.1M, 6 mL) and benzyl alcohol (0.18 mL, 1.82 mmol, 3eq) and 4Å molecular sieves were added. After 30 minutes, the mixture was cooled to 0 °C and DMTST (0.4 g, 1.82 mmol, 3 eq.) was added. Once TLC analysis indicated the reaction had reached completion it was quenched with triethylamine (3 mL), diluted with DCM, and filtered through a plug of Celite®. The solution was washed with saturated aqueous $NaHCO_3$, brine, and dried over $Na_2SO_4$ and concentrated. The material was purified by flash chromatography ($SiO_2$, Hexane/EtOAc) to yield the intermediate benzyl 4,6-*O*-benzylidene-3-*O*-acetyl-2-*O*-(2-naphthalenylmethyl)-1-thio-α-D-mannopyranoside. The intermediate was dissolved in $CH_2Cl_2$ (0.1M, 6 mL) and the pH was raised to pH 8 with a sodium methoxide (approx. 0.1 mL, 0.5M solution). Once complete, the reaction was quenched by the addition of Amberlite™ (Hydrogen foam), filtered, and concentrated under vacuum. The crude material was then dissolved in DMF (6 mL) and Py·$SO_3$ (5 eq) was added and stirred at room temperature. Upon completion of reaction as indicated by TLC (16 h), a premixed solution of $Et_3N$/MeOH (1/1, v/v, 1.0 mL) was added to quench the reaction and stirring was continued for another 30 mins. The reaction mixture was concentrated and purified by flash chromatography ($SiO_2$, MeOH/ $CH_2Cl_2$) to give the intermediate benzyl 4,6-*O*-benzylidene-3-*O*-sulfonate-2-*O*-(2-naphthalenylmethyl)-1-thio-α-D-mannopyranoside. This intermediate was then dissolved in THF: *t*BuOH: $H_2O$ (4 mL, 50:20:30) and 5% Pd-C (200 mg) was added. The reaction was stirred under $H_2$ atmosphere for 12 h. The reaction was filtered through a pad of celite and washed with *t*BuOH and $H_2O$. The filtrates were concentrated *in vacuum*, and dissolved in 3.0 mL of water for RP-HPLC purification (Method 6) to yield **10** (31 mg, 0.12 mmol, 20%).

**$^1$H NMR** (400 MHz, $D_2O$) δ 5.13 (d, *J* = 2.0 Hz, 1H), 4.89 (d, *J* = 1.1 Hz, 0H), 4.47 (dd, *J* = 9.5, 3.2 Hz, 1H), 4.30 – 4.26 (m, 0H), 4.21 (dd, *J* = 3.3, 2.0 Hz, 1H), 3.89 – 3.62 (m, 4H), 3.41 (ddd, *J* = 9.9, 6.0, 2.3 Hz, 0H). **$^{13}$C NMR** (101 MHz, $D_2O$) δ 93.9, 93.4, 78.9, 72.4, 69.2, 64.7, 60.8. **HRMS** QTOF-MS: calcd. $C_6H_{11}O_9S$ for $[M-H]^-$ 259.0129, found 259.0134.

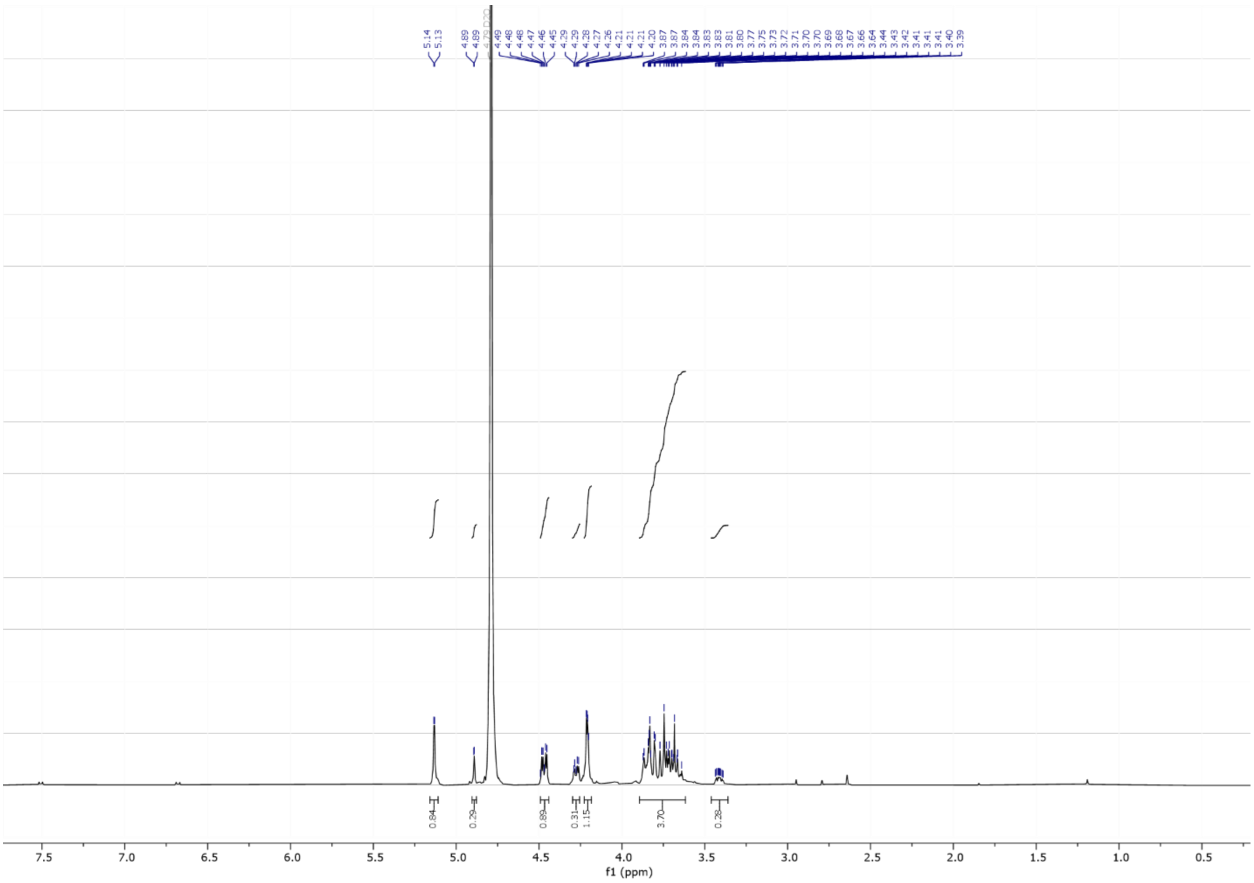


**$^{1}$H NMR**

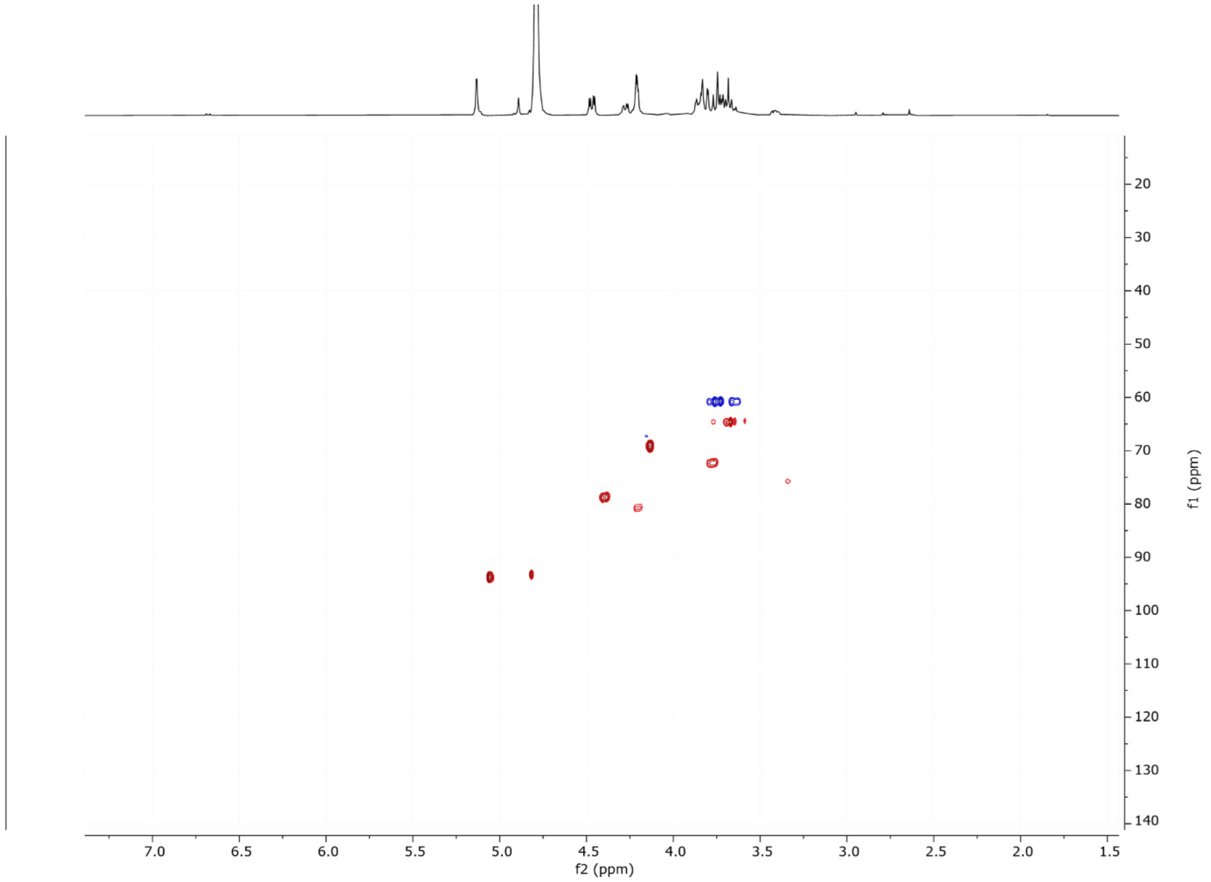


**$^{1}$H-$^{13}$C HSQC NMR**

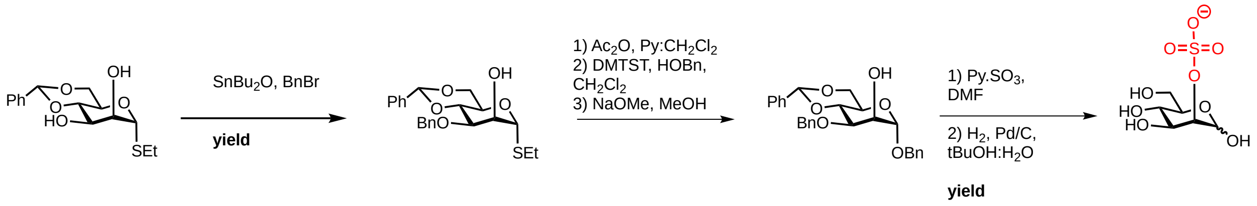


**Scheme X. Synthesis of 2-*O*-sulfated mannose.**

#### Ethyl 4,6-*O*-benzylidene-3-*O*-benzyl-1-thio-α-D-mannopyranoside (11)

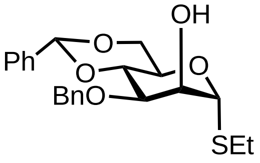


Ethyl 4,6-*O*-benzylidene-1-thio-α-D-mannopyranoside (1.5 g, 4.19 mmol) was dissolved in anhydrous toluene (0.1M, 65 mL), *n*$Bu_2SnO$ (1.25 g, 5.03 mmol, 1.2 eq.) was added, and the reaction was left to reflux overnight. Subsequently, the reaction was brought to 40 °C. Thereafter, benzyl bromide (2.8 g, 12.6 mmol, 3 eq.) and TBAI (0.3 g, 0.83 mmol, 0.2 eq.) was added. Once the reaction was complete, the solution was concentrated under vacuum and purified by flash chromatography ($SiO_2$, hexane/EtOAc) to yield **11** (1.1 g, 2.93 mmol, 70%).

**$^1$H NMR** (400 MHz, $CDCl_3$) δ 7.51 – 7.39 (m, 2H), 7.38 – 7.16 (m, 9H), 5.54 (s, 1H), 5.29 (s, 1H), 4.78 (d, *J* = 11.7 Hz, 1H), 4.62 (d, *J* = 11.9 Hz, 1H), 4.22 – 3.95 (m, 4H), 3.86 – 3.71 (m, 2H), 2.64 – 2.46 (m, 3H), 1.29 – 1.14 (m, 3H). **$^{13}$C NMR** (101 MHz, $CDCl_3$) δ 137.9, 137.6, 134.6, 129.9, 129.1, 129.1, 128.6, 128.4, 128.1, 128.0, 126.2, 101.7, 84.2, 79.2, 76.0, 73.2, 71.5, 68.8, 63.9, 25.0, 15.0. **HRMS** QTOF-MS: calcd. $C_{22}H_{27}O_5S$ for $[M+H]^+$ 403.1579, found 403.1585.

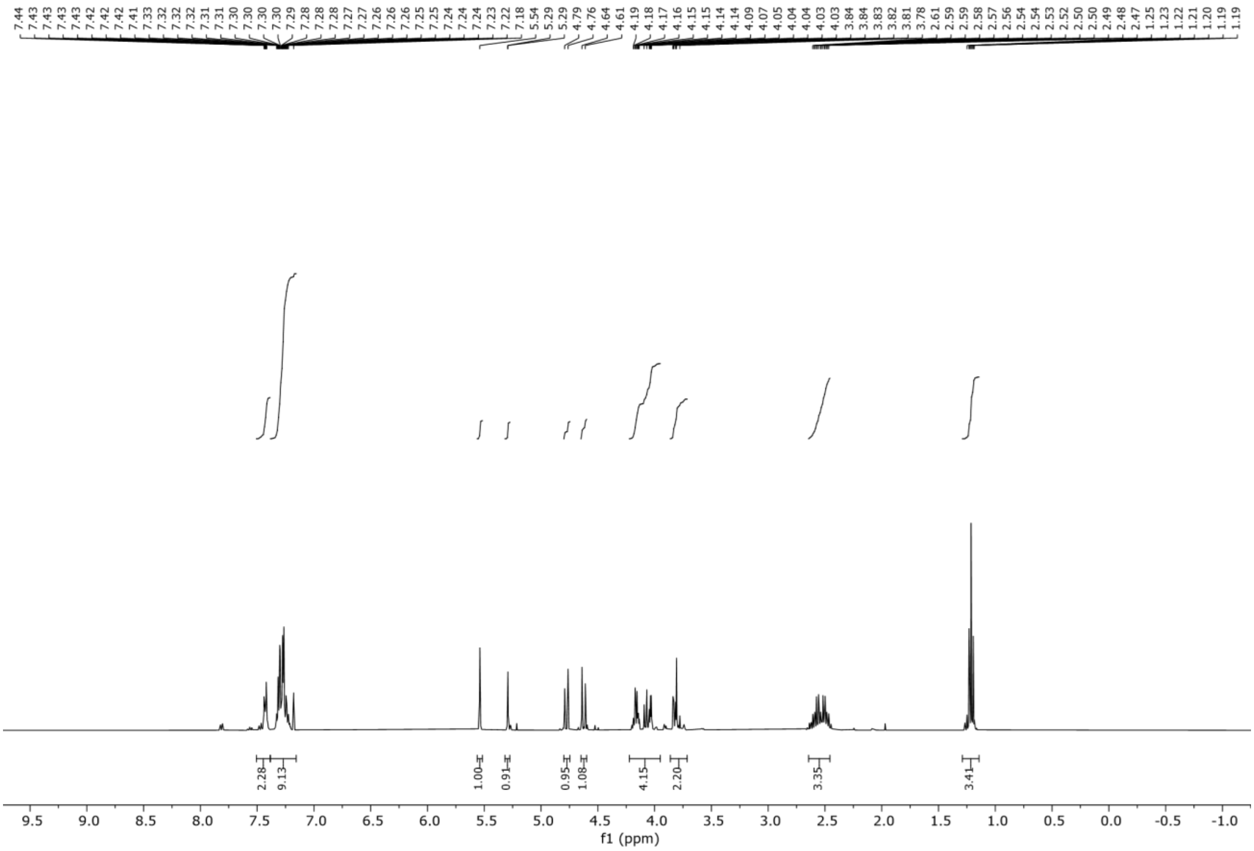


**$^{1}$H NMR**

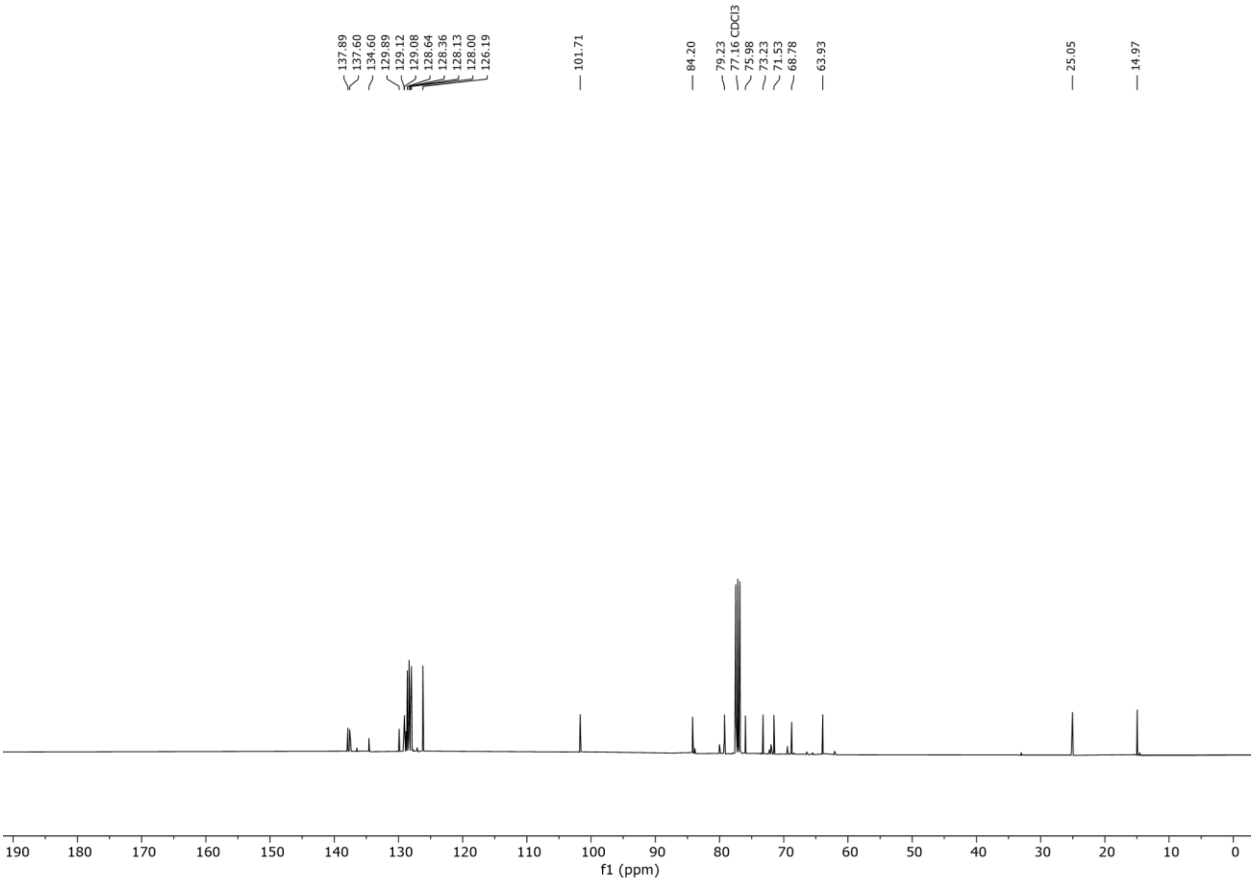


**$^{13}$C NMR**

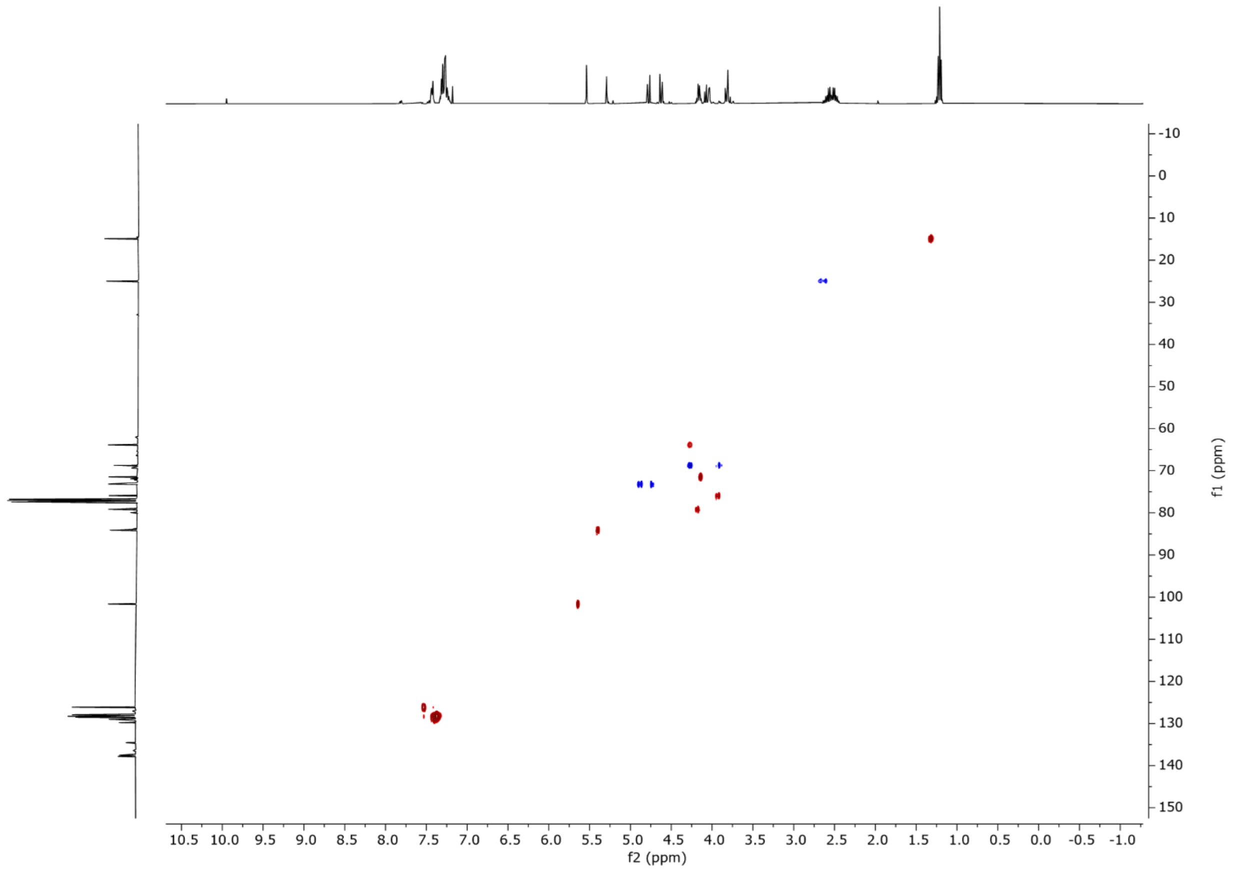


**$^{1}$H-$^{13}$C HSQC NMR**

**2-*O*-sulfonate-D-mannopyranose (12)**

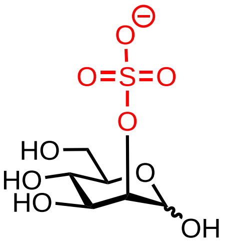


Ethyl 4,6-*O*-benzylidene-3-*O*-benzyl-1-thio-α-D-mannopyranoside (1 g, 2.48 mmol) was dissolved in $CH_2Cl_2$ (0.1 M, 25 mL) at 0 °C. Subsequently, acetic anhydride (0.4 mL, 4.96 mmol, 2 eq.), pyridine (1 mL, 12.4 mmol, 5 eq.) and 4-dimethylaminopyridine (60 mg, 0.49 mmol, 0.2 eq.) were added. The mixture was stirred at room temperature under argon atmosphere. Once complete, as indicated by TLC analysis (3 h), the reaction was quenched with citric acid (10% w/v) and extracted. The organic layer was washed with saturated aqueous $NaHCO_3$, brine, dried over $Na_2SO_4$, and concentrated. The residue was purified by flash chromatography ($SiO_2$, Hexane/Ethyl acetate) to give intermediate ethyl 2-*O*-acetyl-4,6-*O*-benzylidene-3-*O*-benzyl-1-thio-α-D-mannopyranoside. The intermediate was then dissolved in anhydrous $CH_2Cl_2$ (0.1M, 25 mL) and benzyl alcohol (0.7 mL, 7.44 mmol, 3 eq.) and 4Å molecular sieves were added. After 30 minutes, the mixture was cooled to 0 °C and DMTST (1.2 g, 4.96 mmol, 2 eq.) was added. Once TLC analysis indicated the reaction had reached completion it was quenched with triethylamine (3 mL), diluted with DCM, and filtered through a plug of Celite®. The solution was washed with saturated aqueous $NaHCO_3$, brine, and dried over $Na_2SO_4$ and concentrated. The material was purified by flash chromatography ($SiO_2$, Hexane/EtOAc) to yield the intermediate benzyl 2-*O*-acetyl-4,6-*O*-benzylidene-3-*O*-benzyl-1-thio-α-D-mannopyranoside. This intermediate was dissolved in $CH_2Cl_2$ (0.1M, 25 mL) and the pH was raised to pH 8 with a sodium methoxide (approx. 0.2 mL, 0.5M solution). Once complete, the reaction was quenched by the addition of Amberlite™ (Hydrogen foam), filtered, and concentrated under vacuum. Purification by flash chromatography ($SiO_2$, $CH_2Cl_2$/methanol). The benzyl 4,6-*O*-benzylidene-3-*O*-benzyl-1-thio-α-D-mannopyranoside (0.1 g, 0.11 mmol) was dissolved in DMF (1 mL) and Py·$SO_3$ (1 g, mmol, 1.2 eq.) was added and stirred at room temperature. Upon completion of reaction as indicated by TLC (16 h), a premixed solution of $Et_3N$/MeOH (1/1, v/v, 1.0 mL) was added to quench the reaction and stirring was continued for another 30 mins. The reaction mixture was concentrated and purified by flash chromatography ($SiO_2$, MeOH/ $CH_2Cl_2$) to give the intermediate sulfated compound. This intermediate was then dissolved in THF: *t*BuOH: $H_2O$ (4 mL, 50:20:30) and 5% Pd-C (200 mg) was added. The reaction was stirred under $H_2$ atmosphere for 16 h. The reaction was filtered through a pad of celite and washed with *t*BuOH and $H_2O$. The filtrates were concentrated *in vacuum*, and dissolved in 2.0 mL of water for RP-HPLC purification (Method 6) to yield **12** (14 mg, 0.05 mmol, 2%, over 6 steps).

**$^1$H NMR** (400 MHz, $D_2O$) δ 5.39 (s, 1H), 4.47 – 4.43 (m, 1H), 3.91 (dd, *J* = 9.4, 3.9 Hz, 1H), 3.84 – 3.66 (m, 3H), 3.59 (t, *J* = 9.6 Hz, 1H). **$^{13}$C NMR** (101 MHz, $D_2O$) δ 91.6, 77.7, 72.2, 68.7, 66.6, 60.6. **HRMS** QTOF-MS: calcd. $C_6H_{11}O_9S$ for $[M-H]^-$ 259.0129, found 259.0134.

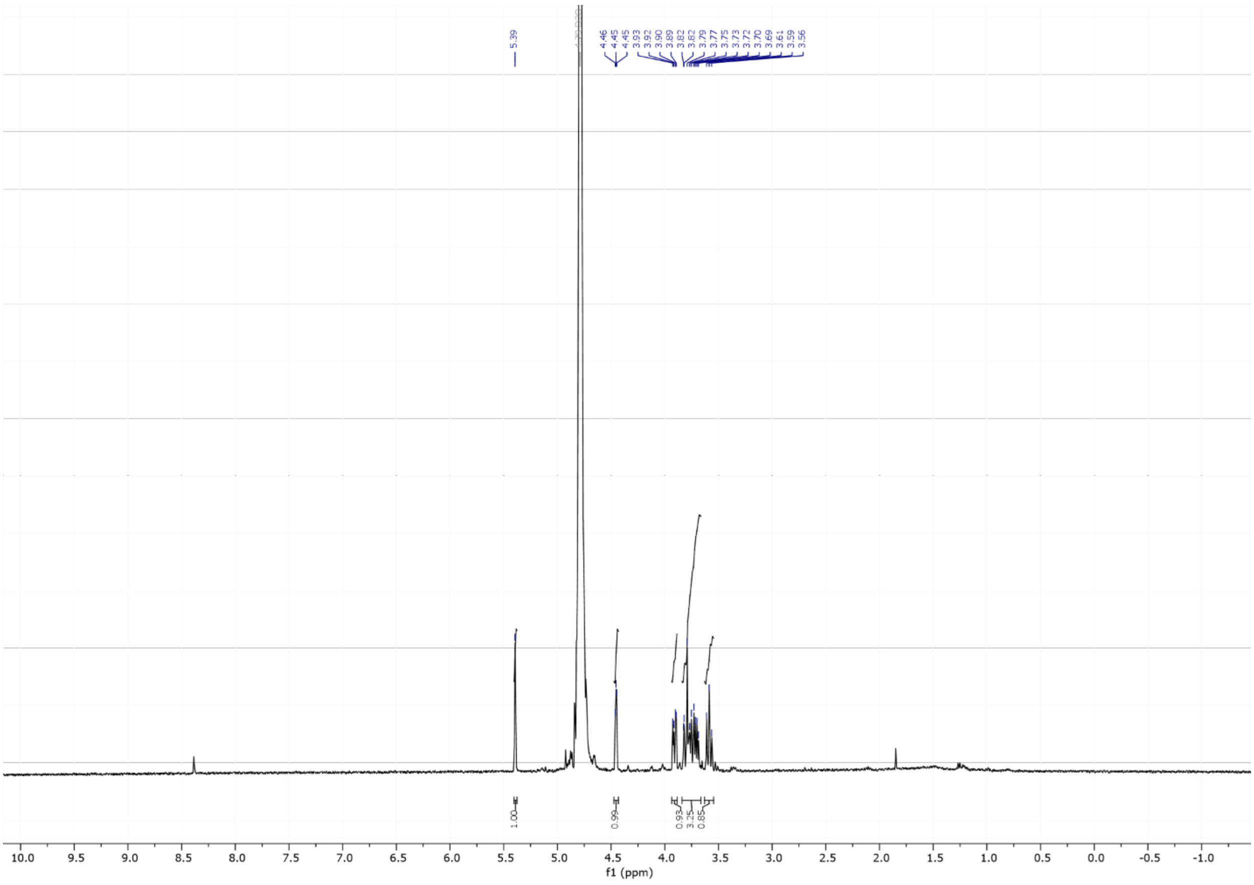


**[1]H NMR**

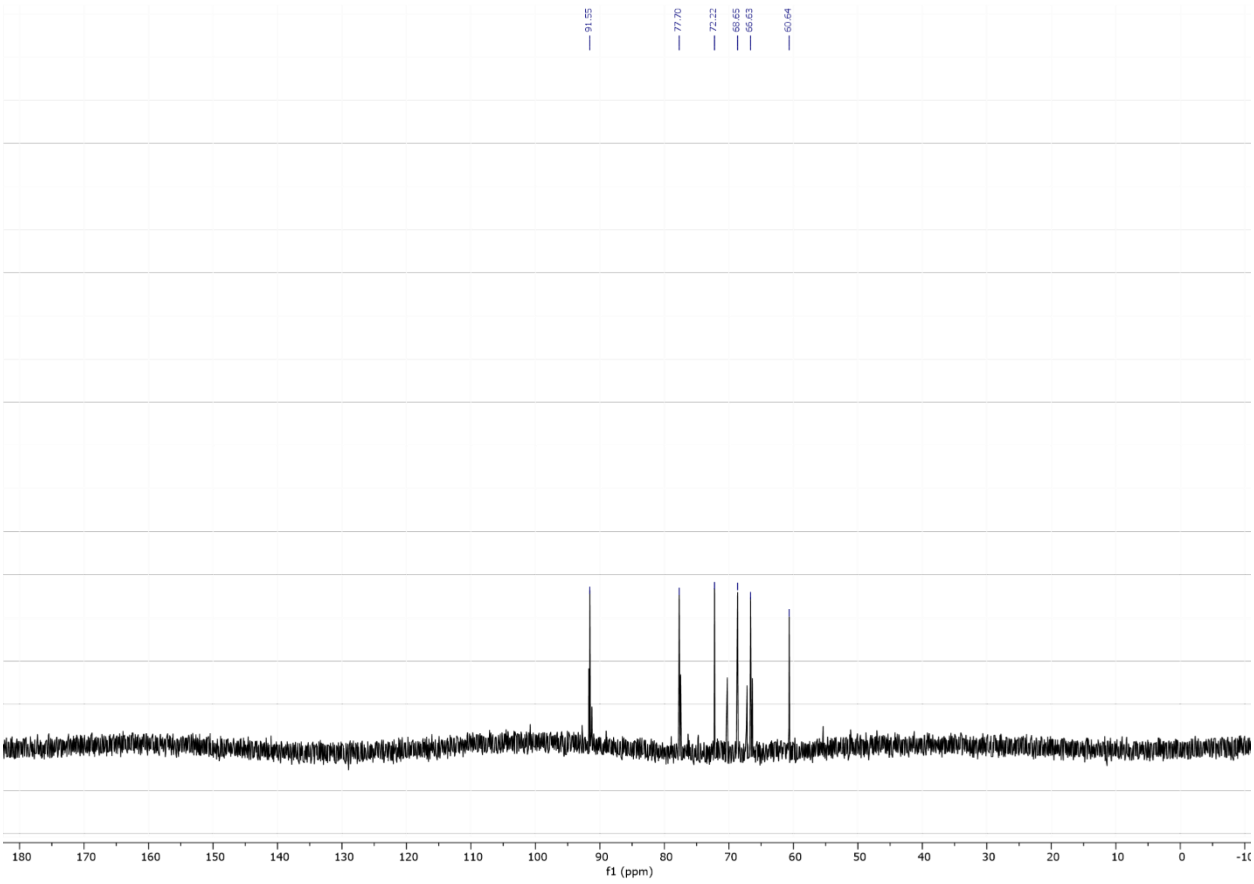


**[13]C NMR**

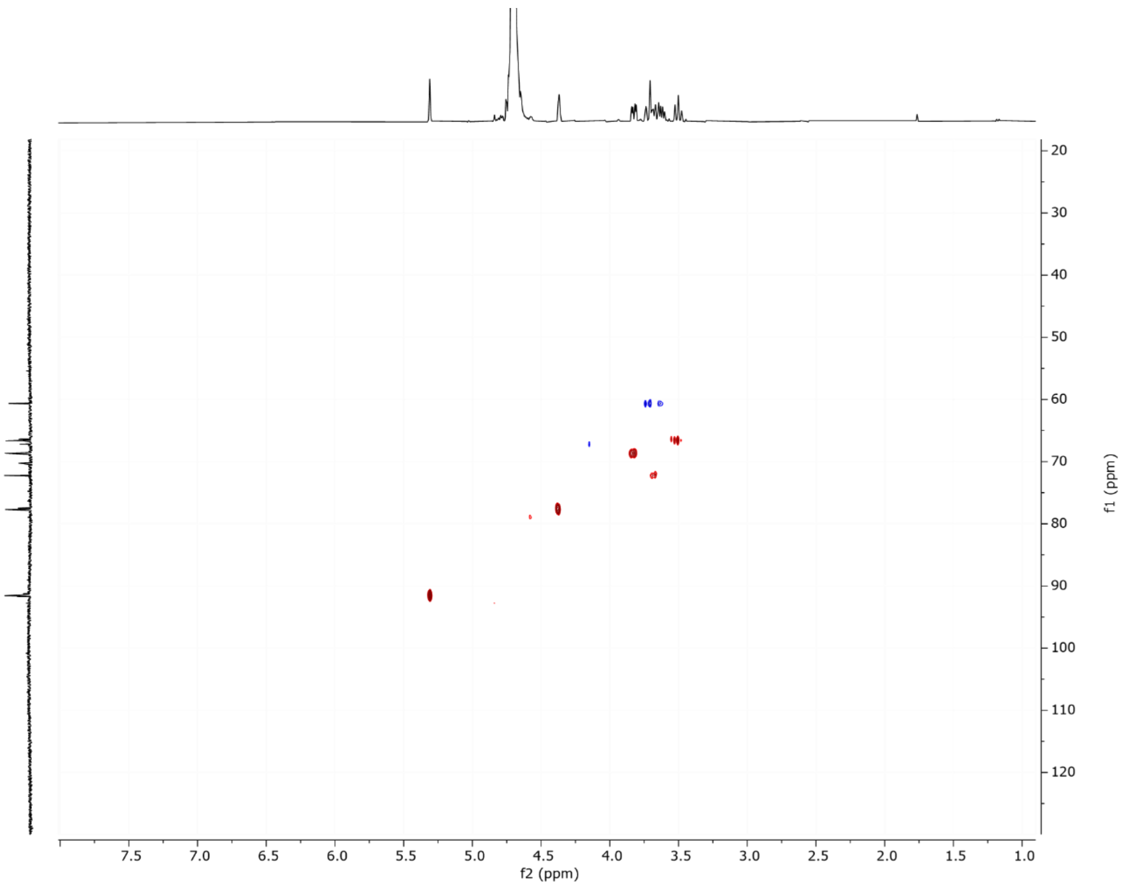
7.5 7.0 6.5 6.0 5.5 5.0 4.5 4.0 3.5 3.0 2.5 2.0 1.5 1.0
f2 (ppm)
20 30 40 50 60 70 80 90 100 110 120
f1 (ppm)

**$^{1}H$-$^{13}C$ HSQC**

**6-*O*-sulfonate-α-D-mannopyranosyl-(1→3)-6-*O*-sulfonate-α-D-mannopyranosyl-(1→3)-6-*O*-sulfonate-D-mannopyranosyl-(1→3)-6-*O*-sulfonate-α-D-mannopyranosyl-(1→3)-6-*O*-sulfonate-α-D- mannopyranosyl-(1→3)-6-*O*-sulfonate-α-D-mannopyranose (2)**

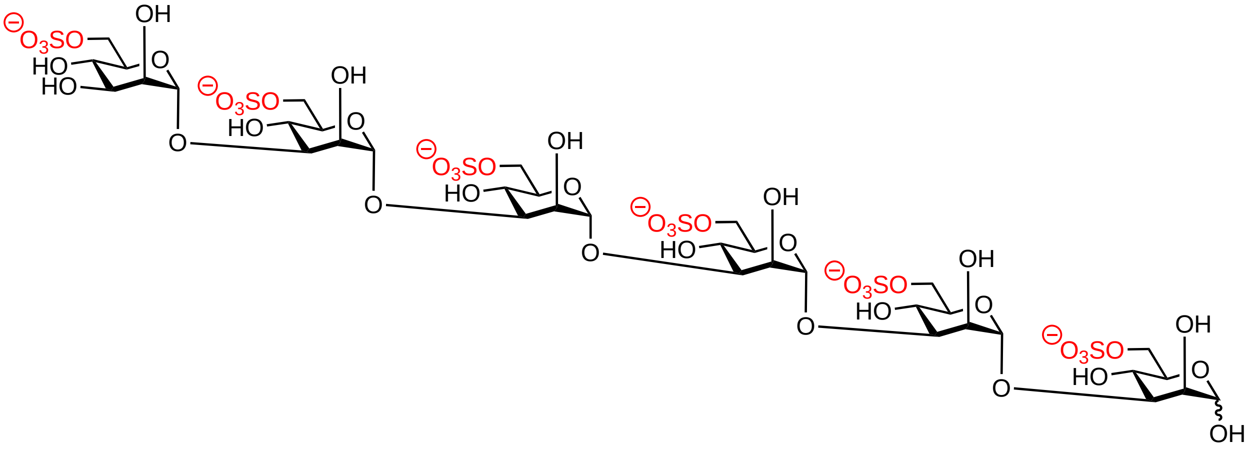

| | | MODULES | NOTES |
|---|---|---|---|
| 1. AGA | | | L2 |
| | **1** | a, b (x2), c, e | x6 |
| | | d | x2 |
| | **Step** | **Module** | **Notes** |
| 2. SOLID-PHASE SYNTHESIS | Sulfation | g | |
| | Methanolysis | f | 72h |
| | photocleavage | h | |
| | **Step** | **Module** | **Notes** |
| 3. SOLUTION-PHASE | Hydrogenolysis | i2 | |
| | Purification | Method 6 | |

The desired fractions were then collected and lyophilized to yield 0.7 mg (2.3%). **$^1$H NMR** (400 MHz, $D_2O$) δ 5.10 (d, *J* = 1.8 Hz, 1H, H-1), 5.08 (d, *J* = 1.7 Hz, 1H, H-1), 5.07 – 5.05 (m, 3H, H-1), 4.89 (s, 0H, H-1), 4.36 – 4.27 (m, 6H, H-6, H-2), 4.26 – 4.13 (m, 6H), 4.11 – 3.94 (m, 5H, H-3), 3.91 – 3.82 (m, 2H, C-3), 3.82 – 3.71 (m, 4H), 3.71 – 3.56 (m, 1H). **$^{13}$C NMR** (101 MHz, $D_2O$) δ 102.6 (C-1), 94.0 (C-1), 79.0 (C-3), 71.3, 69.4, 67.4 (C-6), 65.5. **HRMS** QTOF-MS: calcd. $C_{36}H_{59}O_{49}S_6$ for $[M-3H]^{-3}$ 489.0155, found 489.0159.

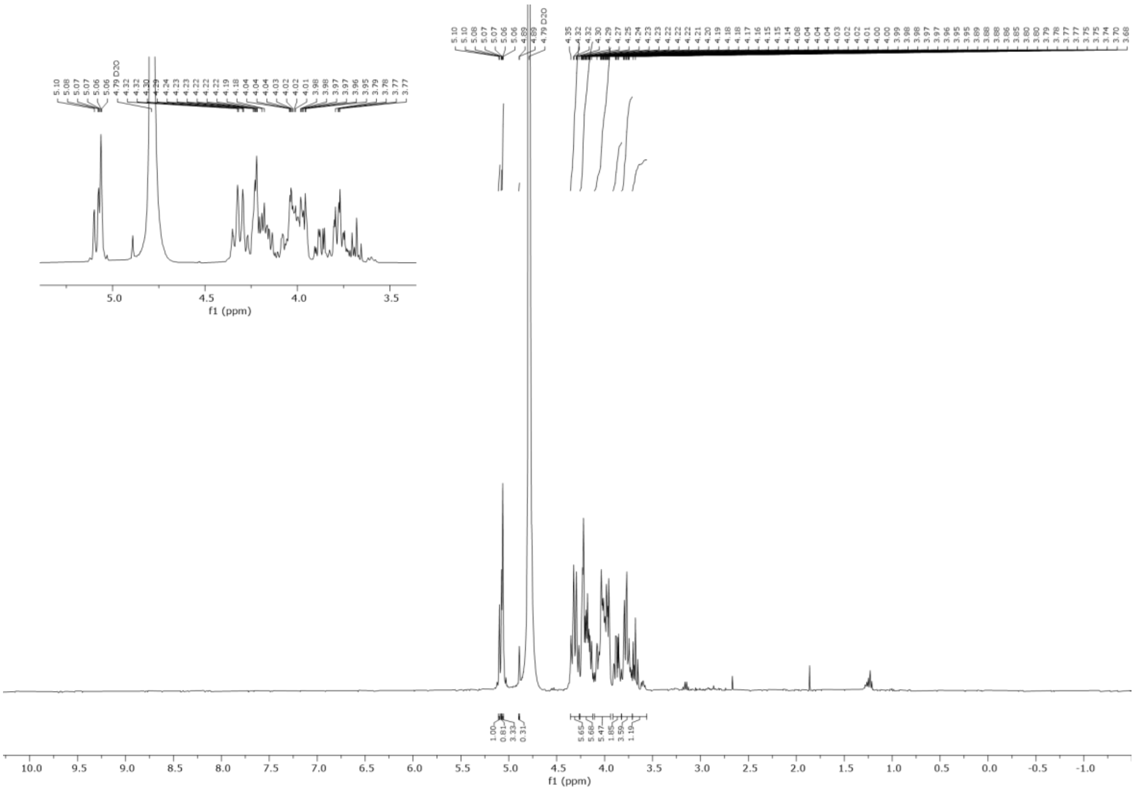


**$^1H$ NMR**

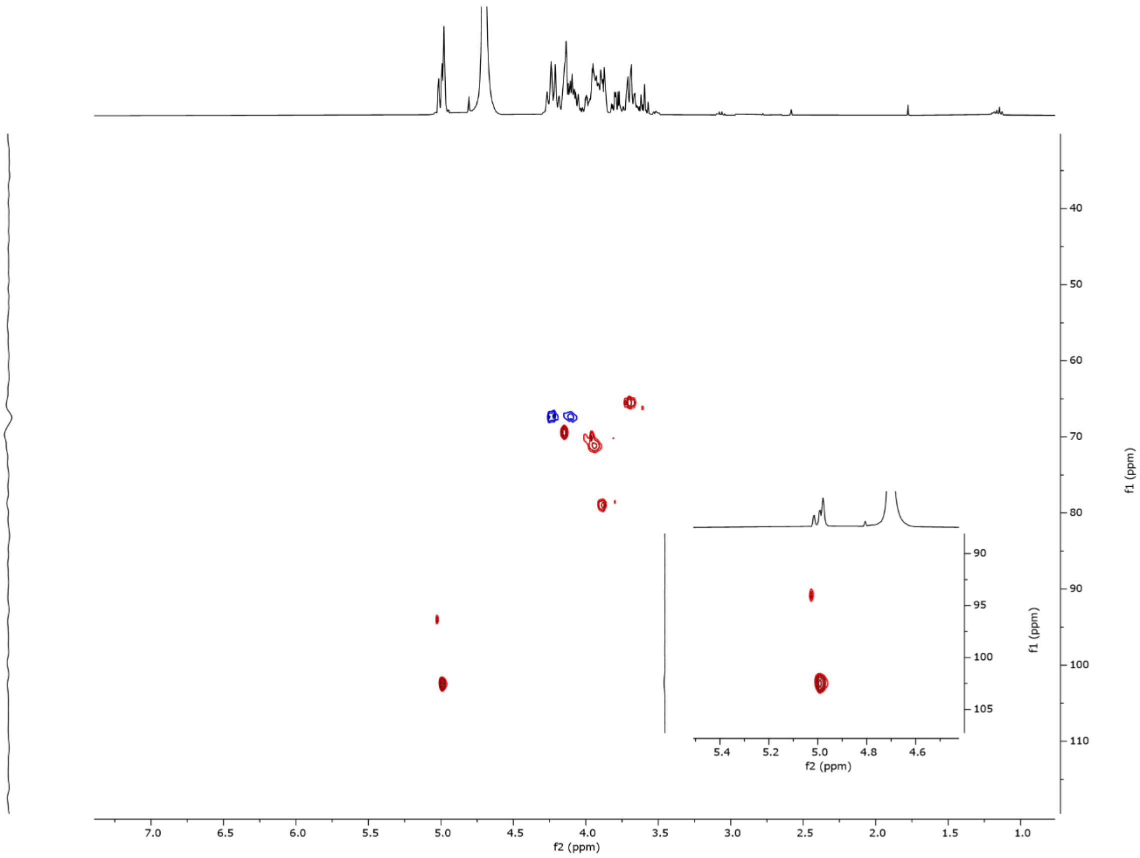


**$^1H$-$^{13}C$ HSQC NMR**

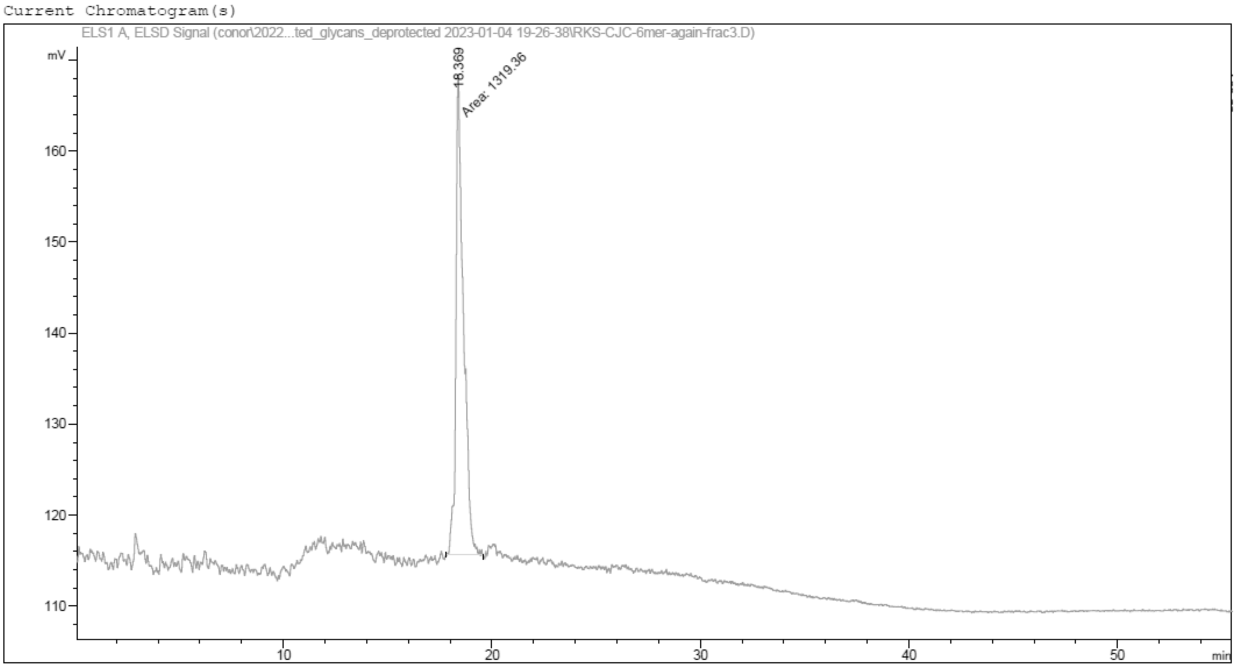


**RP HPLC Trace.** Hypercarb**.**

**α-D-mannopyranosyl-(1→3)-α-D-mannopyranosyl-(1→3)-D-L- mannopyranosyl-(1→3)-α-D-mannopyranosyl-(1→3)-α-D- mannopyranosyl-(1→3)-α-D-mannopyranose (3)**

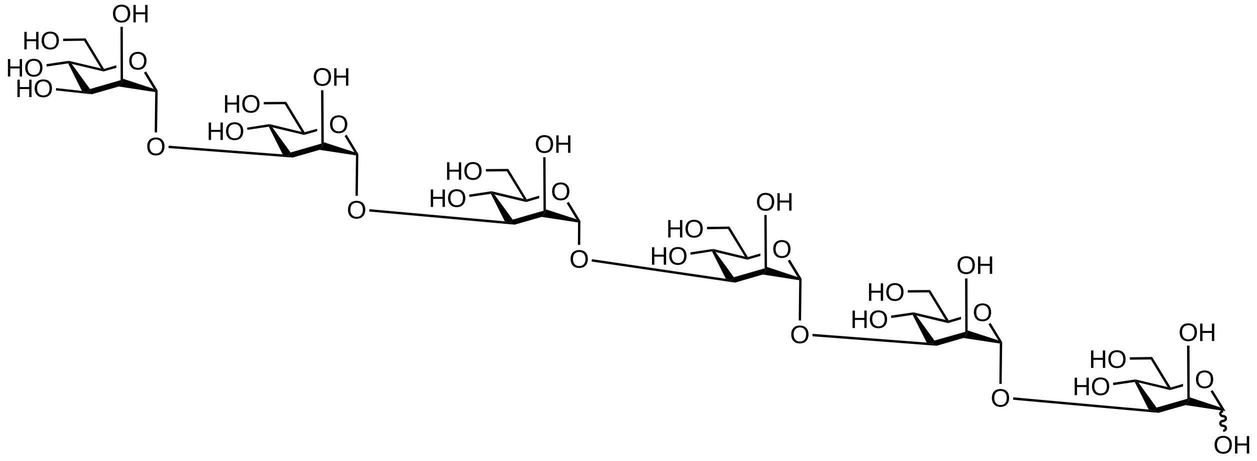

| | | MODULES | NOTES |
|---|---|---|---|
| 1. AGA | | | L2 |
| | **1** | a, b (x2), c, e | x6 |
| | | d | x2 |
| | **Step** | **Module** | **Notes** |
| 2. SOLID-PHASE SYNTHESIS | Methanolysis | f | |
| | photocleavage | h | |
| | **Step** | **Module** | **Notes** |
| 3. SOLUTION-PHASE | Hydrogenolysis | i1 | |
| | Purification | Method 5 | |

The desired fractions were then collected and lyophilized to yield 0.9 mg (4.5%). **$^{1}$H NMR** (600 MHz, $D_2O$) δ 5.15 (d, *J* = 1.9 Hz, 1H, H-1), 5.14 (d, *J* = 1.9 Hz, 1H, H-1), 5.13 – 5.11 (m, 5H, H-1), 4.91 (sf, 0H), 4.26 – 4.22 (m, 6H, H-2), 4.10 – 4.05 (m, 4H), 4.05 – 4.00 (m, 6H, H-3), 3.95 – 3.87 (m, 12H), 3.87 – 3.81 (m, 8H), 3.81 – 3.72 (m, 19H), 3.66 (m, 7H). **$^{13}$C NMR** (151 MHz, $D_2O$) δ 102.2 (C-1), 93.9 (C-1), 78.2 (C-3), 73.5, 70.3, 70.0, 69.7 (C-2), 66.1, 61.0 (C-6). **HRMS** QTOF-MS: calcd. $C_{36}H_{62}NaO_{31}$ for $[M+Na]^{-}$ 1013.3173, found 1013.3179.

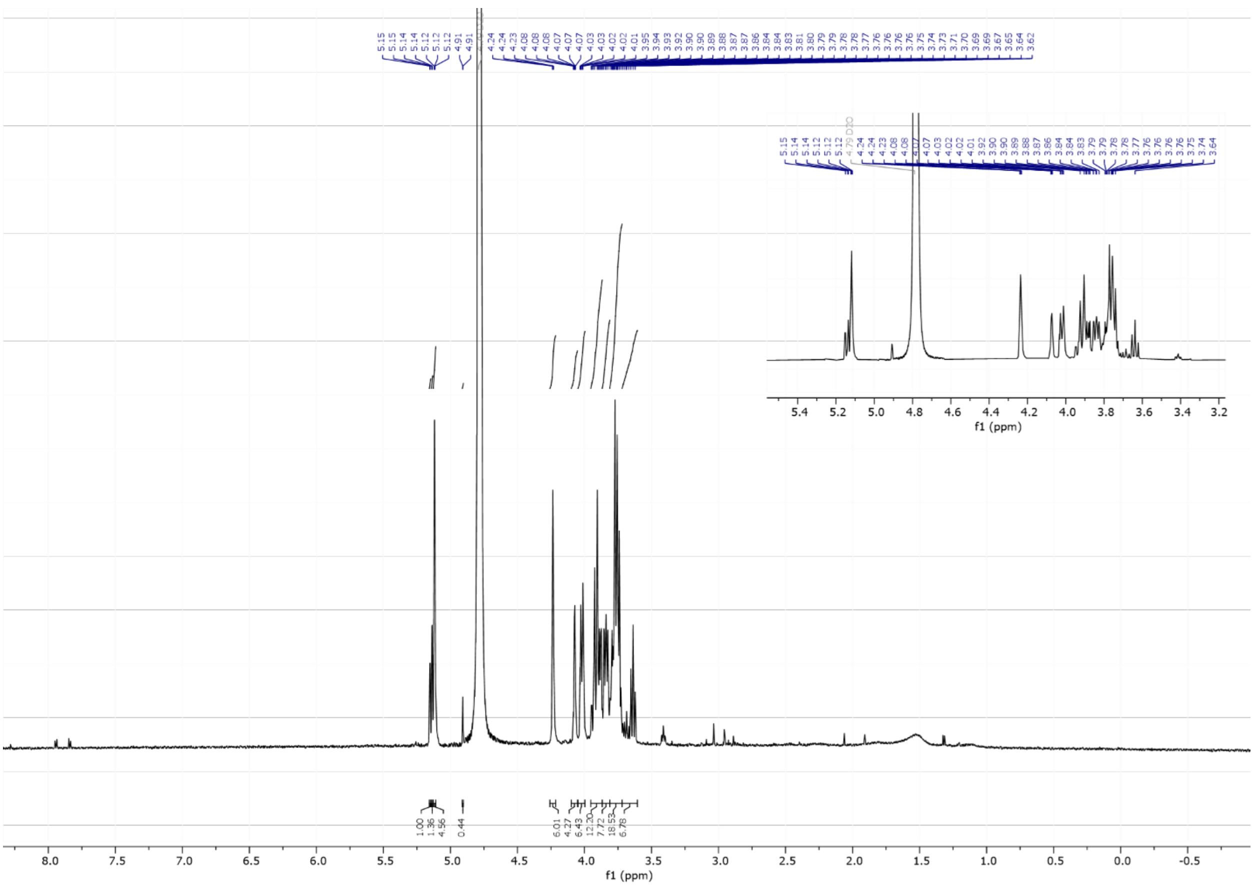

8.0 7.5 7.0 6.5 6.0 5.5 5.0 4.5 4.0 3.5 3.0 2.5 2.0 1.5 1.0 0.5 0.0 -0.5
f1 (ppm)
5.4 5.2 5.0 4.8 4.6 4.4 4.2 4.0 3.8 3.6 3.4 3.2
f1 (ppm)


**$^{1}$H NMR**

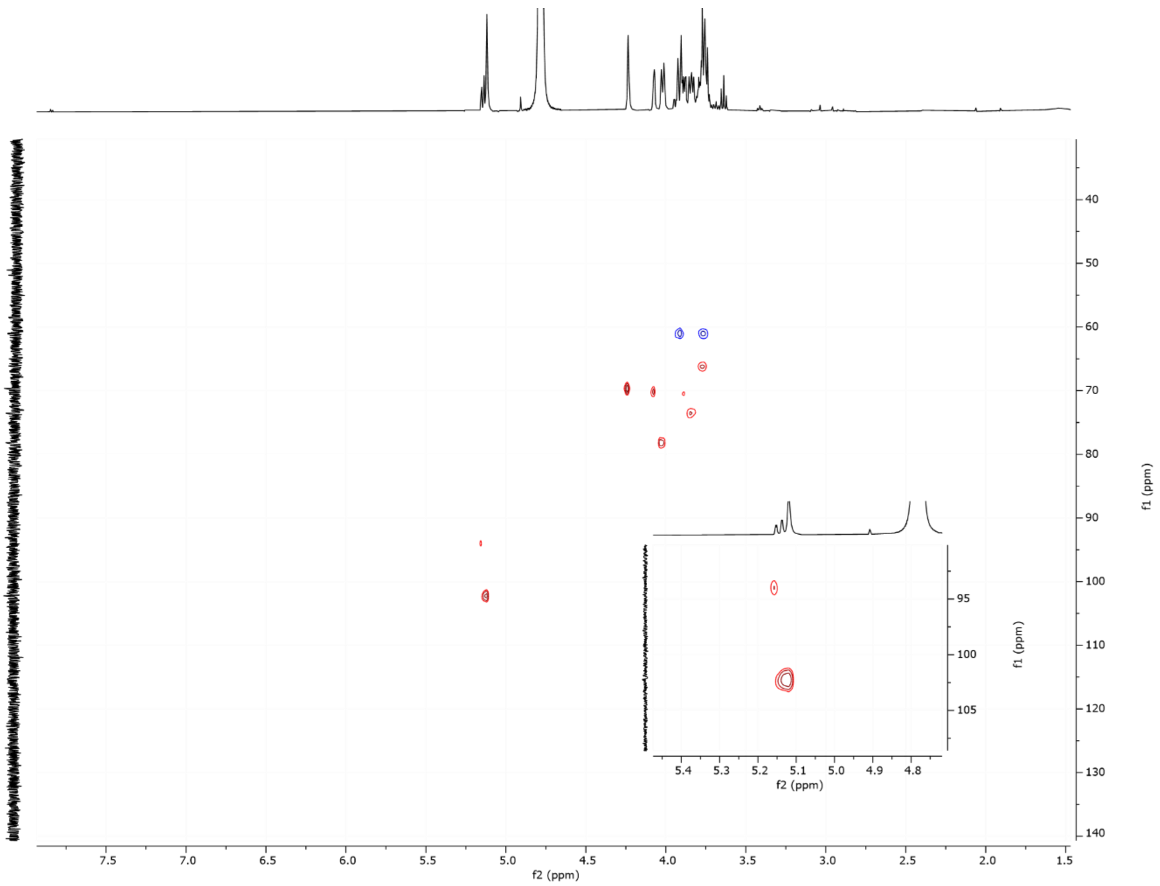


**$^{1}H$-$^{13}C$ HSQC**

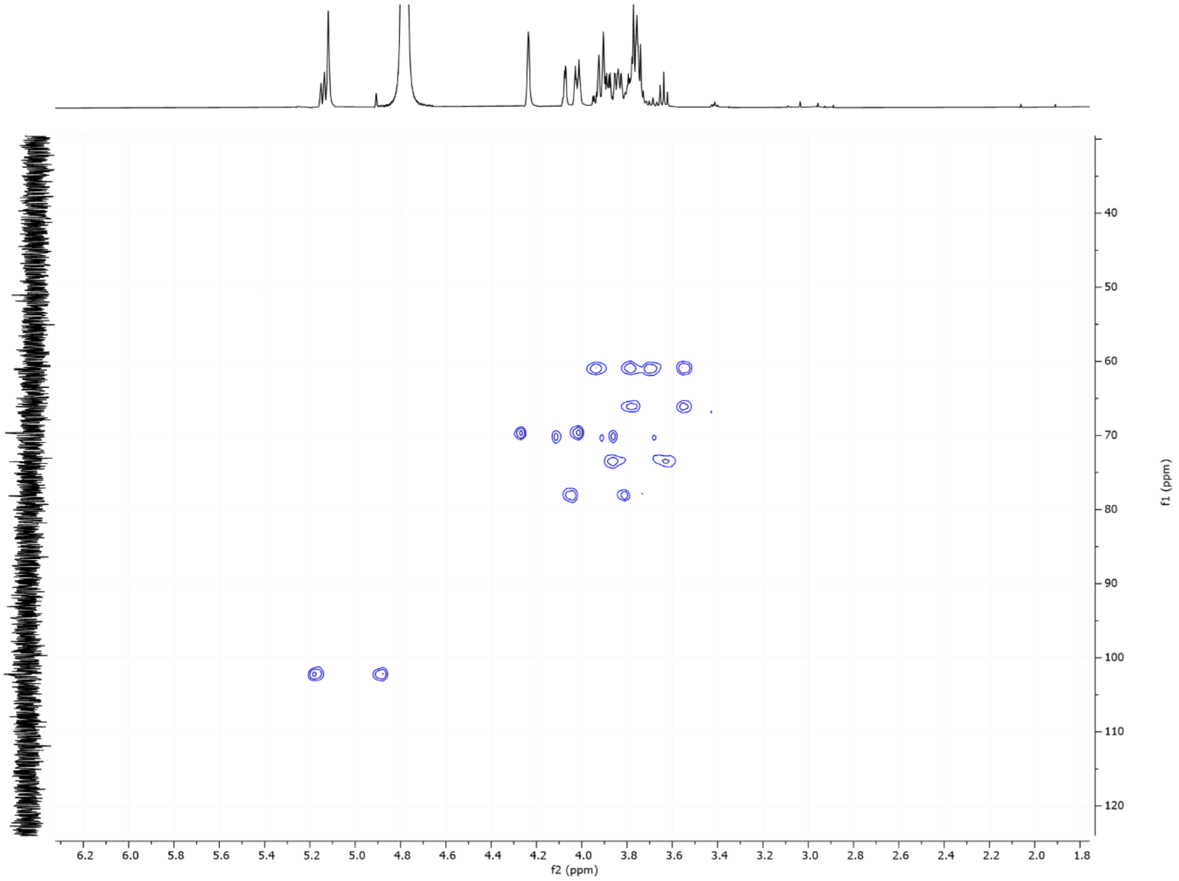


**$^{1}H$-$^{13}C$ HSCQC (coupled)**

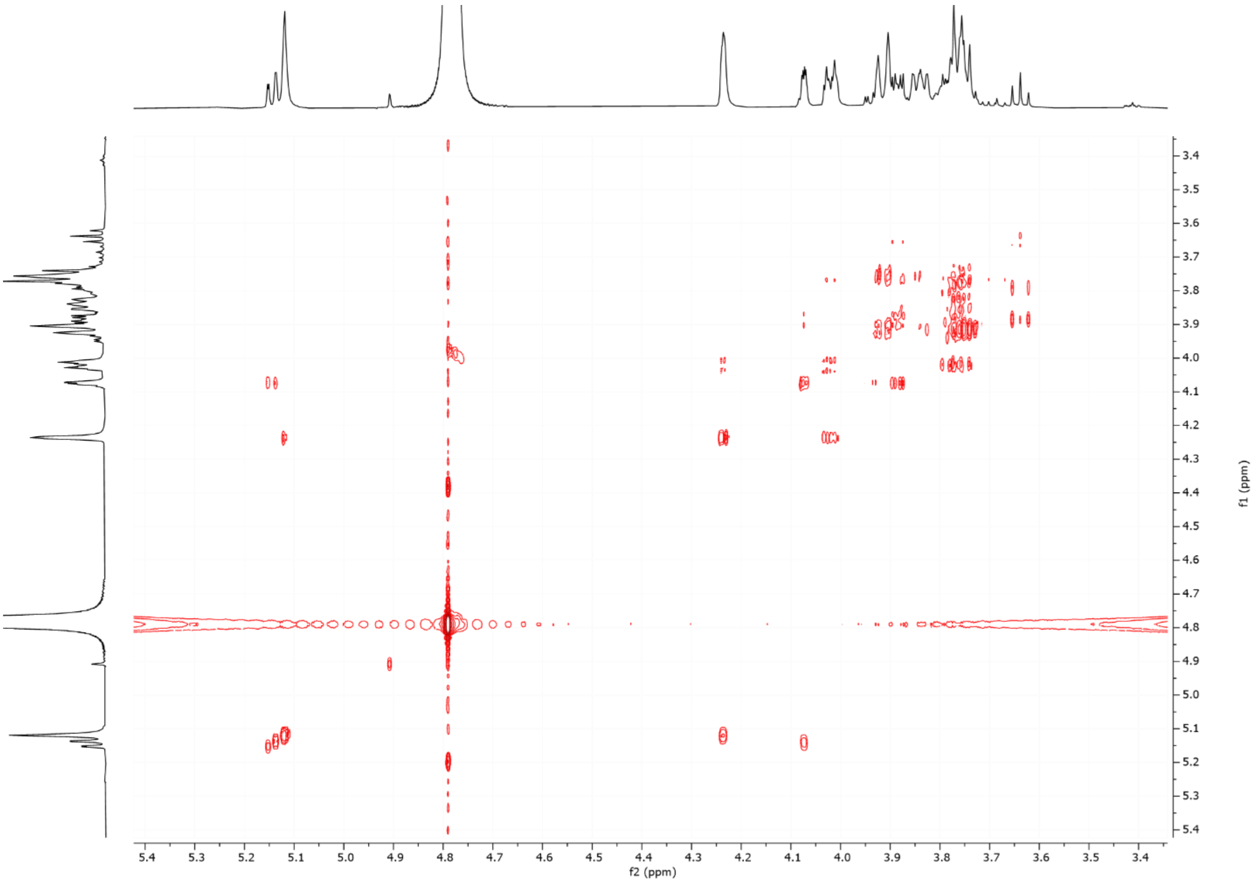


$^{1}H$-$^{1}H$ **COSY NMR**

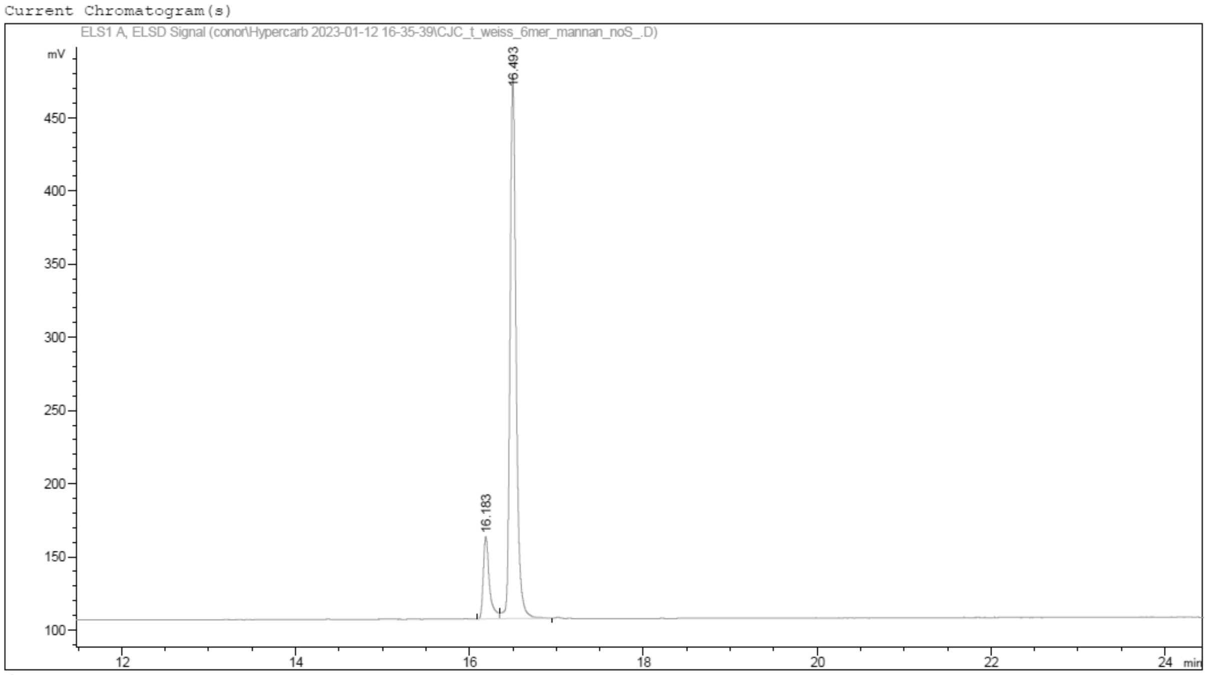


**RP HPLC Trace.** Hypercarb.
